# Prioritized single-cell proteomics reveals molecular and functional polarization across primary macrophages

**DOI:** 10.1101/2022.03.16.484655

**Authors:** R Gray Huffman, Andrew Leduc, Christoph Wichmann, Marco di Gioia, Francesco Borriello, Harrison Specht, Jason Derks, Saad Khan, Luke Khoury, Edward Emmott, Aleksandra A. Petelski, David H Perlman, Jürgen Cox, Ivan Zanoni, Nikolai Slavov

## Abstract

Major aims of single-cell proteomics include increasing the consistency, sensitivity, and depth of protein quantification, especially for proteins and modifications of biological interest. To simultaneously advance all these aims, we developed prioritized Single Cell ProtEomics (pSCoPE). pSCoPE consistently analyzes thousands of prioritized peptides across all single cells (thus increasing data completeness) while analyzing identifiable peptides at full duty-cycle, thus increasing proteome depth. These strategies increased the sensitivity, data completeness, and proteome coverage over 2-fold. The gains enabled quantifying protein variation in untreated and lipopolysaccharide-treated primary macrophages. Within each condition, proteins covaried within functional sets, including phagosome maturation and proton transport. This protein covariation within a treatment condition was similar across the treatment conditions and coupled to phenotypic variability in endocytic activity. pSCoPE also enabled quantifying proteolytic products, suggesting a gradient of cathepsin activities within a treatment condition. pSCoPE is freely available and widely applicable, especially for analyzing proteins of interest without sacrificing proteome coverage. Support for pSCoPE is available at: scp.slavovlab.net/pSCoPE

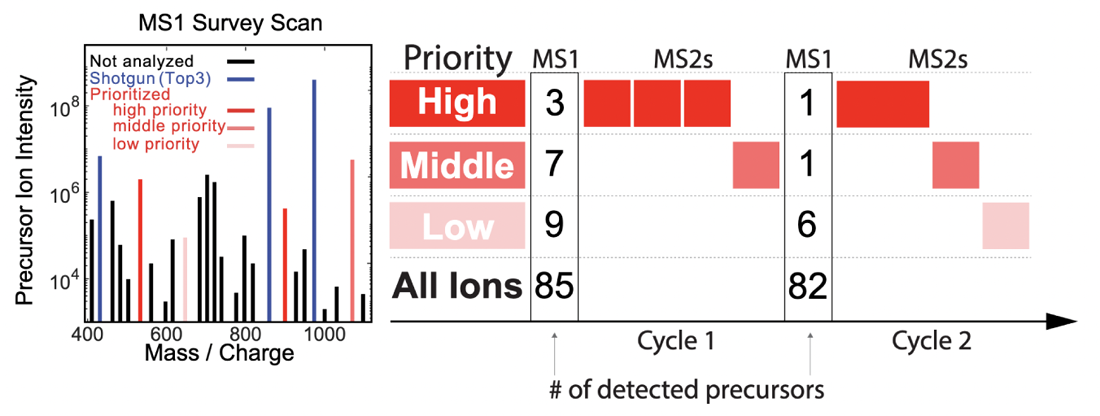

## Introduction

Macrophages are innate immune myeloid cells performing diverse functions in development, tissue homeostasis and immune responses. Despite this diversity, macrophages are traditionally described in terms of dichotomous states (M1, pro-inflammatory; M2, anti-inflammatory). Single-cell measurements, though, have revealed a more complex and continuous spectrum of macrophage polarization in terms of molecular and functional phenotypes^1–3^. Thus, we sought to explore this continuum of polarized states in primary macrophages using single-cell mass-spectrometry (MS). Shotgun MS methods can analyze hundreds of single cells per day and quantify thousands of proteins but remain biased towards abundant proteins^3–11^. This bias reflects an intentionally programmed ‘topN’ heuristic for selecting the *N* most abundant peptide precursors for sequence identification and quantification^12^, as illustrated in Fig. 1a.

**Figure 1.**
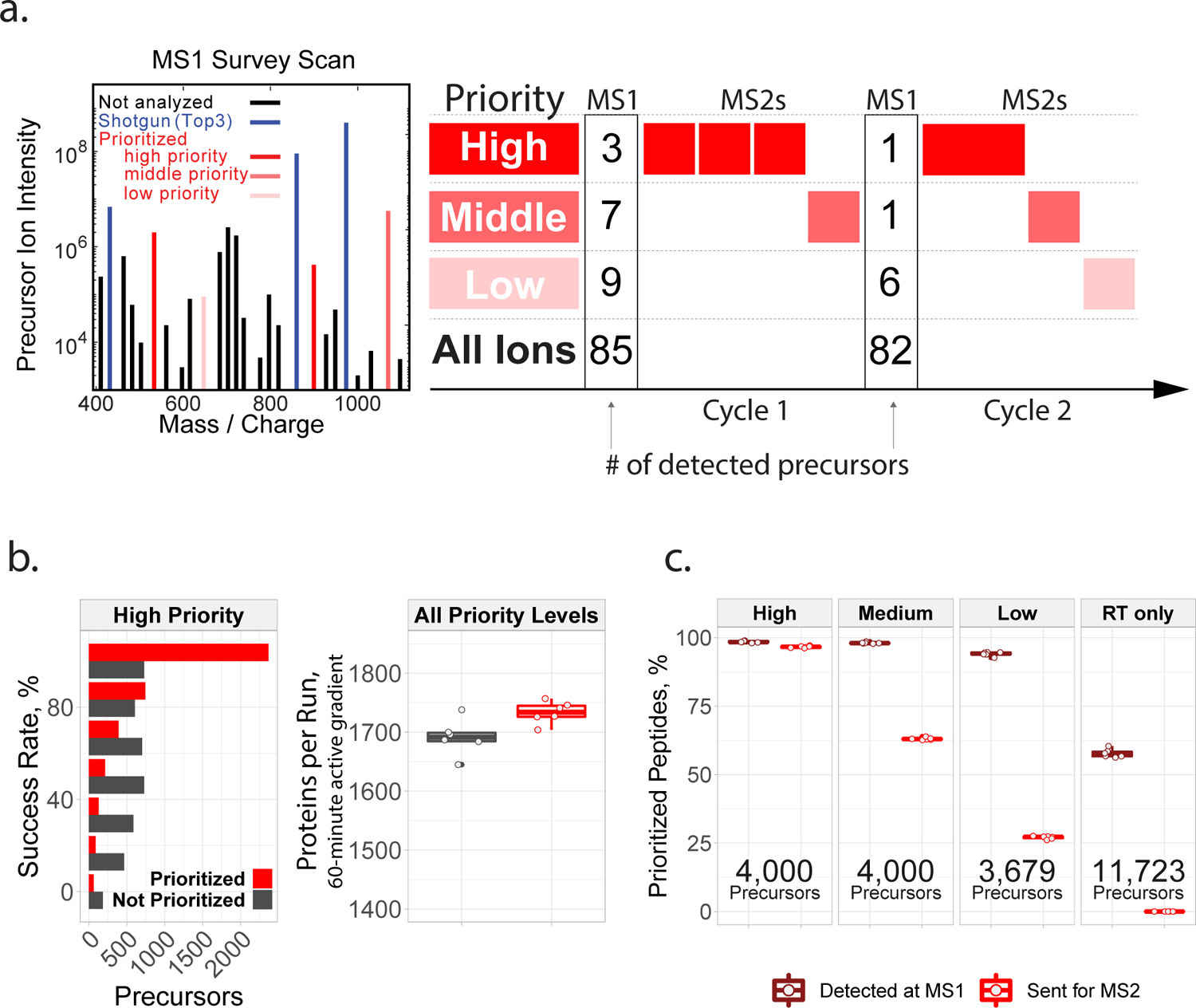
Introducing prioritization to MQ.Live increases identification consistency and protein coverage. (a) Shotgun TopN analysis selects the *N* most abundant precursors for isolation and fragmentation (shown in blue). Among the many detected precursors, prioritized analysis first selects the ones with highest priority (shown in solid red) and then from lower priority tiers (shown with decreasingly saturated red tones). (**b**) Prioritized analysis increases the consistency of peptide identification over default MaxQuant.Live operation for High Priority peptides while also increasing protein coverage per run. The box plots showing proteins identified per experiment containing six points per analysis method, one for each experiment. (**c**) Rates of MS1 detection and MS2 analysis for prioritized precursors from all tiers of the benchmarking experiments displayed in panel b.

Peptide selection by the topN heuristic is limited by three challenges: (i) abundance bias, which limits the dynamic range of quantified proteins; (ii) stochasticity, which limits data completeness across single cells; and (iii) unidentifiable precursors, whose analysis wastes instrument time and limits proteome coverage^13^. Such inefficient use of time is particularly limiting for single-cell proteomics due to the long ion accumulation times needed to sequence and quantify each precursor^3, 14^. While no existing method resolves all 3 challenges, the challenges can be partially mitigated. For example, targeted MS can alleviate challenges (i) and (ii) but has remained limited to analyzing merely hundreds of peptides or fewer^15–23^. Real-time database searching can increase the fraction of sequenced peptide features and alleviate challenge (iii), but it has not allowed for selecting peptides of interest^7, 24^. Targeting peptides from inclusion lists with real-time retention-time alignment ameliorates challenge (i) but faces a trade-off between maximizing coverage (and thus duty cycle usage) and maximizing data completeness^25, 26^. To simultaneously address all three challenges, we introduce multitiered prioritized selection of peptide precursors using the strategy sketched in Fig. 1a.

## Results

Prioritized analysis aims to simultaneously maximize the consistency of peptide analysis, proteome coverage, and instrument time utilization. To achieve these aims, we built upon the real-time retention-time alignment of MaxQuant.Live^26, 27^ and introduced priority levels that define the temporal order of peptide analysis. High priority aim to maximize data completeness when the duty-cycle time is insufficient for analyzing all peptides from the inclusion list whose precursors are detected in survey scans. Two example duty cycles implementing this selection logic are displayed in Fig. 1a.

### Increasing proteome coverage and data completeness

The logic of prioritized peptide acquisition is implemented via new functionality in MaxQuant.Live software that seeks to maximize both data completeness and proteome coverage, Fig. 1a. To maximize proteome coverage, a large inclusion list of previously-identified precursors and real-time alignment allow filling each and every duty cycle with peptide-like features most likely to be identified. To simultaneously maximize data completeness, the sets of high-priority precursors are supplied and always given priority for MS2 analysis over precursors from lower priority tiers. The sets of high-priority precursors can be selected based on biological interest, ease of identification, spectral purity or other relevant metrics. Increased accumulation times can be allocated for them, such as for the top tier peptides in the second duty cycle in Fig. 1a. This increased accumulation time should increase the number of ion copies samples per MS2 analysis^28, 29^.

To benchmark the benefits of prioritization, MaxQuant.Live was used to acquire data with and without prioritization enabled while keeping all other parameters constant, Fig. 1b. To reduce sample related variability, we analyzed injections from a bulk sample diluted to single-cell levels. The inclusion list was composed of the the same precursors for the prioritized and non-prioritized analysis by MQ.Live: over 11,500 precursors selected to be identifiable, along with a comparable number of precursors used only for retention-time calibration. The precursors on the inclusion list were then stratified into three levels of priority by the confidence of their identification and spectral purity in previous analysis. More confidently-identified and less coisolated peptides were assigned to the higher priority levels. Data completeness for the high-priority group of 4,000 peptides increased to 72% when using prioritization, compared to 49% without prioritization. The fraction of peptides identified in 100% of the 6 runs at 1% FDR was 18% without prioritization and 59% with prioritization (Fig. 1b), representing a 228% increased consistency for prioritization. This increased consistency of identification did not impede protein coverage, as prioritization increased the number of quantified proteins per experiment at 1% FDR, Fig. 1b. Consistent with the prioritization logic shown in Fig. 1a, prioritization sent precursors to MS2 scans according to priority: 97% of the 4000 high-priority peptides were sent for MS2 analysis, and lower fractions of the lower priority tiers, Fig. 1c. In contrast, MaxQuant.Live without prioritization sent similar fractions (about 63%) of peptides from all lists for MS2 analysis as shown in Extended Data Fig. 1b.

We then applied prioritized Single-Cell ProtEomics (pSCoPE) to single human cells to evaluate the depth and the consistency of proteome coverage relative to control shotgun analysis, Fig. 2. We prepared single-cell samples from Embryonic Kidney 293 (HEK) and Melanoma cells using nPOP^30^. These samples were analyzed either by shotgun or by prioritized methods using 60-min active chromatographic gradients and narrow (0.5 Th) isolation windows for MS2 scans. The narrow isolation window resulted in good quantitative agreement between different peptides originating from the same protein, Extended Data Fig. 2a,b. To maximize coverage, we prioritized peptides that were previously identified with high confidence and low coisolation; more details can be found in the Methods section. Relative to shotgun analysis, pSCoPE increased the fraction of MS2 spectra assigned to a confident peptide sequence by over 2-fold, reaching 84%, Fig. 2a. The remaining 16% of MS2 spectra correspond to sequences having lower confidence of identification in previous experiments used for generating the inclusion list, Extended Data Fig. 3. The increase in productive MS2 scans doubled the number of unique peptides per run (increased by 103%) and increased the number of quantified proteins per single cell by 106%, Fig. 2a.

**Figure 2.**
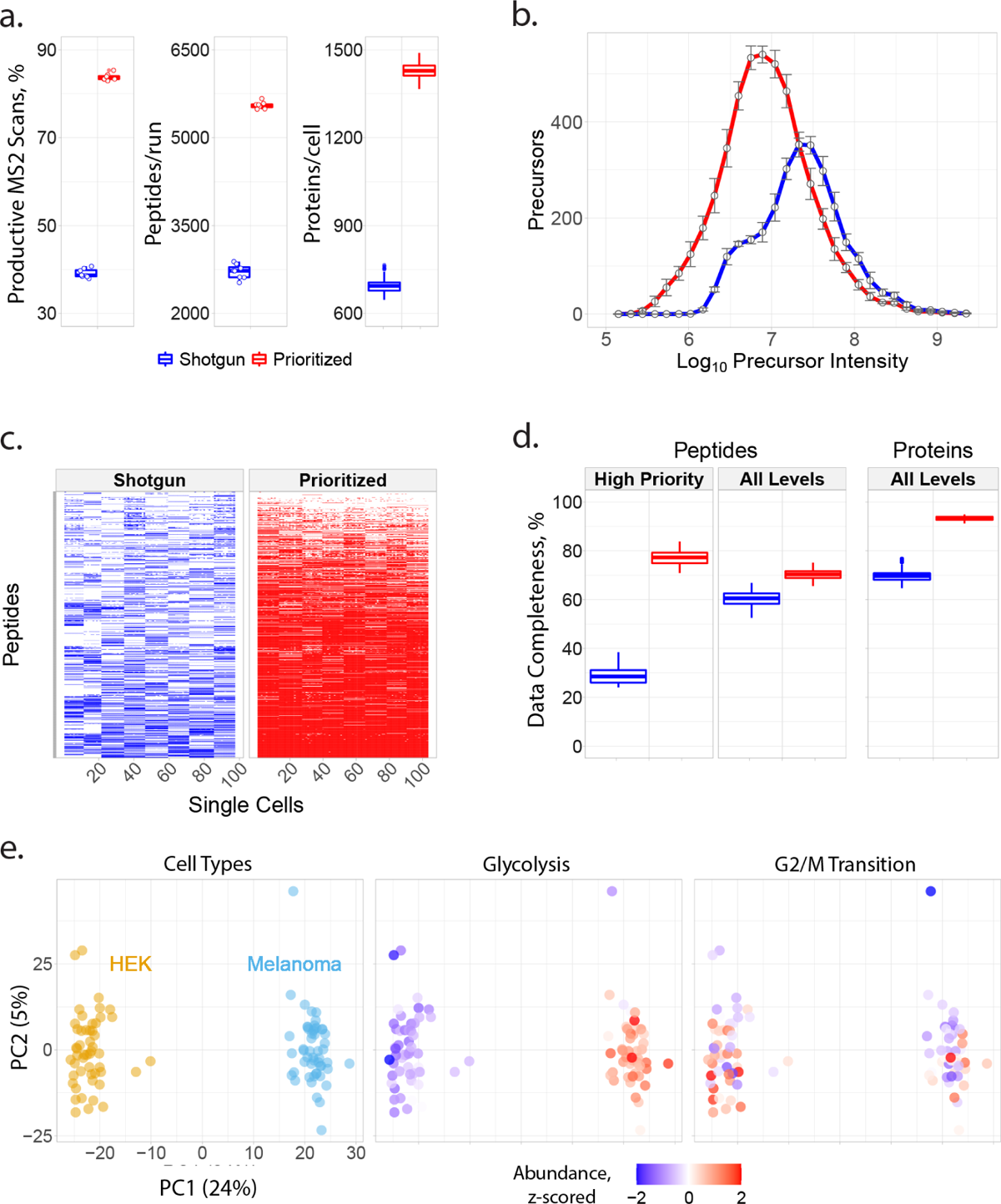
Prioritization increases proteome coverage, sensitivity, and data completeness of single-cell protein analysis. (**a**) Relative to shotgun analysis, prioritization increased the fraction of MS2 scans assigned to peptide sequences, the number of peptides per run (60-min active gradient) and the number of quantified proteins per single cell. (**b**) Prioritized analysis enables increased sensitivity and dynamic range while analyzing more peptides that shotgun analysis with matched parameters. (**c**) A heatmap showing data completeness across single cells (columns) for 1,000 peptides (rows) from the top priority tier. (**d**) Prioritized analysis increases data completeness at both peptide and protein levels across all priority tiers. (**e**) Principal component analysis of the single cells associated with (b) cluster by cell type. Protein sets enriched in the PCs are visualized by color-coding the single cells by the median protein abundance of the set in each cell. All experiments used 60 min active gradients per run and 0.5 Th isolation windows for MS2 scans. All peptide and protein identifications were filtered at 1% FDR with additional filtration criteria detailed in Methods.

**Figure 3.**
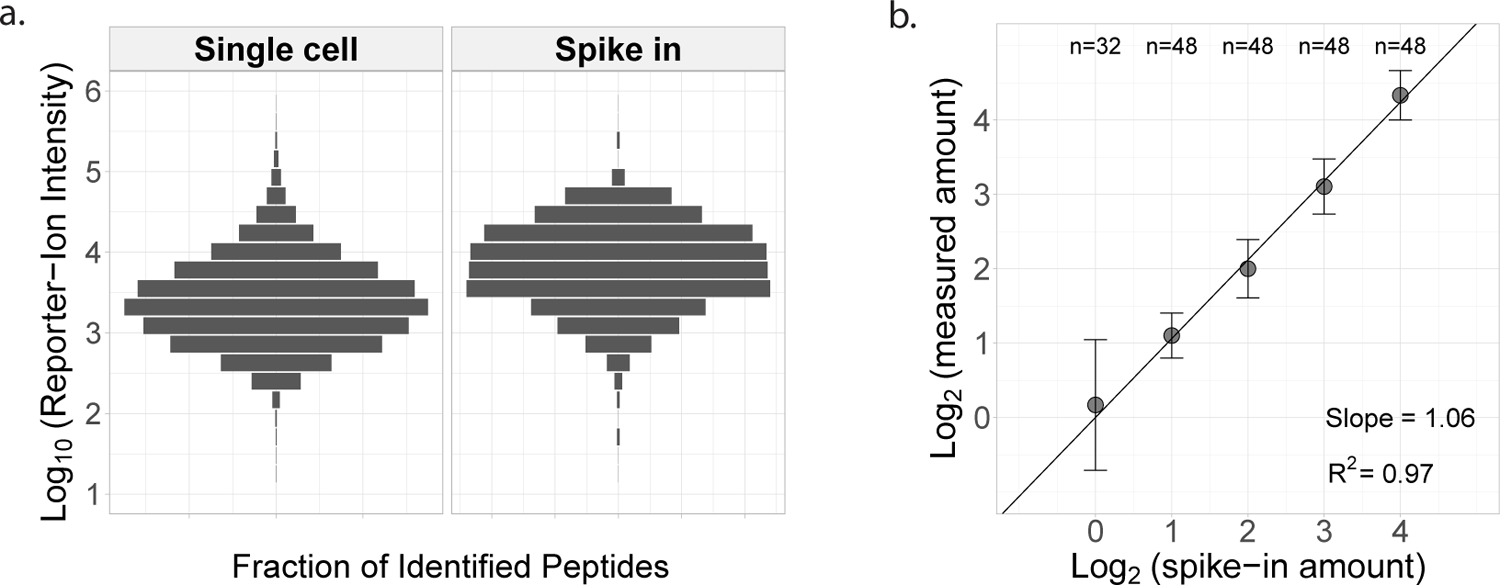
Evaluating quantitative accuracy and precision of pSCoPE with peptide standards. (**a**) Reporter ion intensities for precursors identified at 1% FDR from single cells and from the spike-in peptides, which were dispensed into the single-cell samples across a 16-fold range. (**b**) Normalized reporter ion intensities for all tryptic products from the spike-in peptides plotted against their spike-in amounts, with regression slope and goodness of fit displayed. The data in both panels comes from 8 prioritized experiments.

In addition to increasing the depth of proteome coverage, the prioritized approach increased the dynamic range of proteins quantified in single cells, Fig. 2b. This is enabled by the high-probability with which pSCoPE sends prioritized precursors for MS2 analysis. Specifically, single-cell analyses with pSCoPE results in MS2 scans for 98% of the peptides on a 1,000-peptide high-priority level, 90% of the 4,475 peptides from the top two priority levels, and 71% of the 8,621 peptides on the top three priority levels (Extended Data Fig. 5). These efficiencies were achieved while pSCoPE operated at full duty cycle, demonstrating the ability of tiers (prioritization) to increase the probability of analyzing peptides while running at full duty cycle. This ability to successfully prioritize the analysis of lower abundance precursors resulted in quantifying peptides spanning a wider range of abundances as shown in Fig. 2b. This wider dynamic range includes low abundance ranges not covered by the shotgun data at all. The median precursor intensity of peptides quantified by pSCoPE is 2.5-fold lower relative to the shotgun median, Fig. 2b.

Prioritization also increased data completeness, Fig. 2c,d. This increase is particularly pronounced for challenging peptides, as exemplified with a set of 1,000 peptides identified with less than 50% probability in shotgun SCoPE2 sets, Fig. 2c. Prioritizing these peptides in 8 representative runs increased data completeness by 171% compared to controlled shotgun runs, Fig. 2c. The gains in data completeness extended to peptides from all priority levels, albeit at a lower gain of about 16%. pSCoPE also increased data completeness at the protein level, reaching 93% for all proteins, which represents a 34% gain over the performance of shotgun analysis, Fig. 2d.

To put these gains in context, we compared them to other methods for increasing data completeness, such isobaric match-between-runs (iMBR)^31^. iMBR facilitated the identification and quantification of approximately 170 additional precursors per shotgun experiment compared to 2,595 additional precursors per experiment added by prioritization, Extended Data Fig. 4. In contrast to prioritization, the decrease in missing data afforded by iMBR cannot be applied to peptides pre-selected based on biological considerations.

**Figure 4.**
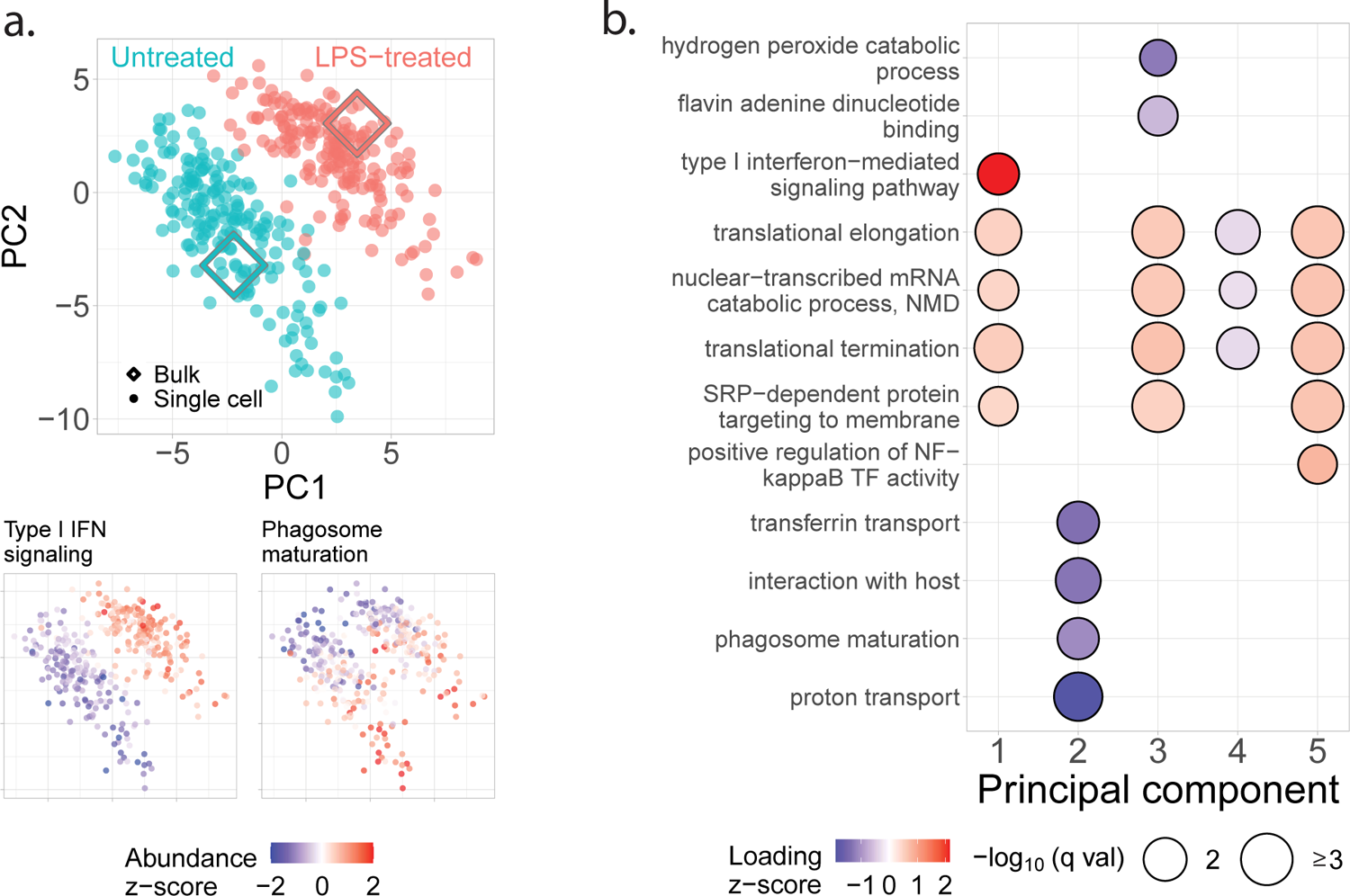
Prioritized analysis of primary macrophages identifies protein variation within and across treatment conditions. (**a**) Principal component analysis of 373 BMDM and 1,123 proteins color-coded by treatment condition. The diamond markers indicate bulk samples projected in the same low-dimensional space as the single cells. The adjoining PCA plots are color-coded by the z-scored median relative abundance of proteins corresponding to type I interferon-mediated signaling and phagosome maturation. Performing this analysis without imputation recapitulates these results, as shown in Extended Data Fig. 7. (**b**) Protein groups identified by protein set enrichment analysis (PSEA) performed using the PC vectors with protein weights from the PCA shown in panel a.

**Figure 5.**
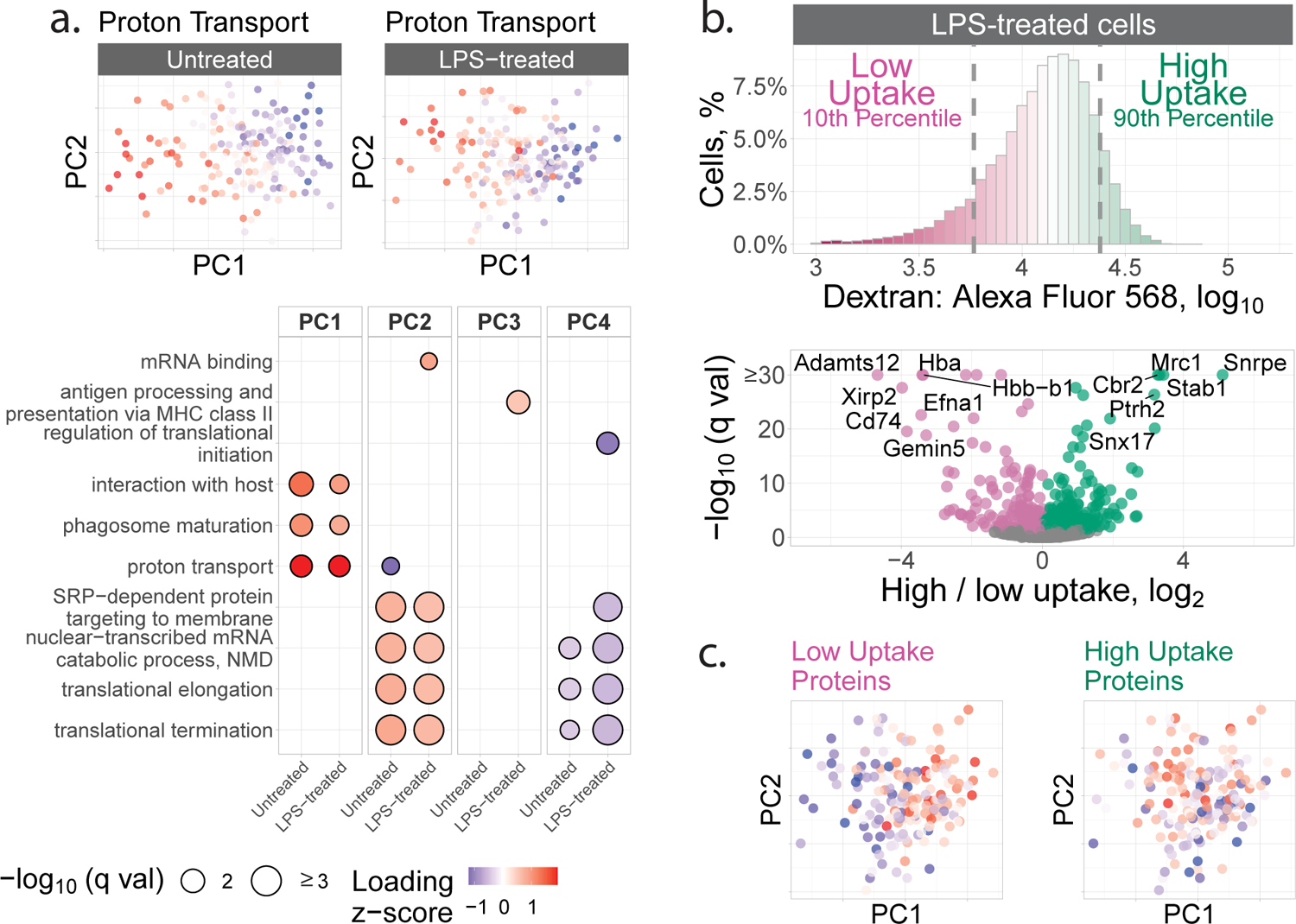
Axes of proteome polarization are similar between untreated and LPS-treated macrophages and correlated to dextran update. (**a**) Untreated and LPS-treated macrophages were analyzed separately by PCA and PSEA performed on the corresponding PCs. The PCA plots are color-coded by the median abundance of proteins annotated to proton transport. (**b**) The uptake of fluorescent dextran by LPS-treated macrophages was measured by FACS, and the cells with the lowest and highest uptake were isolated for protein analysis. The volcano plot displays the fold changes for deferentially abundant proteins and the associated statistical significance. The corresponding analysis for untreated macrophages is displayed in Extended Data Fig. 10. (**c**) The LPS-stimulated macrophages were displayed in the space of their PCs and color-coded by the median abundance of the low-uptake or the high-uptake proteins. The low-uptake proteins correlate to PC1 (Spearman *r* = 0.55, *q* ≤ 3 × 10−15) and the high-uptake proteins correlate to PC2 (Spearman *r* = 0.33, *q* ≤ 2 × 10^−5^).

The pSCoPE sets used for benchmarking performance in Fig. 2 included single cells from two different cell lines and thus allowed us to examine protein variation both between and within each cell line. These cell types were clearly separated by the first principal component (PC) of the data, Fig. 2e. Protein Set Enrichment Analysis (PSEA) performed on the PCs identified enrichment for multiple functional sets of proteins, including glycolysis and G2/M transition of mitotic cell cycle, as shown in Fig. 2e.

### Quantification accuracy of pSCoPE

Next, we benchmarked the quantitative accuracy of pSCoPE by comparing the mixing and measured ratios for a set of peptides spiked-in at known levels, Fig. 3. These peptides contained internal trypsin cleavage sites to control for digestion variability. They were spiked into 8 single-cell pSCoPE sets using a randomized design detailed in **Experimental design for spike-in analyses.** Schematic for the design of the single-cell sets used to benchmark reporter-ion quantitation in Fig. 3. Each row corresponds to a pSCoPE and having the indicated cell type and spike-in concentration for each sample. The sample column headers denote the TMTpro 18plex label associated with each sample, with RI5 indicating 128C, RI6 indicating 129N, and so on. In the “cell type” section for each experiment, “H” denotes HEK293 cells, “M” denotes melanoma cells, and “neg” denotes negative control samples, which were identically processed to the single-cell samples, but which did not contain a cell.. The spiked in levels were chosen and confirmed to span the abundance range of the peptides quantified in the single cells, Fig. 3a. Each peptide ranged in abundance over a 16-fold dynamic range across 5 spiked-in levels. The measured abundances exhibited linear dependence with the spiked-in levels with a slope of 1.06 and a coefficient of determination *R*^2^ = 0.97, Fig. 3b. These results indicate that pSCoPE is able to quantify peptides in single-cell sets with good accuracy and precision.

### Polarized proteome states

Next we used pSCoPE to explore the molecular and functional heterogeneity of murine bone-marrow-derived macrophages (BMDMs) responding to inflammatory stimuli, such as lipopolysaccharide (LPS), the major component of gram-negative bacteria’s outer membrane. The macrophages were differentiated using M-CSF and either treated with LPS for 24 hours or untreated. Single macrophage cells were prepared for MS analysis by nPOP^30^ and were first analyzed by SCoPE2^3, 32^. Peptides identified with high confidence and exhibiting high variability across the single cells were then prioritized for analysis by pSCoPE, using longer accumulation times and narrow isolation windows, as detailed in Methods. pSCoPE quantified 1,123 proteins across 373 single primary macrophage cells, achieving 71% data completeness for all proteins (Extended Data Fig. 6) and good quantitative agreement between peptides originating from the same protein (Extended Data Fig. 2c). The PCA projection of the data results in 2 clusters corresponding to the treatment conditions, Fig. 4a. Projected bulk samples cluster with the corresponding treatment groups, indicating that the cluster separation reflects treatment response^32^. This treatment-specific clustering is also reflected in the abundance of proteins that vary across treatments but not within a treatment, as exemplified by proteins functioning in the type-1 interferon-mediated signaling pathway, Fig. 4a.

**Figure 6.**
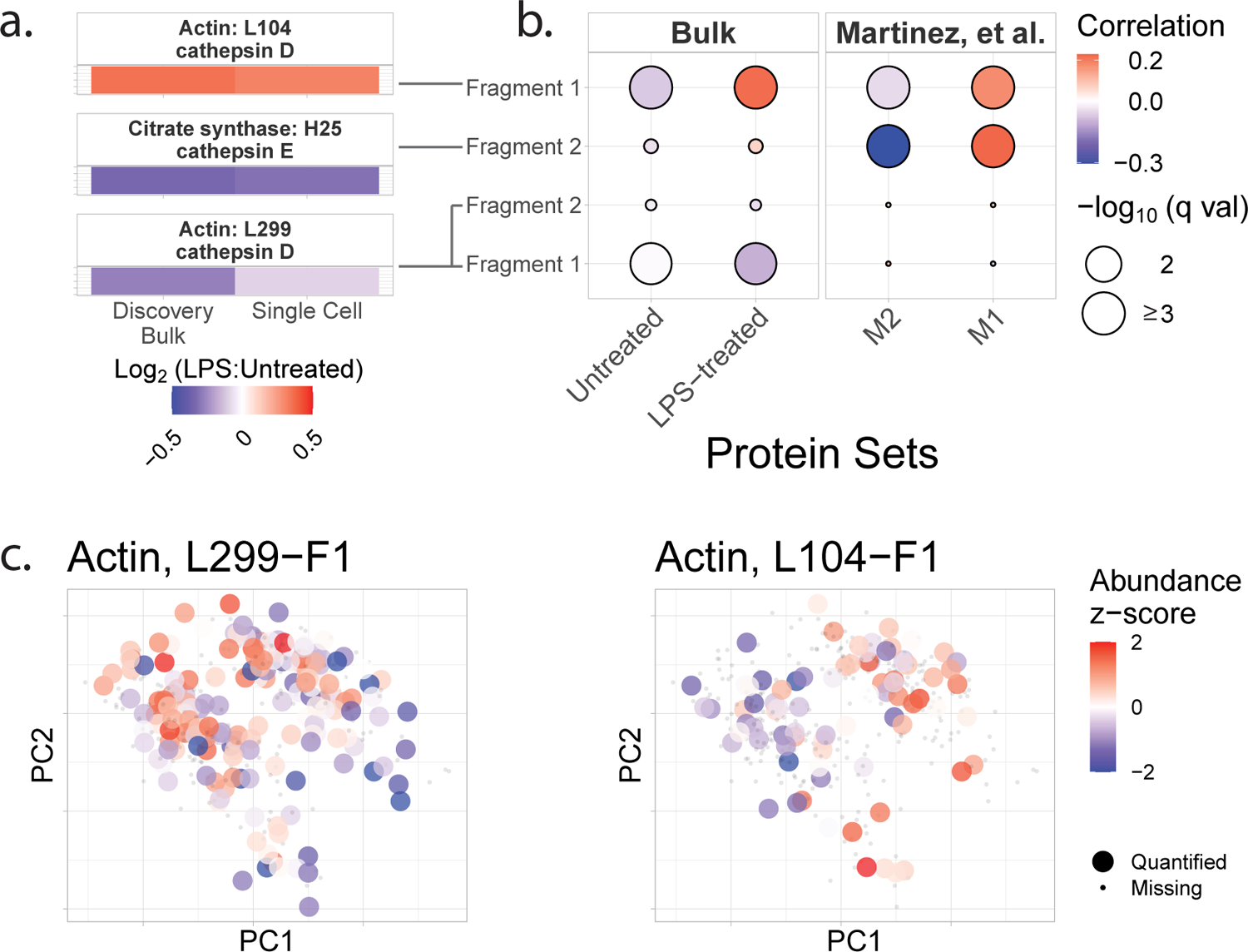
Proteolytic products in individual macrophages correlate to inflammatory markers and vary within treatment groups. (**a**) A comparisons between untreated and LPS-treated ratios of proteolytic products quantified in discovery bulk experiments and in single cells. The annotations are derived from the MEROPS database^35^. (**b**) Correlation analysis of proteolyic products to treatment-group-specific and macrophage-polarization-specific protein panels. (**c**) The untreated and LPS-treated cells were projected by PCA and color-coded by the relative abundance of the indicated actin fragments.

The spread of the macrophage clusters along PC1 suggests that proteins varying across treatment groups may also vary within a treatment group. Indeed, PSEA using the PCA loadings identified such protein sets, as exemplified with the phagosome maturation pathway, Fig. 4a-b. These finding are recapitulated when the PCA is performed without data imputation (Extended Data Fig. 7), suggesting robustness to choices of data analysis^33^. Additionally, color-coding the original PCA by per-sample data completeness indicates that the cross-condition and intra-condition sample separation is not driven by missing data, Extended Data Fig. 8. To systematically investigate proteome variations within a condition, we performed PCA of each treatment group separately and PSEA on the associated PCA protein loading. Remarkably, the first PCs of the treated and un-treated macrophages correlate strongly (*r* = 0.8, *p* < 10*^−^*^15^), suggesting that the within-condition protein variability is similar across the two conditions. This observation is naturally reflected in very similar functional enrichment results for treated and untreated macrophages, Fig. 5a.

These results suggest that a 24-hr LPS treatment does not fundamentally alter the axes of protein variation for murine BMDMs, such as phagosome maturation, proton transport, and protein targeting to the membrane, Fig. 5a. In addition to these shared functional groups, some protein sets vary only within the LPS-stimulated cells, as illustrated by proteins annotated to antigen processing and presentation via MHC class II, mRNA binding, and regulation of translational initiation, Fig. 5a. The coherence of protein variability within functional groups suggests it is functionally relevant^34^, but it does not prove it.

### Connecting protein variation to functional variation

To examine whether the observed protein heterogeneity has functional consequences, we sought to directly measure the endocytic activity of macrophages and its relationship to such protein heterogeneity. To this end, we measured the uptake of fluorescently labeled dextran and FACS sorted macrophages from the top and bottom deciles of the fluorescence distribution, Extended Data Fig. 9. Both the LPS-treated (Fig. 5b) and the untreated macrophages (Extended Data Fig. 10) exhibit large variance in dextran uptake per cell, with the median uptake being higher for the LPS-treated cells. The proteomes of subpopulations sorted based in their dextran uptake were analyzed by data independent acquisition, which allowed us to identify proteins whose abundance is significantly different between the most and least endocytically active cells, Fig. 5b and Extended Data Fig. 10. Then for each cell, we estimated the median abundances of these proteins associated with endocytic activity and correlated them to the PCs for each treatment condition. For the LPS-treated samples, the proteins associated with high dextran uptake (such as Mrc1, Stab1, and Snx17) were found to be significantly correlated to PC1, while the proteins annotated to low dextran uptake were inversely correlated to PC1 and significantly correlated to PC2, Fig. 5c. Notably, some proteins (such as Mrc1 and Stab1, and Cd74) exhibit similar association with dextran uptake both in the untreated and LPS-treated macrophages, Fig. 5b and Extended Data Fig. 10.

To more directly measure regulatory mechanisms, we sought to quantify proteolysis, which plays major functional roles in macrophage activation^36–38^. To avoid products of proteolysis that may occur during sample preparation, we focused only on proteolytic products present in the macrophages prior to trypsin digestion. These products were identified in discovery bulk samples in which amine groups were covalently labeled prior to trypsin digestion as commonly performed^39, 40^. The proteolytic products were matched to annotated proteolytic products in the MEROPS database^35^ and analyzed by pSCoPE in single cells. To evaluate the single-cell quantification, the fold-changes between LPS-treated and untreated cells were compared to the corresponding bulk estimates, Fig. 6a. The good agreement between the measurements from established bulk methods and pSCoPE support the accuracy of the single-cell quantification.

To infer the functional association of the validated proteolytic products, we correlated their single-cell abundances to the abundances of pro- and anti-inflammatory protein panels, Fig. 6b: (i) proteins that we identified as differentially abundant between bulk samples of untreated or LPS-treated macrophages, and (ii) previously reported markers for M1 or M2 macrophages^41^. The cathepsin-D-cleaved actin peptide (L104) and the cathepsin-E-cleaved citrate synthase peptide (H25) were found to be significantly positively correlated to inflammatory markers. Both peptide fragments annotated to cathepsin D cleavage at L299 were inversely correlated to the set of proteins which were more abundant in LPS-treated BMDMs.

Having established the reliability of single-cell quantification of proteolytic products across conditions, we next examined their abundance within a treatment condition, Fig. 6c. The data indicate that the actin proteolytic products exhibit significant variability within each treatment condition. For example, the actin fragment cleaved at L299 correlates significantly to PC1 (Spearman *r* = −0.32, *p* < 2 × 10*^−^*^5^), Fig. 6c. These results point to the possibility of using pSCoPE for analysing proteolytic activity at single-cell resolution.

## Discussion

Our analysis demonstrates the potential of prioritized data acquisition to simultaneously optimize multiple aspects of single-cell proteomics, including the consistency, sensitivity, depth, and accuracy of protein quantification, Fig. 1-3. These gains are achieved using multiplexed and widely accessible workflows^32^ and a new software module that is freely available. The performance gains by SCoPE2 demonstrate the potential of innovations in data acquisition to drive single-cell proteomics^13^. pSCoPE enabled us to quantify molecular and functional diversity of primary macrophages, even of post-transcriptionally modified (PTM) peptides (Fig. 6). This analysis of PTM peptides is enabled by (i) the ability of pSCoPE to selectively send for MS2 analysis even lowly abundant precursors and (ii) the narrow isolation windows that reduce the coisolation signal for such peptides. We expect these methodological benefits to generalize to diverse biological problems^42^.

Many MS methods allow for analyzing a pre-selected group of peptides. They range from targeted methods that maximize sensitivity and probability of target quantification^18–23, 43^ to directed methods that use inclusion lists^23, 43^. Some directed methods^25, 26^ can be used in hybrid mode with a small inclusion list and shotgun analysis (scan while idle), which can be seen as a single level of priority. However, these methods suffer from the low identification rate for MS2 scans acquired in shotgun mode. pSCoPE extends the directed family of methods by introducing a generalized tiered approach that allows the prioritization of thousands of peptides for isolation and fragmentation (thus achieving 96% success rate of sending high-priority peptides for MS2) while working at a full duty cycle and thus achieving high proteome coverage, Fig. 2. The multi-tiered approach allowed for high identification rates of MS2 scans from all tiers (including the low tiers) for an average sequence assignment of 84% at 1 % FDR even when using 0.5 Th isolation windows. This prioritized algorithm (Fig. 1) introduced here may also be implemented with other approaches for performing real-time retention time alignment^44, 45^ and may be extended to single-cell proteomics multiplexed by non-isobaric mass tags^46^.

We measured protein covariation of functionally related proteins within primary macrophages not only between treatment groups but also within a treatment group, as shown Fig. 4a for phagosome maturation proteins. The proteins exhibiting such within condition variability are similar for the treated and untreated macrophages, Fig. 5a. This similarity in protein covariation is remarkable because LPS treatment substantially remodels the proteome, and yet protein covariation remains similar for treated and untreated macrophages. A possible explanation for this finding is that protein covariation reflects the topology of regulatory interactions^34^, and many of these regulatory interactions remain similar between the untreated and LPS treated macrophages. This interpretation is consistent with the observation that the proteins associated with dextran uptake are similar for the two conditions as shown in Fig. 5b and Extended Data Fig. 10. The robustness of the results to choices of data analysis, such as different treatments of missing data (Extended Data Fig. 7), bolsters their validity^33^.

Our prioritization approach is broadly accessible as the software is free and compatible with the Thermo Fisher Q-Exactive series, Orbitrap Exploris as well as Orbitrap Eclipse, Supporting Fig. S1. The newer instruments have quadruples that are likely to substantially improve the efficiency of isolating ions with narrow isolation windows (0.5 Th) for MS2 scans, and thus achieve higher sensitivity and precision of quantification than those demonstrated here with a Q-Exactive classic instrument.

Prioritization can help increase the throughput of single-cell proteomics by enabling consistent analysis of proteins of interest on short chromatographic gradients^14^. All results presented here used 60 min active gradients, though shorter gradients may increase both the throughput and the sensitivity (via narrower elution peaks) while still affording enough time to analyze thousands of prioritized peptides^28^. Thus, pSCoPE may provide accurate and consistent protein quantification across many single cells to support sufficiently powered biological investigations of primary cells and tissues^14, 47^.

## Data Availability

Metadata, raw data, and processed data are organized according to community recommendations^33^ and are available at scp.slavovlab.net/Huffman et al 2022 and at MassIVE: MSV000090383.

## Code Availability

Code, and protocols are organized according to community recommendations^33^ and are available at scp.slavovlab.net/pSCoPE, and github.com/SlavovLab/pSCoPE.

## Acknowledgments

We thank L. Reiter and T. Gandhi for making Spectronaut available for this project and to T. Colombani and M. Jovanovic for discussions and constructive comments. This work was funded by a New Innovator Award from the NIGMS from the National Institutes of Health to N.S. under Award Number DP2GM123497, an Allen Distinguished Investigator award through The Paul G. Allen Frontiers Group to N.S., a Seed Networks Award from CZI CZF2019-002424 to N.S., and through a Merck Exploratory Science Center Fellowship, Merck Sharpe & Dohme Corp. to N.S.

## Correspondence

Correspondence and materials requests should be addressed to

## Author Contributions

Experimental design: R.G.H., N.S.

LC-MS/MS: R.G.H., A.L., H.S., J.D., S.K., A.P, E.E.

Sample preparation: R.G.H., A.L., M.G., F.B.

Cell culture: A.L., M.G., F.B, S.K., L.K., R.G.H.

Raising funding: N.S.

Supervision: N.S.

Data analysis: R.G.H., N.S.

Software development: C.W.

Initial draft: R.G.H., N.S.

Results interpretation: N.S, J.C., I.Z., R.G.H., A.L., H.S., J.D., S.K., M.G., A.P., D.H.P.

Writing: All authors approved the final manuscript.

## Methods

### Implementation of prioritized analysis

To maximize the probability of analyzing high-priority peptides (i.e., peptides of high experimental importance) when operating at full duty cycle, we developed a new feature of MaxQuant.Live^26^: *multi-tier prioritization*. Multi-tier prioritization uses the real-time instrument control capabilities of MaxQuant.Live and adds a priority feature that determines which precursors are analyzed when duty-cycle time becomes limiting. The initial priority for each peptide is a user-defined integer number which is by default set to zero. By assigning non-zero values, it is possible to prioritize a single set of peptides or to implement a multi-tier approach, depending on the experimental objectives. During data acquisition, the peptides are selected for fragmentation based on their priority. After each fragmentation event, the corresponding peptide priority value is decremented unless fragmentation occurred outside of the retention-time tolerance. The prioritization feature is part of the latest release of MaxQuant.Live (version 2.1), available at: MaxQuant.Live and scp.slavovlab.net/pSCoPE.

The initial user-defined priorities are set via a column in the inclusion list table. This column was added to allow for easy definition of priority for every peptide on the list. The higher the integer number associated with a peptide (and thus its tier), the higher the probability that it will be chosen for fragmentation when duty cycle is limited. MaxQuant.Live was tested on a Q Exactive (as described below), but it was written to be compatible with all Orbitrap instruments.

### Prioritization workflow

All prioritized single-cell experiments followed the four stages of the workflow displayed in Supporting Fig. S1 and described below.

1. Compilation of proteins of interest from literature or prior LC-MS/MS analyses. Detailed information regarding the construction of inclusion lists used in the analysis of the SQC samples, HEK and Melanoma samples, or BMDM samples can be found in the Prioritized Inclusion List Construction section, below.
2. DIA analysis of a 1x concentrated injection of the combined carrier-reference sample to generate accurate retention times for precursors which will subsequently be prioritized.
3. This step is enabled by using a spectral library generated from prior DIA analysis of a 5-10x concentrated injection of the combined carrier-reference samples.
4. Assignment of precursors identified in step 2 to priority tiers based on proteins of interest defined in step 1.
5. The minimal set of precursor characteristics needed for a prioritized inclusion list are the mass, expected apex retention time, and priority.
6. Acquire data from SCoPE samples using MaxQuant.Live’s prioritization feature and the inclusion list generated in step 3.
7. Performing a test run on a 1x injection of the combined carrier and reference samples can be useful for troubleshooting methods before acquiring data from single cells.

### Benchmarking MaxQuant.live with and without prioritization enabled, Fig. 1

These experimental sets, the results of which were presented in Fig. 1 were designed to benchmark the performance of prioritization against MaxQuant.Live’s default global targeting mode with respect to consistency of peptide identification across experiments, as well as protein coverage. The experiments presented in Fig. 1 are a matched set of six experiments acquired via MaxQuant.Live’s default global targeting mode and 6 experiments acquired with prioritization enabled. The parameters for experiments that directly compared MaxQuant.Live’s default operation and prioritized analysis were identical, including LC gradients and data acquisition parameters. Additional information regarding sample preparation, instrument parameters, MaxQuant.Live parameters, prioritized inclusion list design, analysis of raw data, single-cell data processing, and figure generation can be found in their respective sections. **The active gradient in all experiments was 60 min.**

### Comparing prioritized and shotgun analysis, Fig. 2

These experimental sets, the results of which were presented in Fig. 2, were designed to assess the relative performance of shotgun and prioritized methods with respect to sequence coverage and consistency of quantification across single-cell samples. The experiments presented in Fig. 2a are a matched set of eight shotgun analyses and eight prioritized analyses; the experiments presented in Fig. 2b,c,d,e are a matched set of eight shotgun analyses and eight prioritized analyses. The parameters for experiments that directly compared shotgun and prioritized analysis were identical, including LC gradients and data acquisition parameters with the only exception of increasing fill times for selected prioritized precursors as explicitly described in the main text. Additional information regarding sample preparation, instrument parameters, MaxQuant.Live parameters, prioritized inclusion list design, analysis of raw data, single-cell data processing, and figure generation can be found in their respective sections. **The active gradient in all experiments was 60 min.**

### Bone-Marrow-Derived Macrophage samples prepared by nPOP, Fig. 4/5/6

These experiments were designed to present a use case for prioritized LC-MS/MS methods. Twenty shotgun and forty prioritized single-cell experiments containing samples from both treatment conditions (untreated or treated for 24hrs with LPS) were conducted as part of this module. A side-by-side comparison of the twenty shotgun experiments and the first twenty prioritized experiments can be found in Extended Data Fig. 6. Only the results of the forty prioritized analyses were included in Fig. 4, Fig. 5, and Fig. 6. Additional information regarding sample preparation, instrument parameters, MaxQuant.Live parameters, prioritized inclusion list design, analysis of raw data, single-cell data processing, and figure generation can be found in their respective sections. **The active gradient in all experiments was 60 min.**

### Endocytosis experiments, BMDM samples, Fig. 5

In order to identify protein sets associated with endocytosis that were specific to murine BMDMs, bulk samples from each treatment condition (untreated or treated for 24hrs with LPS) were incubated with fluorescently labeled dextran, and samples from the top and bottom deciles of dextran uptake were isolated by FACS for downstream LC-MS/MS analysis. Protein sets found to be differential between dextran uptake deciles were then added to the top priority tier in subsequent prioritized analyses of single-cell BMDM samples. Additional information regarding sample preparation, instrument parameters, raw data analysis, and differential protein detection can be found in their respective sections.

### MEROPS experiments, BMDM samples, Fig. 6

Bulk BMDM samples from each treatment condition (untreated or treated for 24hrs with LPS) were lysed, cysteine residues were reduced and alkylated, and samples were incubated with TMTpro so that all pre-digestion n-termini would be distinguishable from neo-n-termini produced by a subsequent tryptic digestion. The raw LC-MS/MS data was then searched with a FASTA database containing all murine SwissProt reviewed sequences, as well as semitryptic peptides consistent with MEROPS-annotated proteolytic cleavage sites. These experiments were used to validate semitryptic MEROPS-annotated peptides observed in the prioritized single-cell samples. Additional information regarding sample preparation, MEROPS database integration, instrument parameters, and data analysis can be found in their respective sections.

### Bulk BMDM sample analyses by DDA and DIA

Bulk BMDM samples from each treatment condition (untreated or treated for 24hrs with LPS) were lysed, digested, and labeled with TMTpro for DDA analysis as a duplex sample or sequentially analyzed as labeled single-condition samples via DIA. These experiments were used to identify differentially abundant proteins between the treatment conditions which were then added to the top priority tier in subsequent prioritized analyses of single-cell BMDM samples. Additional information regarding sample preparation, instrument parameters, raw data analysis, and differential protein detection can be found in their respective sections.

### BMDM samples prepared via mPOP methods

This set of experiments represents an early troubleshooting investigation to both assess the sizes of the BMDMs from each treatment condition by using the cellenONE’s optical system (Scienion) and contrast against data generated in a prior set of single-cell BMDM samples isolated via FACS that may have experienced sorting issues. The results from this set were not used to generate any of the publication figures and are included merely for completeness, as a subset of identifications from these experiments informed the inclusion-list construction of the nPOP-prepared pSCoPE sets. Additional information regarding sample preparation, MEROPS database integration, instrument parameters, and raw data analysis can be found in their respective sections.

### LC-MS platform

The LC-MS/MS equipment and setup used for all analyses are detailed in the SCoPE2 protocol^32^. Briefly, samples were separated via online nLC on a Dionex UltiMate 3000 UHPLC; 1*µl* of sample was loaded onto a 25cm x 75*µm* IonOpticks Aurora Series UHPLC column (AUR2-25075C18A); mass spectrometry analyses were performed via a Thermo Scientific Q Exactive mass spectrometer; an Active Background Ion Reduction Device (ABIRD, by ESI Source Solutions, LLC, Woburn MA, USA) was used at the ion source to remove background contaminants. In the LC separations, buffer A was 0.1% formic acid in LC-MS-grade water, and buffer B was 80% Acetonitrile, 0.1% formic acid in LC-MS grade water; all buffer B percentages described in the subsequent instrument methods are relative to this concentration.

Instrument methods used in this study can be found in Supporting Table S2 and Supporting Table S5. Chromatographic methods used in this study can be found in Supporting Table S3 and Supporting Table S6.

### MaxQuant.Live parameters

Instrument method parameters followed the MQ.live listening scan guidelines: Two full MS - SIM scans were applied from minute 25 to 30 to trigger MaxQuant.live. Both MS - SIM scans had the following parameters in common: 70k resolution, 3e6 AGC target, and a 300ms maximum injection time. The first MS - SIM scan covered the scan space from 908Th to the total acquisition time plus 1000; for a total acquisition time of 95-minutes, the upper bound of the scan range would be 1070Th (95 minutes minues the initial 25 minutes before acquisition was triggered, plus 1000). The second MS - SIM scan covered the scan space from 909Th to the m/z corresponding to the MaxQuant.live method index to call.

The MaxQuant.Live parameters used for each sample analysis can be found in Supporting Table S10, Supporting Table S12, Supporting Table S11, and Supporting Table S13.

### Cell Culture

#### Culturing Melanoma cells

The melanoma cells (WM989-A6-G3, a kind gift from Sydney Shaffer, University of Pennsylvania) were cultured in TU2% media, composed of 80% MCDB 153 (Sigma-Aldrich M7403), 10% Leibovitz L-15 (ThermoFisher 11415064), 2 % fetal bovine serum (FBS, Millipore Sigma F4135), 0.5% penicillin-streptomycin (pen/strep, Thermo Fisher 15140122), and 1.68mM Calcium Chloride (Sigma-Aldrich 499609). Cells were passaged at 80% confluence in T75 flasks (Millipore-Sigma, Z707546) using 0.25% Trypsin-EDTA (ThermoFisher 25200072).

#### Culturing HEK293 cells

The HEK293 cells (CRL-1573, ATCC) were cultured in DMEM, supplemented with 10% fetal bovine serum (FBS, Millipore Sigma F4135) and 1% penicillin-streptomycin (pen/strep, Thermo Fisher 15140122). Cells were passaged at 80% confluence in T75 flasks (Millipore-Sigma, Z707546) using 0.25% Trypsin-EDTA (ThermoFisher 25200072).

#### Culturing and harvesting U937 cells

The U937 cells (CRL-1593.2, ATCC) cells were grown as suspension cultures in RPMI medium (HyClone 16777-145) supplemented with 10% fetal bovine serum (FBS, Millipore Sigma F4135) and 1% penicillin-streptomycin (pen/strep, Thermo Fisher 15140122). Cells were passaged when a density of 10^6^ cells/ml was reached. U937 cells were harvested by pelleting, before washing with 1x PBS at 4 °C. The washed cell pellets were diluted in 1x PBS at 4 *°C* and their density estimated by counting at least 1000 cells using a hemocytometer. Cells which were harvested for the SQC sample were resuspended in water (Optima LC/MS Grade, Fisher Scientific W6500.

#### Harvesting Melanoma and HEK cells

Prior to harvesting, media was removed from cell cultures, which were then rinsed with 0.25% Trypsin-EDTA (ThermoFisher 25200072) at 4 °C. After rinsing, adherent cultures were incubated with 4 °C 0.25% Trypsin-EDTA (ThermoFisher 25200072) for 15 minutes, until cells were detached from the culture vessel. Cold 1x PBS was added to each culture vessel, and the resulting suspension was pelleted via centrifugation at 250g, before being washed with 1x PBS and repelleted at 250g. The washed cell pellets were diluted in 1x PBS at 4 *°C* and their density estimated by counting at least 1000 cells using a hemocytometer. Cells which were harvested for carrier, reference, and SQC samples were resuspended in water (Optima LC/MS Grade, Fisher Scientific W6500) and frozen at −80 °C. Cells which were harvested for single-cell sorting on the cellenONE system were diluted in 1x PBS to a concentration of 300 cells/*µl* and placed on ice.

#### Culturing and harvesting Bone-Marrow-Derived Macrophages (BMDMs)

C57BL/6J (Jax 000664) mice were purchased from The Jackson Laboratory. Bone-marrow-derived macrophages (BMDMs) were differentiated from bone marrow in Dulbecco’s modified Eagle medium (DMEM; Thermo Fisher Scientific), 30% L929-M-CSF supernatant and 10% fetal bovine serum (FBS). After 7 days, BMDMs were replated at 1×10^6^ cells/ml in DMEM supplemented with 10% FBS, and each plate was either stimulated for 24 hours with LPS (Serotype O55:B5, Enzo Life Sciences) at 1*µg*/ml or allowed to rest. Prior to harvesting, cells were washed twice with 1x PBS and incubated with PBS-2mM EDTA to detach from the plate. Cells were then spun down at 300g for 5 minutes and washed with 1x PBS before being resuspended. The washed cell pellets were diluted in 1x PBS at 4 *°C* and their density estimated by counting at least 1000 cells using a hemocytometer. Cells which were harvested for carrier and reference samples were resuspended in water (Optima LC/MS Grade, Fisher Scientific W6500) and frozen at −80 °C. Cells which were harvested for single-cell sorting on the cellenONE system were diluted in 1x PBS to a concentration of 300 cells/*µl* and placed on ice.

### Spike-in peptide selection

Spike-in peptides were used in order to benchmark the accuracy and precision of reporter-ion quantitation in single-cell analyses. These spike-in peptides were selected on the basis of ionizeability and identifiability from the search results of an LC-MS/MS analysis of a yeast standard sample.

A DDA analysis of a TMT-labeled yeast standard sample, m13306.raw, was downloaded from Massive (MSV000084263), and searched with MaxQuant (v. 1.6.7.0). Trypsin/P was selected as the enzyme, and TMT (+224.152478 Da) was enabled as a variable modification on lysines and peptide n-termini. All other settings were kept as default. The *S. cerevisiae* reference proteome was downloaded from UniProt and used as the sequence database for this search (`uniprot-organism-yeast.fasta`).

The evidence.txt file from the MaxQuant search results was imported into the R environment. Peptides containing methionine, glutamine, and asparagine were removed from the search results, and peptides less than 9 amino acids and greater than 11 amino acids were removed. Peptides in the 25th percentile of PEP and the 75th percentile of precursor intensity and score were selected for further analysis. Peptide sequences present in the human proteome (`swissprot_human_20181210.fasta`) were also filtered out. Four tryptic sequences (AYF-TAPSSER, VEVDSFSGAK, TSIIGTIGPK, and ELYEVDVLK) from the filtered search results were then selected such that their retention times differed by more than a minute, subjecting them to different groups of coeluting peptides. These four sequences were then grouped into two pairs, and the pairs concatenated into single sequences (AYFTAPSSERVEVDSFSGAK and TSIIGTIGPKELYEVDVLK) for synthesis by JPT Peptide Technologies. These concatenated sequences were then used as trypsin-cleavable spike-in peptides to benchmark reporter ion quantitation.

### Sample preparation

#### Standards used for evaluating prioritization in Fig. 1

In order to provide a controlled comparison of MaxQuant.Live’s default global targeting method and the prioritized sample analysis method shown in Fig. 1, a standardized TMT-labeled sample was used (SQC standards).

Serially diluted bulk samples were multiplexed as previously described^3,32^. Each injection of this TMT-labeled sample contained one 50-cell-level carrier channel per cell type (U937: 126C; HEK293: 127N), three single-cell-level channels per cell type (U937: 128C, 129C, 130C; HEK293: 129N, 130N, 131N), one half-cell-level channel per cell type (U937: 131C; HEK293: 132N), and one quarter-cell-level channel per cell type (U937: 132C; HEK293: 133N).

#### Single-cell samples

All single-cell samples were prepared using the droplet nano-ProteOmic sample Preparation (nPOP) method as detailed in Leduc *et al.*, 2021^48^. In addition to sorted single cells, the SCoPE sets contained negative control samples to be used for downstream quality control purposes. These negative control samples received all reagents and proceeded through all sample handling steps, but no single cells were dispensed into these droplets^32^. The distribution of protein-level CVs (i.e. quantification variability) associated with the single cell and control samples for these experiments can be found in Extended Data Fig. 2a,b,c,d.

### HEK293 and melanoma single-cell sample preparation

A *∼*200-cell carrier and *∼*5-cell reference composed of HEK293 and melanoma cell lines were prepared using nPOP and following the guidelines of the SCoPE2 protocol^32^. In addition to serving as the carrier and reference for all single-cell sets analyzed in the technical section, the combined carrier and reference sample was used in all spectral-library-generation and retention-time-calibration experiments for the coverage and consistency experiments shown in Fig. 2.

### Spike-in peptide preparation for HEK293 and melanoma single-cell sample preparation

AYFTAPSSERVEVDSFSGAK and TSIIGTIGPKELYEVDVLK were ordered from JPT Peptide Technologies and resuspended at a concentration of 2.5 mM in LC-MS-quality water for storage at *−*20 °C. Spike-in concentrations of these two peptides were then examined empirically to determine a spike-in level where their tryptically digested fragments were readily detectable at MS1 and the associated reporter-ion intensities (when serially diluted across a 16-fold range) spanned the full dynamic range of endogenous-peptide reporter-ion intensities in single-cell samples. The lowest spike-in level was then denoted as the “1x” concentration. For the carrier and reference samples, the spike-in peptides were then added at a 400x and 20x concentration per set, respectively, so that they were 100-fold and 5-fold more abundant than then median spike-in level of 4x. For the 14 single-cell and control samples that were part of each SCoPE set, both spike-in peptides were serially diluted, dried down in a speed vac, and resuspended in 30 *µl* of LC-MS-quality DMSO, such that the following concentrations of each peptide were added to the indicated number of samples per set: 1x (2 samples), 2x (3 samples), 4x (3 samples), 8x (3 samples), 16x (3 samples). To achieve this addition, 5 DMSO aliquots containing the spike-in peptides were dispensed to form the 14-droplet clusters of each nPOP-prepared SCoPE set prior to cell dispensing; each droplet contained 8 nl of its respective spike-in dilution in DMSO. Spike-in amounts were randomized relative to TMT labels and cell types. Except for the addition of spike-in peptides, the single-cell samples were prepared as detailed in Leduc *et al.*, 2021^48^. Specifically, a single cell was added to each spike-in-contaning DMSO droplet, then digested, labeled, and quenched. The samples for each SCoPE set were pooled and transferred into the well of a 384-well plate that was loaded into the autosampler for LC-MS/M analysis.

### BMDM single-cell nPOP sample prep

Carrier and reference samples composed of equivalent amounts of untreated and LPS-stimulated murine BMDMs were prepared following the SCoPE2 protocol^29, 32^, such that the carrier was composed of *∼*200 cells and the reference was composed of *∼*5 cells. This sample design was then used in the preparation of single-cell sets by nPOP^48^, as well as in the generation of spectral libraries and retention-time-calibration experiments for the experiments shown in Fig. 4, Fig. 5, and Fig. 6, as well as Extended Data Fig. 6, Extended Data Fig. 7, and Extended Data Fig. 8.

The 24-hr LPS-treated and the untreated cells were combined within each SCoPE set. The majority (87%) of the labeled sets also contained negative control samples for quality-control purposes. These control samples received all reagents and proceeded through all sample handling steps, but no single cells were dispensed into these droplets. The distribution of protein-level CVs (i.e. quantification variability) associated with the single cell and control samples for these experiments can be found in Extended Data Fig. 2d.

### BMDM single-cell mPOP sample prep

Carrier and reference samples composed of equivalent amounts of untreated and LPS-stimulated murine BMDMs were prepared following the SCoPE2 protocol^29, 32^, such that the carrier was composed of *∼*200 cells and the reference was composed of *∼*5 cells. This sample design was then used in the preparation of single-cell sets by mPOP^49^, in which single-cells from each condition (untreated and 24-hr LPS treated) were sorted into a 384-well plate (Thermo AB1384) via the cellenONE liquid handling system (Scienion). The mixed carrier and reference sample was also used in the generation of retention-time estimate runs for the set of 10 samples analyzed by pSCoPE.

### Endocytosis assay samples

In order to facilitate an analysis of functional heterogeneity in the single-cell BMDM samples, markers of endocytic competency were identified from bulk analyses of untreated and 24hr LPS-treated BMDMs isolated by FACS on the basis of dextran uptake.

### BMDM endocytosis assay

Murine BMDMs were differentiated and divided into treatment groups, as indicated previously, and incubated with dextran conjugated to Alexa Fluor 568 (Thermo, D22912) at a final concentration of 0.5 mg/ml for 45 minutes at 37 °C. After the incubation period, cells were washed twice with 1x PBS and incubated with PBS-2mM EDTA to detach from the plate. Prior to FACS analysis, cells were spun down at 300g for 5 minutes and washed with 1x PBS before being resuspended. Using a Sony MA900, Dextran-AF568 fluorescence in the PE-Texas Red channel was then analyzed for cells from each treatment condition, and a minimum of 70,000 cells from the top and bottom *∼*10% of the PE-TR fluorescence distribution were then sorted for downstream sample preparation and mass spectrometry analysis.

### Preparation of endocytosis-assay samples for LC-MS/MS analysis

Each FACS-isolated sample was lysed using a freeze-heat cycle as part of mPOP^49^. Post-lysis, approximately 70,000 cells worth of lysate was digested for 12 hours at 37 °C using 11ng/*µl* of trypsin gold and 150 mM TEAB in 65*µl*. Samples were then stage-tipped^50^, and *∼*10,000 cells worth of digest was injected in 0.1% formic acid for analysis by mass spectrometry using via DIA method 5 using DIA gradient 4, detailed below.

### MEROPS Bulk validation experiments, BMDMs

In order to validate proteolytically regulated substrates detected in single-cell BMDM samples, LC-MS/MS analyses were performed on bulk samples prepared using a workflow previously applied to the identification and quantification of viral protease cleavage products^40^.

Murine BMDMs were differentiated, divided into treatment groups, and harvested, as indicated previously. Samples initially contained 125,000 BMDMs in 62.5 *µl* of LC-MS water (Optima LC/MS Grade, Fisher Scientific W6500). SDS (Sigma, L3771-100G) and HEPES (Thermo, Fisher Scientific AAJ63218AE) were added to final concentrations of 1% and 0.1M, respectively. cOmplete Protease inhibitor (Roche, Sigma Aldrich 05892791001) was then added to a 2x final concentration. The samples were then heated to 95 °C for 5 minutes and subsequently chilled at −80 °C for 10 minutes. 1U of benzonase (Millepore, Sigma Aldrich E1014-25KU) was added and allowed to incubate at room temperature for 30 minutes. 500mM DTT (Pierce, Thermo Fisher A39255) was added to a final concentration of 15mM and allowed to incubate for 30 minutes. Iodoacetamide (Pierce, Thermo Fisher A39271) was added to 15mM final concentration and incubated at room temperature in the dark for 30 minutes. DTT was then added a second time to a 15mM final concentration and incubated for 1 hr. SP3 Beads (Cytiva, Fisher Scientific 09-981-123; Cytiva, Fisher Scientific 09-981-121) were prepared and mixed following manufacturer recommendations.

2.5*µl* of prepared SP3 beads (100 *µg*/*µl*) were added to each of the four samples. 17.3*µl* of LC-MS grade water was added to each tube resulting in a total volume of 141*µl*. 564*µl* of ethanol (200 proof, HPLC/spectrophotometric grade, Sigma 459828-1L), was added to each sample and incubated for 18 minutes. Samples were then incubated for 5 minutes on a magnetic stand, the supernatant was removed, and the beads were washed twice with 400*µl* of 90% ethanol, after which the remaining supernatant was removed.

Each sample was resuspended in 22.5*µl* 6M GuCl (Sigma, G-3272), 30*µl* of 0.5M HEPES pH 8, and TCEP (10mM final concentration) (Supelco, Millepore Sigma 646547). Samples were then incubated for 30 minutes at room temperature. 57*µl* of TMTpro (Thermo, A44520) at 8ng/*µl* was then incubated in each sample for 1.5 hours, with the untreated condition being labeled with 127C and the LPS-treated condition being labeled with 128N. Samples were then quenched with 6*µl* of 1M TRIS (Thermo Fisher, AM9855G) for 45 minutes. Following quenching, 1.2*µl* of SP3 beads (100*µg*/*µl*) were added to each TMT-labeled sample. 484.4*µl* of 100% ethanol was added to each sample and allowed to incubate for 15 minutes. Samples were then placed on magnetic stand for 5 minutes, the supernatant was removed, and the beads were washed twice with 600*µl* of 90% ethanol. The samples were then centrifuged and the remaining liquid was removed.

Samples were resuspended in 100*µl* to a final concentration of 200mM HEPES and 12 ng/*µl* of trypsin gold (Promega, V5280). Samples were then placed in a bioshaker (Bulldog Bio, VWR 102407-834) and digested at 37 °C, 200 rpm for 18 hours. After digestion, samples were removed from the bioshaker, briefly sonicated, spun down, vortexed, spun down again, and incubated on a magnetic stand for 5 minutes. The supernatant was then removed and stored at −80 °C. Before analysis by LC-MS/MS, the samples were stage-tipped^50^. Samples were resuspended in 0.1% formic acid at approximately 1 *µg* worth of digest per *µl* in glass HPLC inserts (Thermo Fisher C4010-630) prior to analysis, then injected and analyzed via via DIA method 4 using DIA gradient 3, detailed below (raw files: eGH692-eGH694). TMT labeling was used in these experiments to facilitate identification of neo-N termini produced prior to tryptic digestion and was not used for multiplexed in-set quantitation; each TMT labeled sample was analyzed individually.

### Bulk TMTpro-labeled BMDM samples for differential protein analysis

10,000 cells from each treatment condition (24hr LPS-treated and untreated), resuspended in LC-MS water, were frozen at −80 degrees for 20 minutes, before being lysed at 90 °C in a thermal cycler (BioRad T1000) for 10 minutes. After lysis, benzonase was added to a final concentration of 1U and allowed to incubate for 10 minutes. Trypsin Gold (Promega Trypsin Gold, Mass Spectrometry Grade, PRV5280) was added to a final concentration of 16 ng/*µl* and triethylammonium bicarbonate (TEAB, Millipore Sigma T7408-100ML) was added to a final concentration of 150mM. The samples were then allowed to digest overnight for 16 hours. Post-digestion, samples were allowed to return to room temperature and were labeled with 85mM TMT 128N (untreated sample) or 85mM TMT 127C (LPS-treated sample). The reaction was then quenched with 0.5*µl* of 0.5% hydroxylamine (Millipore Sigma 467804-10ML) for 1 hr. Samples were centrifuged briefly to collect liquid following all reagent addition. After labeling, about 6000-cells worth of labeled material from each treatment condition were combined in a mass spec insert (Thermo Fisher C4010-630)) and dried down in a speed vacuum (Eppendorf, Germany) before being reconstituted in 3.3 *µl* 0.1% formic acid (Thermo Fisher 85178) and analyzed via shotgun MS instrument methods 1 and 2, using gradient 1, below.

Separate samples containing approximately 1000 cells per injection of the 128N-labeled untreated BMDMs or 127C-labeled 24-hr LPS-treated BMDMs were injected and analyzed via DIA bulk BMDM analysis instrument method 1, below. Each TMT-labeled sample was injected separately; TMT reporter ions were not used for in-set quantification in this analysis. Proteins which were differentially abundant between the two conditions analyzed via DIA were identified using the process outlined in the Differential protein analysis for DIA samples section, below, and these proteins make up set *ζ* in the description of the high priority level composition in the Prioritized Inclusion List Construction section, also found below.

### Spectral-library-generating samples

Prior to performing retention-time-calibration, scout, or prioritized experiments, spectral libraries were generated by analyzing bulk injections of SQC sample or mixed carrier and reference samples. These spectral libraries were used to facilitate precursor identification in the lower abundance retention time calibration samples.

For the SQC sample, 1*µl* injections of a 10x concentrated aliquot of the SQC sample were analyzed via DIA method 1 and 2 and DIA gradient method 1, and two subsequent 1*µl* injection of a 1x concentrated aliquot of the SQC sample were analyzed via DIA method 1 and DIA gradient method 1. For the HEK293 and melanoma samples, 1*µl* injections of a 10x concentrated aliquot of the mixed carrier and reference sample were analyzed by DIA instrument methods 1 and 2 and DIA gradient method 1, and two subsequent 1x concentrated aliquot of the mixed carrier and reference sample was injected and analyzed by DIA method 1 and DIA gradient method 1. For the BMDM samples, 1*µl* injections of a 5x concentrated aliqout of carrier and reference sample and a 1x concentrated aliquot of carrier and reference sample were sequentially analyzed via DIA Instrument Method 3 using DIA gradient method 1. Additional information regarding the instrument methods and search-engine parameters can be found in their respective sections.

### Scout experiments

Prior to assembling an inclusion list for prioritized SQC sample or single-cell sample analysis, a prioritized analysis of a 1x concentrated version of the SQC sample or a 1x concentrated version of the mixed carrier and reference sample was performed to generate a set of additional DDA-identifiable precursors. Information regarding the inclusion-list construction for these scout experiment, MaxQuant.Live parameters, and analysis of raw data can be found in their respective sections.

### Retention-time-calibration experiments

Retention-time-calibration experiments were used to generate accurate retention times for identifiable precursors to be used in subsequent scout experiments and prioritized single-cell analyses.

### SQC samples, Figure 1

A 1*µl* injection of a 1x concentrated aliquot of the SQC sample was analyzed via DIA method 1 using DIA gradient 1 and searched via DIA-NN (v.1.8.2 beta 2) with the spectral library generated from the corresponding spectral-library-generating experiments (`library_TMTpro.tsv`, 28,537 precursors). The precursor m/z range was set to 450-1600 Th, carbamidomethylation of cysteine was deselected as a fixed mod, the protease was set to trypsin, and the neural net classifier was set to double-pass mode. The following command line options were enabled: –no-ifs-removal, –full-unimod, and –report-lib-info. All other settings were left as default.

### HEK and melanoma samples, Figures 2/3

A 1*µl* injection of a 1x concentrated aliquot of the mixed carrier and reference sample was analyzed via DIA method 1 using DIA gradient 1 and searched via DIA-NN (v.1.8.1 beta 23) with the corresponding spectral-library-generating experiments (`Rebuttal_library.tsv`, 32,897 precursors). The precursor m/z range was set to 450-1600 Th, carbamidomethylation of cysteine was deselected as a fixed mod, the protease was set to trypsin, and the neural net classifier was set to double-pass mode. The following command line options were enabled: –no-ifs-removal and –report-lib-info. All other settings were left as default.

### BMDM nPOP samples, Figures 4/5/6

A 1*µl* injection of a 1x concentrated aliquot of mixed carrier and reference sample was injected and analyzed via DIA method 3 using DIA gradient 1 and searched via Spectronaut (v. 15.1) with the spectral library generated from the corresponding spectral-library-generating experiments (`20210809_120040_Priori_comb_080921.kit`). All search parameters were kept as default, except for: template correlation profiling enabled for profiling strategy, minimum q-value row selection for profiling row selection, and Biognosys’ iRT kit was indicated as not being used.

### BMDM mPOP samples

A 1*µl* injection of a 1x concentrated aliquot of mixed carrier and reference sample was injected and analyzed via DIA method 3 using DIA gradient 2 and searched by Spectronaut (v. 15.0) in directDIA mode using a FASTA containing the SwissProt database for *Mus musculus*, as well as MEROPS cleavage fragments generated as indicated in the MEROPS database preparation section, below (`musmusculus_SPonly_MEROPS_012221.fasta`, 27,117 protein entries). Trypsin was specified as the enzyme for in silico digestion, TMTpro (+ 304.2071 Da) was selected as a fixed modification on lysines, and the following variable mods were used: protein n-terminal acetylation (+ 42.01056 Da), methionine oxidation (+ 15.99492 Da), and TMTpro modification of peptide n-termini. The results were then prefiltered in Spectronaut to only contain precursors with at least one TMTpro modification. All other search settings were kept as default.

### Prioritized Inclusion List Construction

A mapping between inclusion lists and samples can be found in Supporting Table S7, Supporting Table S8, and Supporting Table S9.

### Scout experiments associated with the MaxQuant.Live feature contrast

A set of four prioritized analyses of the 1x mixed carrier and reference sample were conducted to generate a library of DDA-identifiable precursors from an initial DIA retention-time-calibration experiment. The search results from the retention-time-calibration experiment were filtered to include only fully labeled peptides, and peptide sequences with multiple charge states were condensed to the single most confidently identified charge state by PEP. This collection of peptides will be referred to as Group A, within this subsection. The data-processing pipeline available used in the construction of these inclusion lists is available at github.com/SlavovLab/pSCoPE.

- **Scout Run 1:** Peptides from Group A were stratified into high, medium, and low priority analysis groups on the basis of precursor intensity, such that peptides in the top third of intensities were placed in the high priority group (8,154 peptides), peptides in the middle third of intensities were placed in the middle priority group (7,914 peptides), and peptides in the bottom third of intensities were placed in the low priority group (7,915 peptides). This forms Inclusion List 1.
- **Scout Run 2:** Peptides from Group A were stratified into high, medium, and low priority analysis groups on the basis of precursor intensity, such that peptides in the top third of intensities were placed in the high priority group (3,775 peptides), peptides in the middle third of intensities were placed in the middle priority group (6,814 peptides), and peptides in the bottom third of intensities were placed in the low priority group (7,491 peptides). Peptides previously identified in a scout experiment were placed on a base priority level that was only sent for analysis to keep duty cycles full when no peptides of higher priority were available (5,903 peptides). This forms Inclusion List 2.
- **Scout Run 3:** Peptides from Group A were stratified into high, medium, and low priority analysis groups on the basis of precursor intensity, such that peptides in the top third of intensities were placed in the high priority group (2,558 peptides), peptides in the middle third of intensities were placed in the middle priority group (5,150 peptides), and peptides in the bottom third of intensities were placed in the low priority group (7,062 peptides). Peptides previously identified in a scout experiment were placed on a base priority level that was only sent for analysis to keep duty cycles full when no peptides of higher priority were available (9,366 peptides). This forms Inclusion List 3.
- **Scout Run 4:** Peptides from Group A were stratified into high, medium, and low priority analysis groups on the basis of precursor intensity, such that peptides in the top third of intensities were placed in the high priority group (1,677 peptides), peptides in the middle third of intensities were placed in the middle priority group (4,246 peptides), and peptides in the bottom third of intensities were placed in the low priority group (6,622 peptides). Peptides previously identified in a scout experiment were placed on a base priority level that was only sent for analysis to keep duty cycles full when no peptides of higher priority were available (11,591 peptides). This forms Inclusion List 4.

### MaxQuant.Live analysis of SQC sample with and without prioritization, Figure 1b/c

The search results from the retention-time-calibration experiment were filtered for use as an inclusion list via the data-processing pipeline available at github.com/SlavovLab/pSCoPE. The library of identifiable precursors referred to below was assembled from a set of four prioritized scout runs aimed at assembling a list of DDA identifiable peptides from the 1x retention-time calibration run. This library of PSMs was then filtered at 1% FDR, contaminants and reverse matches were removed, and multiple charge states of the same modified sequence were condensed to the entry with the highest spectral confidence of identification. While the inclusion list below (Inclusion List 5) was used for the default MaxQuant.Live global targeting analyses, as well, the prioritization feature was not enabled for those experiments.

- **High Priority:** Peptides from the filtered library referenced above (Group A) were then reduced to a subset that included the top-4 peptides per protein by spectral confidence (Group B). Group B was then filtered to only include peptides with spectral confidences *≤* 0.05 and precursor purities *≥* 0.8. An additional 810 peptides were added to this tier from Group B, such that they were the most confident previously unselected identifications whose PIFs were *≥* 0.5. This priority level featured 4,000 peptides.
- **Middle Priority:** Excluding peptides previously selected for a priority tier, Group B was filtered to only include peptides with PEPs *≤* 0.05 and precursor purities (PIFs) *≥* 0.5. This priority level featured 4,000 peptides.
- **Low Priority:** Excluding peptides previously selected for a priority tier, peptides from Group A were then filtered to only include peptides with PEP *≤* 0.05. This priority level featured 3,679 peptides.
- **Retention-time-calibration peptides:** All remaining precursors identified in the retention-time calibration run were selected to participate in the real-time retention-time-alignment algorithm, but disabled from being sent for MS2 analysis. This priority level featured 11,723 precursors.

### Scout experiments associated with HEK and Melanoma analyses

A set of four prioritized analyses of the 1x mixed carrier and reference sample were conducted to generate a library of DDA-identifiable precursors from an initial DIA retention-time calibration experiment. The search results from the retention-time-calibration experiment were filtered to include only fully labeled peptides, and peptide sequences with multiple charge states were condensed to the single most confidently identified charge state by PEP. This collection of peptides will be referred to as Group A, within this subsection. The data-processing pipeline available used in the construction of these inclusion lists is available at github.com/SlavovLab/pSCoPE.

- **Scout Run 1:** Peptides from Group A were stratified into high, medium, and low priority analysis groups on the basis of precursor intensity, such that peptides in the top third of intensities were placed in the high priority group (9,014 peptides), peptides in the middle third of intensities were placed in the middle priority group (8,749 peptides), and peptides in the bottom third of intensities were placed in the low priority group (8,749 peptides). This forms Inclusion List 6.
- **Scout Run 2:** Peptides from Group A were stratified into high, medium, and low priority analysis groups on the basis of precursor intensity, such that peptides in the top third of intensities were placed in the high priority group (4,467 peptides), peptides in the middle third of intensities were placed in the middle priority group (7,549 peptides), and peptides in the bottom third of intensities were placed in the low priority group (8,287 peptides). Peptides previously identified in a scout experiment were placed on a base priority level that was only sent for analysis to keep duty cycles full when no peptides of higher priority were available (6,209 peptides). This forms Inclusion List 7.
- **Scout Run 3:** Peptides from Group A were stratified into high, medium, and low priority analysis groups on the basis of precursor intensity, such that peptides in the top third of intensities were placed in the high priority group (2,316 peptides), peptides in the middle third of intensities were placed in the middle priority group (6,197 peptides), and peptides in the bottom third of intensities were placed in the low priority group (7,674 peptides). Peptides previously identified in a scout experiment were placed on a base priority level that was only sent for analysis to keep duty cycles full when no peptides of higher priority were available (10,325 peptides). This forms Inclusion List 8.
- **Scout Run 4:** Peptides from Group A were stratified into high, medium, and low priority analysis groups on the basis of precursor intensity, such that peptides in the top third of intensities were placed in the high priority group (1,467 peptides), peptides in the middle third of intensities were placed in the middle priority group (4,695 peptides), and peptides in the bottom third of intensities were placed in the low priority group (7,132 peptides). Peptides previously identified in a scout experiment were placed on a base priority level that was only sent for analysis to keep duty cycles full when no peptides of higher priority were available (12,948 peptides). This forms Inclusion List 9.

### HEK and melanoma samples, Figure 2a

The search results from the retention-time-calibration experiment were filtered for use as an inclusion list (Inclusion List 10) via the data-processing pipeline available at github.com/SlavovLab/pSCoPE.

The library of identifiable precursors referred to below was assembled from the corresponding shotgun analyses and a set of four prioritized scout runs aimed at assembling a list of DDA identifiable peptides from the 1x retention-time calibration run. This library of PSMs was then filtered at 1% FDR, contaminants and reverse matches were removed, and multiple charge states of the same modified sequence were condensed to the entry with the highest spectral confidence of identification. Peptides found to be identified in 50% or fewer of the shotgun analyses were excluded from this list of identifiable peptides.

- **High Priority:** Peptides from the filtered library referenced above (Group A) were then reduced to a subset that included the top-4 peptides per protein by spectral confidence (Group B). Group B was then filtered to only include peptides with PEPs *≤* 0.05 and PIFs *>* 0.8 for inclusion on the high priority level. This priority level featured 4,013 peptides.
- **Middle Priority:** Excluding peptides previously selected for a priority tier, Group B was filtered to only include peptides with PEPs *≤* 0.05 and PIFs *>* 0.7 for inclusion on the middle priority level. This priority level featured 4,166 peptides.
- **Low Priority:** Excluding peptides previously selected for a priority tier, Group A was then filtered to only include peptides with PEPs *≤* 0.05 for inclusion on the low priority level. This priority level featured 5,407 peptides.
- **Retention-time-calibration peptides:** All remaining precursors identified in the retention-time calibration run were selected to participate in the real-time retention-time-alignment algorithm, but disabled from being sent for MS2 analysis. This priority level featured 11,540 precursors.

### HEK and melanoma samples, Figure 2b/c/d/e

The search results from the retention-time-calibration experiment were filtered for use as an inclusion list (Inclusion List 11) via the data-processing pipeline available at github.com/SlavovLab/pSCoPE.

The library of identifiable precursors referred to below was assembled from the corresponding shotgun analyses and a set of four prioritized scout runs aimed at assembling a list of DDA identifiable peptides from the 1x retention-time-calibration run. This library of PSMs was then filtered at 1% FDR, contaminants and reverse matches were removed, and multiple charge states of the same modified sequence were condensed to the entry with the highest spectral confidence of identification.

- **High Priority:** Peptides from the filtered library referenced above (Group A) were then reduced to a set that had been identified in 50% or fewer of the associated shotgun experiments. Peptides with greater than 96% missing data in the single-cell samples were filtered out. Then peptides were subset into groups identified in 1, 2, 3, or 4 of the associated shotgun experiments, and 250 peptides from each of these groups were randomly sampled to form a 1000-peptide list of difficult-to-identify peptides.
- **Medium-High priority:** Excluding peptides previously selected for a priority tier, Group A was subset to include the top-4 peptides per protein by PEP (Group B). Group B was then filtered to only include peptides with PEPs *≤* 0.01 and PIFs *≥* 0.8. This priority level featured 3,475 peptides.
- **Medium-Low Priority:** Excluding peptides previously selected for a priority tier, the remaining library of identifiable precursors was filtered to only include peptides with PEPs *≤* 0.05 and PIFs *≥* 0.7. This priority level featured 4,146 peptides.
- **Low Priority:** Excluding peptides previously selected for a priority tier, the remaining library of identifiable precursors was filtered to only include peptides with PEPs *≤* 0.05. This priority level featured 5,009 peptides.
- **Retention time-calibration peptides:** Excluding peptides previously selected for a priority tier, the remaining library of precursors identified from the DIA retention-time-calibration run were selected to participate in the real-time retention-time-alignment algorithm, but disabled from being sent for MS2 analysis. This priority level featured 11,496 precursors.

### HEK and melanoma samples for spike-in analysis, Figure 3

The search results from the retention-time-calibration experiment were filtered for use as an inclusion list (Inclusion List 12) via the data-processing pipeline available at github.com/SlavovLab/pSCoPE. The library of identifiable precursors referred to below was assembled from the corresponding shotgun analyses and a set of four prioritized scout runs aimed at assembling a list of DDA identifiable peptides from the 1x retention-time calibration run. This library of PSMs was then filtered at 1% FDR, contaminants and reverse matches were removed, and multiple charge states of the same modified sequence were condensed to the entry with the highest spectral confidence of identification. Peptides found to be identified in 50% or fewer of the shotgun analyses were excluded from this list of identifiable peptides.

- **High Priority:** Identified precursors corresponding to the yeast-derived spike-in peptides (AYFTAPSSER, VEVDSFSGAK, TSIIGTIGPK, and ELYEVDVLK) were selected for this priority tier, which featured 8 precursors.
- **Medium-High Priority:** Peptides from the filtered library referenced above (Group A) were then reduced to a subset that included the top-4 peptides per protein by PEP (Group B). Group B was then filtered to only include peptides with PEPs *≤* 0.05 and PIFs *>* 0.8 for inclusion on the high priority level. An addition 134 peptides with PEPs *≤* 0.05 and PIFs *>* 0.5 was added from Group B. This priority level featured 4,000 peptides.
- **Medium-Low Priority:** Excluding peptides previously selected for a priority tier, Group B was filtered to only include peptides with PEPs *≤* 0.05 and PIFs *>* 0.7 for inclusion on the middle priority level. This priority level featured 4,695 peptides.
- **Low Priority:** Excluding peptides previously selected for a priority tier, Group A was then filtered to only include peptides with and FDR *≤* 1% for inclusion on the low priority level. This priority level featured 4203 peptides.
- **Retention time calibration peptides:** All remaining precursors identified in the retention-time calibration run were selected for this priority level. This priority level featured 9,258 precursors.

### Scout experiment for BMDM nPOP-samples, Figures 4/5/6

The search results from the retention-time-calibration experiment, Group A, were then filtered to meet the following criteria for use as an inclusion list (Inclusion List 13): elution group PEP *≤* 0.02, elution group q-value *≤* 0.05, TMTpro labeling modifications (+ 304.2071 Da) on the n-terminus or lysine residues. These filtered precursors form Group B.

- **High Priority:** Precursors from Group B in the top intensity tertile of Group A. This priority level featured 1,466 precursors.
- **Medium Priority:** Precursors from Group B in the middle intensity tertile of Group A. This priority level featured 2,429 precursors.
- **Low Priority:** Precursors from Group B in the low intensity tertile of Group A. This priority level featured 1,808 precursors.
- **Retention-time-calibration peptides:** Excluding peptides previously selected for a priority tier, the remaining library of precursors identified from the DIA retention-time-calibration run were selected to participate in the real-time retention-time-alignment algorithm, but disabled from being sent for MS2 analysis. This priority level featured 3,091 precursors.

### BMDM nPOP samples, Figures 4/5/6

The prioritized inclusion list for the BMDM samples (Inclusion List 14) was constructed by importing the search results from the DIA retention-time-calibration run into the R environment and subsetting the detected peptides into 4 tiers. Peptides and proteins of special biological interest or experimental utility were assembled from the following sources: all precursors identified at or below a PEP of 0.05 in the scout experiment (set *α*); proteins significantly correlated to PC1 (5% FDR) in a cross-condition PCA generated from the 20 initial shotgun analyses of single-cell BMDM samples (set *β*); proteins significantly correlated to PC1 (5% FDR) in a PCA of the LPS-treated single cells generated from the 20 initial shotgun analyses of BMDM samples (set *γ*); proteins significantly correlated to PC1 (5% FDR) in a PCA of the untreated single cells generated from 20 initial shotgun analyses of BMDM samples (set *δ*); proteins with *|log*_2_(fold changes)*| ≥* 1 between the low and high dextran uptake conditions of each treatment group analyzed by DIA which were found to be statistically significant, (set *E*, statistical process described in differential protein analysis for DIA samples section, below); proteins with *|log*_2_(fold changes)*| ≥* 1 between LPS-treated and untreated bulk BMDM samples analyzed via DIA which were found to be statistically significant, (set *ζ*, statistical process described in differential protein analysis for DIA samples section); proteins significantly correlated to PC1 (5% FDR, *|*Spearman correlation*| >* 0.35) from single-condition PCAs generated from prior mPOP-prepared single-cell BMDM analyses (set *η*); all precursors identified at 1% FDR in the 20 initial shotgun analyses of the single-cell BMDM samples (set *θ*); precursors identified in the retention-time-calibration run that were also contained in sets *β*, *γ*, and *δ* and identified in fewer than 50% of the corresponding shotgun analyses of single-cell BMDMs (set *ι*).

Regarding precursor-intensity-dependent fill times, if a top tier precursor appeared in the bottom intensity tertile, it was allotted an MS2 fill time of 1000ms; if a top tier precursor appeared in the middle intensity tertile, it was allotted an MS2 fill time of 750ms; if a top-tier precursor appeared in the top intensity tertile, it was allotted an MS2 fill time of 500ms. These intensity tertiles were calculated across all filtered PSMs from the retention-time-calibration run. All other precursors were allotted an MS2 fill time of 500ms.

- **High Priority:** Peptides from the retention-time-estimation run were selected to correspond to: up to the top 125 most abundant precursors from set *ι*; the top 35 most abundant MEROPS-annotated precursors; all precursors in set *θ* that mapped to proteins in set *ζ*; up to the top 100 most abundant precursors that were in common between set *θ* and precursors derived from proteins in set *η*; up to the top 100 most abundant precursors in common between set *θ* and precursors derived from proteins in set *E*; up to the top 5 most abundant precursors per protein in the intersection between set *θ* and precursors derived from proteins in set *δ*; up to the top 6 most abundant precursors per protein in the intersection between set *θ* and precursors derived from proteins in set *γ*; up to the top 4 most abundant precursors per protein in the intersection between set *θ* and precursors derived from proteins in set *β*. This priority level featured 589 precursors.
- **Medium Priority:** Excluding peptides previously selected for a priority tier, remaining precursors from set *θ*, the set of peptides identified at 1% FDR in the accompanying single-cell shotgun analyses, were selected for this priority level. This priority level featured 1353 precursors.
- **Low Priority:** Excluding peptides previously selected for a priority tier, all remaining peptides identified in the scout experiment, set *α*, were selected for this priority level. This priority level featured 2656 peptides.
- **Retention-time-calibration peptides:** Excluding peptides previously selected for a priority tier, all remaining peptides identified by the retention-time-calibration experiment were selected for participation in the real-time retention-time alignment algorithm, but were not enabled for MS2 analysis. This priority level featured 4675 peptides.

### mPOP-prepared BMDM troubleshooting samples

The search results from the DIA retention-time-calibration experiment were first filtered such that all remaining entries had an elution group PEP *<* 0.05 and an elution group Q value *<* .05., as well as at least one TMTpro modification (+ 304.2071 Da) on either the peptide n-terminus or lysine residue. The prioritized inclusion list (Inclusion List 15) for the mPOP BMDM samples was constructed such that the top tier contained the following types of precursors which intersected with the retention-time-calibration experiment identifications: precursors identified in less than 50% of the corresponding 13 shotgun experiments, precursors featuring a MEROPS-annotated cleavage site, precursors mapping to proteins of biological interest (TLRs,interleukin-associated proteins, lyosomal-associated membrane proteins, interferon-associated proteins, caspases, NF-kappaB-associated proteins, transcription factors, gasdermin, signal transducers, macrophage scavenger receptors, and proteins annotated to macrophage function). Precursors on the top priority tier were allocated fill-times dependent upon their precursor intensities in the following manner: precursors in the top abundance tertile were allocated a 600ms MS2 fill time, precursors in the middle abundance tertile were allocated a 750ms MS2 fill time, and precursors in the bottom abundance tertile were allocated a 900ms MS2 fill time. Precursors identified in the retention-time-calibration experiment that were part of a previous targeting experiment were placed on the middle priority tier along with the precursors that had been identified in the corresponding mPOP shotgun experiments. The bottom priority tier, although redundant to the top and middle tiers in composition, served to keep the instrument duty cycles full for optimal elution peak sampling. All precursors enabled for MS2, as well as all remaining precursors identified in the filtered retention-time-calibration run were enabled for participation in MaxQuant.Live’s real-time retention-time calibration algorithm.

### Analysis of raw MS data

#### MEROPS database preparation and FASTA modification

The MEROPS database was downloaded from https://www.ebi.ac.uk/merops/download35 and converted into a .csv for import into the R environment. All cleavage patterns consistent with trypsin (e.g., R or K as the P1 residue), were removed from the database. Then the SwissProt-annotated Mus musculus FASTA file was read into the R environment, and the sequence for each protein with an annotated MEROPS cleavage site was split between the P1 and P1’ residues. Both halves of the MEROPS-cleaved peptide were then subjected to an in silico tryptic digestion such that the tryptic digest produced a fragment at least 6 amino acids long. The two semi-tryptic peptide halves were then added to the existing FASTA as separate entries, including annotations from the MEROPS database for the enzyme, cleavage residue number, and whether the peptide fragment contained the neo-C terminus or neo-N terminus.

### DDA Data

#### Scout experiments of SQC samples associated with Figure 1

The four scout experiments associated with Figure 1 were searched with MaxQuant (1.6.17.0) using a FASTA containing all entries from the human SwissProt database and two yeast-derived spike-in proteins (`2022-06-20-reviewed-UP000005640_withSpikeIn.fasta`, 20,373 proteins). TMTpro 16plex was enabled as a fixed modification on n-termini and lysines via the reporter ion MS2 submenu. Methionine oxidation (+ 15.99492 Da) was enabled as a variable modification, and trypsin was selected for the in silico digestion with enzyme mode set to specific. Up to 2 missed cleavages were allowed per peptide with a minimum length of 7 amino acids. Second peptide identifications were disabled, calculate peak properties was enabled, and msScans was enabled as an output file. PSM FDR and protein FDR were set to 1. False discovery rate (FDR) calculations were performed in the R programming environment by calculating the PEP threshold at which 1% of the entries were decoy identifications.

#### SQC samples analyzed by MaxQuant.Live with and without Prioritization, Figure 1

The 6 matched analyses of the standard SQC sample conducted via MaxQuant.Live with and without prioritization enabled were searched with MaxQuant (1.6.17.0) using a FASTA containing all entries from the human SwissProt database and two yeast-derived spike-in proteins (`2022-06-20-reviewed-UP000005640_withSpikeIn.fasta`, 20,373 proteins). TMT-pro 16plex was enabled as a fixed modification on n-termini and lysines via the reporter ion MS2 submenu. Methionine oxidation (+ 15.99492 Da) was enabled as a variable modification, and trypsin was selected for the in silico digestion with enzyme mode set to specific. Up to 2 missed cleavages were allowed per peptide with a minimum length of 7 amino acids. Second peptide identifications were disabled, calculate peak properties was enabled, and msScans was enabled as an output file. PSM FDR and protein FDR were set to 1. False discovery rate (FDR) calculations swere performed in the R programming environment by calculating the PEP threshold at which 1% of the entries were decoy identifications.

#### Shotgun analyses of HEK and melanoma single-cell samples, Figure 2

Shotgun analyses of the HEK and melanoma samples were searched with MaxQuant (1.6.17.0) using a FASTA containing all entries from the human SwissProt database and two yeast-derived spike-in proteins (`2022-06-20-reviewed-UP000005640_withSpikeIn.fasta`, 20,373 proteins). TMTpro 18plex was enabled as a fixed modification on n-termini and lysines via the reporter ion MS2 submenu. Methionine oxidation (+ 15.99492 Da) and protein n-terminal acetylation (+ 42.01056 Da) were enabled as variable modifications, and trypsin was selected for the in silico digestion with enzyme mode set to specific. Up to 2 missed cleavages were allowed per peptide with a minimum length of 7 amino acids. Second peptide identifications were disabled, calculate peak properties was enabled, and msScans was enabled as an output file. PSM FDR and protein FDR were set to 1. False discovery rate (FDR) calculations were performed in the R programming environment by calculating the PEP threshold at which 1% of the entries were decoy identifications.

#### Scout experiments of HEK and melanoma carrier and reference materials associated with Figures 2/3

The four scout experiments associated with Fig. 2 and Fig. 3 were searched with MaxQuant (1.6.17.0) using a FASTA containing all entries from the human SwissProt database and two yeast-derived spike-in proteins (`2022-06-20-reviewed-UP000005640_withSpikeIn.fasta`, 20,373 proteins). TMTpro 18plex was enabled as a fixed modification on n-termini and lysines via the reporter ion MS2 submenu. Methionine oxidation (+ 15.99492 Da) was enabled as a variable modification, and trypsin was selected for the in silico digestion with enzyme mode set to specific. Up to 2 missed cleavages were allowed per peptide with a minimum length of 7 amino acids. Second peptide identifications were disabled, calculate peak properties was enabled, and msScans was enabled as an output file. PSM FDR and protein FDR were set to 1. False discovery rate (FDR) calculations were performed in the R programming environment by calculating the PEP threshold at which 1% of the entries were decoy identifications.

#### pSCoPE analyses of HEK and melanoma single-cell samples, Figures 2/3

In all 3 data sets, the prioritized runs corresponding to Fig. 2a, Fig. 2b/c/d/e, and Fig. 3, the same search settings were used as in the accompanying shotgun data sets, with the exception of only specifying methionine oxidation (+ 15.99492 Da) as a variable modification in the prioritized analyses. False discovery rate (FDR) calculations were performed in the R programming environment by calculating the PEP threshold at which 1% of the entries were decoy identifications.

#### Isobaric Match Between Runs Contrast, Figure S7

The shotgun analyses of HEK and Melanoma samples originally associated with Figure 2 were searched as in the associated section, but Match Between Runs was enabled. No changes were made to the search strategy employed for the prioritized analyses originally associated with Figure 2.

#### TMT-labeled bulk BMDM sample analyses for differential protein analysis

TMT-labeled and mixed bulk samples of 24-hr LPS-treated and untreated BMDMs analyzed via DDA MS Methods 1 and 2 and DDA Gradient Method 1 (wGH215 and wGH216, respectively) were searched with MaxQuant (v. 1.6.17.0) using a FASTA containing all entries from the SwissProt database for Mus musculus, as well as MEROPS-annotated cleavage products generated as indicated previously (`musmusculus_SPonly_MEROPS_012221.fasta`), for a total of 27,117 protein entries. TMTpro 16plex was enabled as a fixed modification on n-termini and lysines via the reporter ion MS2 submenu. Methionine oxidation (+ 15.99492 Da) and protein n-terminal acetylation (+ 42.01056 Da) were enabled as variable modifications, and trypsin was selected with specific cleavage. Second peptide identifications were disabled, calculate peak properties was enabled, and msScans was enabled as an output. PSM FDR and protein FDR were set to 1.

#### Scout experiment for inclusion-list generation for BMDM nPOP samples

The raw file generated by this prioritized analysis was searched by MaxQuant (v 1.6.17.0) using a FASTA containing all entries from the murine SwissProt database (`musmusculus_SPonly_012221.fasta`, 17,056 proteins). TMTpro 16plex was enabled as a fixed modification on n-termini and lysines via the reporter ion MS2 submenu. Methionine oxidation (+ 15.99492 Da) and protein n-terminal acetylation (+ 42.01056 Da) were enabled as variable modifications, and trypsin was selected with specific cleavage. Second peptide identifications were disabled, calculate peak properties was enabled, and msScans was enabled as an output. PSM FDR and protein FDR were set to 1.

#### Shotgun analyses of BMDM nPOP samples, Figures 4/5/6

Shotgun analyses of the nPoP-prepared murine BMDM samples were searched with MaxQuant (2.0.3.0) using a FASTA containing all entries from the murine SwissProt database with additional entries for cleaved peptides consistent with the MEROPS database appended (`musmusculus_SPonly_MEROPS_012221.fasta`), for a total of 27,117 protein entries. TMTpro was enabled as a fixed modification on n-termini and lysines via the reporter ion MS2 submenu. Methionine oxidation (+ 15.99492 Da) and protein n-terminal acetylation (+ 42.01056 Da) were enabled as variable modifications, and trypsin was selected for the in silico digestion with enzyme mode set to specific. Up to 2 missed cleavages were allowed per peptide with a minimum length of 7 amino acids. Second peptide identifications were disabled, calculate peak properties was enabled, and msScans was enabled as an output file. PSM FDR and protein FDR were set to 1.

#### pSCoPE analyses of BMDM nPOP samples, Figures 4/5/6

The same search settings were used as in the accompanying shotgun data sets, with the exception of the FASTA database. For the prioritized samples, a reduced version of the murine SwissProt database with appended MEROPS entries was used which contained only those proteins whose peptides were on the inclusion list (`musmusculus_SPonly_MEROPS_012221_lim2.fasta`; 1,234 proteins). Subsequent to the MaxQuant search, the 20 shotgun-analyzed nPOP-prepared SCoPE experiments, 40 pSCoPE-analyzed nPOP-prepared SCoPE experiments, and their preceding DIA-analyzed retention-time-calibration experiment were analyzed together by DART-ID^51^ for retention-time-dependent PSM confidence updating. A DART-ID configuration file is included in the Massive repository associated with this publication.

#### Shotgun and pSCoPE analyses of BMDM mPOP troubleshooting samples

Shotgun and pSCoPE analyses of the murine BMDM single-cell samples prepared by mPOP were searched with MaxQuant (1.6.7.0) using a FASTA containing all entries from the murine SwissProt database with additional entries for cleaved peptides consistent with the MEROPS database appended (`musmusculus_SPonly_MEROPS_012221.fasta`), for a total of 27,117 protein entries. TMTpro was enabled as a fixed modification on n-termini and lysines via the reporter ion MS2 submenu. Methionine oxidation (+ 15.99492 Da) and protein n-terminal acetylation (+ 42.01056 Da) were enabled as variable modifications, and trypsin/P was selected for the in silico digestion with enzyme mode set to specific. Up to 2 missed cleavages were allowed per peptide with a minimum length of 7 amino acids. Second peptide identifications were disabled, calculate peak properties was enabled, and msScans was enabled as an output file. PSM FDR and protein FDR were set to 1.

### DIA Data

DIA analysis was used to identify many precursors and their associated retention times, so that they can be used to compiling inclusion lists, Supporting Fig. S1.

### SQC samples, spectral-library-generating search

1*µl* injections of a 10x concentrated aliquot of the SQC sample were analyzed by DIA instrument methods 1 and 2 using DIA gradient 1, and 2 1*µl* injections of a 1x concentrated aliquot of the SQC sample were analyzed by DIA instrument method 1 using DIA gradient 1. These four sample analyses were used to construct a spectral library via FragPipe for the analysis of a 1x mixed carrier and reference sample analyzed via DIA instrument methods 1 using DIA gradient 1 (i.e., a retention-time-calibration experiment). The spectral library contained a total of 28,537 precursors, and was generated from a directDIA search using `2022-06-20-decoys-reviewed-contam-UP000005640_SpikeIns.fasta` (20,373 proteins). Default search parameters were used from the `DIA_speclib_quant` workflow, with the following exceptions: the enzyme was set to trypsin; n-terminal acetylation, nQnC, and nE were disabled as variable modifications; Carbamidomethylation of cysteine residues was disabled as a fixed modification; TMTPro (+ 304.207146 Da) on peptide n-termini and lysine residues were enabled as fixed modifications; quantify with DIA-NN was disabled. The spectral library produced by this search was named `library_TMTPro.tsv`.

### HEK and melanoma samples, spectral-library-generating search

1*µl* injections of a 10x concentrated aliquot of the mixed carrier and reference samples were analyzed by DIA instrument methods 1 and 2 using DIA gradient 1, and 2 1*µl* injections of a 1x concentrated aliquot of the mixed carrier and reference samples were analyzed by DIA instrument method 1 using DIA gradient 1. These four sample analyses were used to construct a spectral library via FragPipe for the analysis of a 1x mixed carrier and reference sample analyzed via DIA instrument methods 1 using DIA gradient 1 (i.e., a retention-time-calibration experiment). The spectral library contained a total of 32,897 precursors, and was generated from a directDIA search using `2022-06-20-decoys-reviewed-contam-UP000005640_SpikeIns.fasta` (20,373 proteins). Default search parameters were used from the `DIA_speclib_quant` work-flow, with the following exceptions: the enzyme was set to trypsin; n-terminal acetylation, nQnC, and nE were disabled as variable modifications; Carbamidomethylation of cysteine residues was disabled as a fixed modification; TMTPro (+ 304.207146 Da) on peptide n-termini and lysine residues were enabled as fixed modifications; quantify with DIA-NN was disabled. The spectral library produced by this search was named `Rebuttal_library.tsv`.

### BMDM samples, spectral-library-generating search

1*µl* injections of both a 5x concentrated aliquot and a 1x concentrated aliquot of mixed carrier and reference samples were sequentially analyzed via DIA instrument method 3 using DIA gradient 1. The two sample analyses indicated above were used to construct a spectral library using Spectronaut (v. 15.1) via a directDIA search using the `musmusculus_SPonly_MEROPS_012221.fasta` (27,117 proteins). Methionine oxidation (+ 15.99492 Da), n-terminal acetylation (+ 42.01056 Da), and n-terminal TMTpro labeling (+ 304.2071 Da) were enabled as variable modifications, while TMTpro labeling of lysines was enabled as a fixed modification. The resulting spectral library contained a total of 11,701 precursors. Default search parameters were used, with the following exceptions: allow source specific iRT calibration was enabled, Biognosys’ iRT-kit alignment peptides were specified as unused, and profiling strategy was set to template correlation profiling. The spectral library produced by this search was named `20210809_120040_Priori_comb_080921.kit`.

### SQC sample, retention-time-calibration experiment, Figure 1b/c

Raw data was searched via DIA-NN (v. 1.8.2 beta 2) with the sample-specific spectral library (`library_TMTPro.tsv`, 28,537 precursors) discussed above to provide accurate retention times for the subsequent MaxQuant.live-enabled prioritized single-cell analyses. The reference FASTA for this spectral library was `2022-06-20-decoys-reviewed-contam-UP000005640_SpikeIns.fasta` (20,373 proteins). The protease was set to Trypsin, N-term M excision was enabled, carbamidomethylation of cysteine residues was disabled, Precursor m/z range was set to 450 to 1600Th, fragment m/z range was set to 200 to 2000Th, double-pass mode was enabled, and the following command line arguments were enabled: the –no-ifs-removal, –full-unimod, and –report-lib-info. All other options were kept as default.

### HEK and melanoma samples, retention-time-calibration experiment, Figures 2/3

Raw data was searched via DIA-NN (v. 1.8.1 beta 23) with the sample-specific spectral library (`Rebuttal_library.tsv`, 32,897 precursors) discussed above to provide accurate retention times for the subsequent MaxQuant.live-enabled prioritized analyses. The reference FASTA for this spectral library was `2022-06-20-decoys-reviewed-contam-UP000005640_SpikeIns.fasta` (20,373 proteins). The protease was set to Trypsin, N-term M excision was enabled, carbamidomethylation of cysteine residues was disabled, Precursor m/z range was set to 450 to 1600Th, fragment m/z range was set to 200 to 2000Th, double-pass mode was enabled, and the following command line arguments were enabled: –no-ifs-removal and –report-lib-info. All other options were kept as default.

### BMDM samples, pre-scout retention-time-calibration experiment, Figures 4/5/6

Raw data was searched via Spectronaut (v. 15.1) with the BMDM-specific spectral library: `20210809_120040_Priori_comb_080921.kit`. The reference FASTA for this spectral library was `musmusculus_SPonly_MEROPS_012221.fasta` (27,117 proteins). The iRT-kit alignment peptides were specified as unused, template correlation profiling was selected as the profiling strategy, minimum q-value row selection was selected for the profiling row selection method, “allow source specific iRT calibration” was set to true. All other options kept as default

### BMDM nPOP samples, retention-time-calibration experiment, Figures 4/5/6

Raw data was searched via Spectronaut (v. 15.1) with the following spectral library: `20210809_120040_Priori_comb_080921.kit`. The reference FASTA for this spectral library was `musmusculus_SPonly_MEROPS_012221.fasta` (27,117 proteins). The iRT-kit alignment peptides were specified as unused, template correlation profiling was selected as the profiling strategy, minimum q-value row selection was selected for the profiling row selection method, “allow source specific iRT calibration” was set to true. All other options kept as default

### TMTpro-labeled bulk BMDM samples for differential protein analysis

TMT-labeled, unmixed bulk samples (wGH217/218/220) analyzed by DIA Instrument Method 3 and DIA Gradient Method 1 were searched with Spectronaut (v. 14.1) in directDIA mode using `musmusculus_SPonly_MEROPS_012221.fasta` containing 27,117 proteins. Methionine oxidation (+ 15.99492 Da), n-terminal acetylation (+ 42.01056 Da), and n-terminal TMTpro labeling (+ 304.2071 Da) were enabled as variable modifications, while TMTpro labeling of lysines was enabled as a fixed modification. Trypsin was specified as the enzyme for in silico digestion, and results were filtered to contain only precursors with TMTpro labeling modifications.

### TMTpro-labeled bulk MEROPS validation samples for BMDM analysis, Figure 6

The Raw files from the MEROPS analyses were then searched with Spectronaut’s (v. 15.4) direct-DIA analysis feature, using a FASTA containing all entries from the SwissProt database for Mus musculus, as well as MEROPS cleavage fragments generated as indicated previously, which contained 27,117 protein entries (`musmusculus_SPonly_MEROPS_012221.fasta`). Cysteine carbamidomethylation was set as a fixed modification and the following variable modifications were used: protein n-terminal acetylation (+ 42.01056 Da), methionine oxidation (+ 15.99492 Da), TMTpro modification (+ 304.2071 Da) of lysine and peptide n-termini. Trypsin enzymatic cleavage rules were enabled, allowing for a minimum peptide length of 7 and a maximum peptide length of 52. Up to 2 missed cleavages were allowed. All other search settings were left at their default values.

### Label-free bulk endocytosis samples for BMDM analysis, Figure 5

Raw data from the bulk endocytosis sample analyses was searched with Spectronaut (v. 14.10) via DirectDIA, using a FASTA containing all entries from the SwissProt database for Mus musculus, as well as selected isoforms and MEROPS-annotated cleavage products generated as indicated previously, for a total of 33,996 protein entries (`Mouse_ONLYsp_plusMEROPS_v2.fasta`). Peptides with lengths between 6 and 52 amino acids, with up to 2 missed cleavages were permitted. Trypsin/P was selected for cleavage, and Protein n-term acetylation (+ 42.01056 Da) and methionine oxidation (+ 15.99492 Da) were enabled as variable modifications. A PEP Cut-off of 1 was selected, although downstream filtration (PEP *≤* 0.01) was performed in the differential protein analysis script. All other search settings were kept as default.

### BMDM mPOP samples retention-time-calibration experiment

The pre-prioritization retention-time-calibration experiment was searched by Spectronaut (v. 15.0) in directDIA mode using a FASTA containing the SwissProt database for Mus musculus, as well as MEROPS cleavage fragments generated as indicated previously, containing 27,117 protein entries (`musmusculus_SPonly_MEROPS_012221`). Trypsin was specified as the enzyme for in silico digestion, TMTpro (+ 304.2071 Da) was selected as a fixed modification on lysines, and the following variable mods were used: protein n-terminal acetylation (+ 42.01056 Da), methionine oxidation (+ 15.99492 Da), and TMTpro modification of peptide n-termini. The results were then prefiltered in Spectronaut to only contain precursors with at least one TMTpro modification. All other search settings were kept as default.

### Processing and normalizing single-cell MS data

#### Shotgun and pSCoPE HEK and Melanoma analyses, Figures 2

Sets pertaining to Fig. 2a and Fig. 2b/c/d/e were processed separately. Single-cell MS data were processed via the SCoPE2 single-cell proteomics pipeline^3,32^. Peptides with precursor ion fractions below 50% or a mean RI intensity across the single cells greater than 10% of the intensity in the carrier channel were removed from the data set. Peptides were filtered at 1% FDR by determining the PEP threshold at which 1% of the entries were reverse matches. Cells with mean protein CVs greater than 0.4 were filtered out from the data set. Samples and precursors were then filtered to have less than 99% missing data, before being log transformed and aggregated to protein-level abundance by taking the median abundance of the protein-specific peptides. The following intermediate data frames were generated for subsequent analysis from both data sets: the matrix of single cells by precursors, unfiltered for missingness; the matrix of unimputed protein abundances by single-cells, containing missing values. For the sets corresponding to Fig. 2b/c/d/e, the following additional data matrices were produced: the complete matrix of protein abundances by single cells, containing imputed values; the batch-corrected complete matrix of protein abundances by single cells, containing imputed values; the re-normalized batch-corrected complete matrix of protein abundances by single cells, containing imputed values.

#### pSCoPE analyses of BMDM samples, Figures 4/5/6

Single-cell data from 40 prioritized analyses were processed via the single-cell pipeline^3,32^. Peptides with precursor ion fractions below 50% were removed from the data set, as were peptides with a mean intensity across the single cells greater than 2% of the intensity in carrier channel. Peptides were filtered at 1% FDR using the DART-ID^51^ q-value column. Cells with mean protein CVs greater than 0.4 were filtered out from the data set. Samples and precursors were then filtered to have less than 99% missing data, before being log transformed and aggregated to protein-level abundance by taking the median abundance of the protein-specific peptides. The following intermediate data frames were generated for subsequent analysis: the matrix of single cells by precursors, unfiltered for missingness; the matrix of unimputed protein abundances by single-cells, containing missing values; the complete matrix of protein abundances by single cells, containing imputed values; the batch-corrected complete matrix of protein abundances by single cells, containing imputed values; the re-normalized batch-corrected complete matrix of protein abundances by single cells, containing imputed values.

#### Shotgun and pSCoPE analyses of BMDM samples, Extended Data Figure 6

Single-cell data from 20 shotgun analyses and the first 20 prioritized analyses were processed via the single-cell pipeline^3,32^ using the same parameters as indicated above for the pSCoPE-only BMDM analyses.

### Data Analysis

#### Differential protein analysis for DIA samples

Differential protein abundance was assessed by modeling the distribution of noise as a function of average precursor intensity. To perform this analysis, a single sample was injected and analyzed twice by DIA, then searched with Spectronaut as described above. Precursor-level fold-changes between replicate injections of the same sample should cluster around 1:1, deviations from this expected 1:1 ratio reflect noise in the measurement, and can be used as a null distribution to test for differential protein abundance. However, because precursor quantitation is more accurate at higher absolute intensities, the null distribution of precursor fold-changes were split evenly into 15 bins with respect to the average precursor intensity of the pair. Fold-changes between experimental conditions were then calculated, and converted to a z-score using its corresponding null distribution of fold-changes (based on intensity). Lastly, the precursors for each protein were t-tested against a standard-normal null distribution of 10,000 values with mean = 0, and standard deviation = 1, then p-values were converted to q values using the Benjamini and Hochberg approach.

### Figure 1, MaxQuant.Live contrast with and without prioritization

#### Consistency of identification bar plot

The evidence.txt file containing the 6 SQC experiments acquired by MaxQuant.Live without prioritization enabled (wGH0727,wGH0729,wGH0731,wGH0733,wGH0735,wGH0737) and with prioritization enabled (wGH0728,wGH0730,wGH0732,wGH0734,wGH0736,wGH0738) were filtered at 1% FDR, and new column for precursor identity was created by concatenating the modified sequence and charge state of each entry. Entries were then aggregated by precursor on a per-experiment basis, selecting the identification with the lowest PEP. This data frame was then reduced to a column of precursor identities and raw files. The number of experiments in which a precursor was identified was tallied for each analysis method, and these per-platform tallies were then left joined to the inlclusion list, and resulting NA values for precursors not identified in any experiment were set to 0. The per-analysis-method per-precursor sums were then divided by the total number of experiments in each analysis set (6) and multiplied by 100 to generate the identification rate as a percentage. The data was then subset to only include information for precursors in the high priority group (4000 precursors), and the resulting data frame was summarized such that the number of precursors identified with a particular frequency was tallied on a per-analysis-method basis; this data was plotted to produce the leftmost plot in Fig. 1b.

#### Proteins quantified per run boxplot

The evidence.txt file containing the 6 SQC experiments acquired by MaxQuant.Live without prioritization enabled (wGH0727,wGH0729,wGH0731,wGH0733,wGH0735,wGH0737) and with prioritization enabled (wGH0728,wGH0730,wGH0732,wGH0734,wGH0736,wGH0738) were filtered at 1% FDR, and the tally of unique proteins quantified per experiment was conducted using the Leading.razor.protein column. These tallies were then presented as box plots in rightmost plot of Fig. 1b.

#### Prioritized peptides detected at MS1 and sent for MS2 analysis boxplot

The MaxQuant.Live log files corresponding to the 6 experiments for each analysis method were parsed using an R script available on github.com/SlavovLab/pSCoPE to extract the list of precursors detected at MS1 and sent for MS2 analysis. The precursors detected at MS1 or sent for MS2 were then summed on a per-experiment and per-priority-level basis and divided by the total number of precursors corresponding to that priority level on the inclusion list, before being multiplied by 100 to yield a percentage detection or isolation rate. These per-experiment and per-priority-level detection and isolation percentages were then plotted in Fig. 1c, with each point corresponding to an experiment, of which there were six for each analysis method.

### Figure 2, HEK and Melanoma samples

#### Productive MS2 scans boxplot

The msms.txt files from the shotgun and pSCoPE MaxQuant search results were filtered at 1% FDR, by determining the PEP threshold at which 1% of the entries were reverse matches. All contaminant and reverse matches were then removed from the resulting data frames. The number of remaining PSMs was tallied and divided by the total number of MS2 scans recorded per experiment, determined from the msmsScans.txt output of MaxQuant. This fraction was multiplied by 100 and presented in Fig. 2a as the percentage of productive MS2 scans per experiment.

#### Peptides/run boxplot

The evidence.txt files from the shotgun and pSCoPE MaxQuant search results were filtered at 1% FDR, by determining the PEP threshold at which 1% of the entries were reverse matches. All contaminant and reverse matches were then removed from the resulting data frames, and peptides with multiple charge states were collapsed to a single entry per experiment. The number of PSMs remaining was then tallied on a per experiment basis and presented as a boxplot in Fig. 2a.

#### Proteins/cell boxplot

Using the matrix of unimputed protein abundances by single-cell samples produced by the single-cell pipeline^3, 32^, the number of proteins per single-cell sample with detectable reporter ion intensities was tallied and presented in Fig. 2a.

#### Sensitivity analysis

Eight matched experiments for shotgun (wGH0643, wGH0644, wGH0645, wGH0646, wGH0647, wGH0648, wGH0649, wGH0650) and pSCoPE (wGH0660, wGH0661, wGH0663, wGH0665, wGH0666, wGH0667, wGH0668, wGH0669) were used in this comparison. The filtered evidence.txt file produced after removal of contaminant and reverse matches and PIF (*>* 0.5), mean reporter ion ratio between the single-cell samples and the carrier (*≤* 0.1), and FDR (*≤* 1%) filtration was used as the data source for this figure. The distribution of precursor intensities for all PSMs were divided into 30 intensity bins on a per-experiment basis, and the number of identifications per bin per experiment was calculated. The median and standard deviation of these per-bin identifications was calculated on a per-platform basis (shotgun or prioritization). The median and standard deviation were then represented as circles and error bars, respectively, on the associated figure.

#### Consistency heatmap

The precursors by single-cells matrix produced after PIF (*>* 0.5), mean reporter ion ratio between the single-cell samples and the carrier (*≤* 0.1), FDR (*≤* 1%), and CV (*≤* 0.4) filtration was sub-setted to contain only those precursors on the top tier of the consistency experiment inclusion list, which contained 1000 precursors identified in 50% or fewer of the shotgun experiments. The precursors were then aggregated to the peptide level, and their abundance measurements were binarized such that precursors with NA intensities for a single-cell sample were given a value of zero while precursors with detected reporter ion intensity were given a value of 1. The resulting binary dataframe was then presented as a heatmap with peptides on the y-axis and single cells on the x-axis to contrast data completeness for difficult-to-identify peptides placed on the top two tiers of a prioritized inclusion list. The set of peptides was filtered to only include entries with at least one cell with detected reporter ion intensity, as shown in Fig. 2c.

#### Peptide and protein-level data completeness contrast box plots

To produce the peptide-level data-completeness boxplot shown in Fig. 2d, the precursors by single-cells matrix produced after PIF (*>* 0.5), mean reporter ion ratio between the single-cell samples and the carrier (*≤* 0.1), FDR (*≤* 1%), and CV (*≤* 0.4) filtration was subsetted into two dataframes: one containing precursors from the top priority tier of the prioritized inclusion list and one containing precursors that were enabled for MS2 analysis from any priority tier. Precursor abundances were aggregated to the peptide level by summing the relative intensities observed for multiple charge states on a per-sample basis; this sum was not used for relative quantitation, only for representing whether detectable reporter ion signal was observed for a peptide in a single-cell sample, as opposed to an NA value. The fraction of peptides with detected reporter ion signal per single cell was computed for these two priority categories before being multiplied by 100 to produce the percent data completeness per cell.

To generate the protein-level data completeness box plot shown in Fig. 2d, the matrix of protein abundances by single cells produced by the scp pipeline^3, 32^ was subsetted to include only the proteins whose precursors were specified for MS2 analysis on the prioritized inclusion list. The fraction of these proteins with detected reporter ion signal per single cell was computed and then multiplied by 100 to produce the percent data completeness per cell.

#### HEK and melanoma PCA color-coded by median protein-set abundance

PCA was performed on the imputed, batch-corrected, and normalized proteins by cells matrix, using the prcomp function in R. For PCAs color-coded by the median protein set abundance, the median abundance of all proteins mapping to a protein set was calculated on a per-cell basis, using the unimputed batch-corrected data matrix. The vector of single-cell protein set abundances was then z-scored, and the resulting vector was joined with the vector of principal component coordinates (the scores vectors) by sample ID. The protein sets presented in this analysis were selected from the results of PC-weight-based Protein Set Enrichment Analysis (PSEA), as described below.

### Figure 3, HEK and melanoma samples

#### Reporter Ion Intensity Distributions

The evidence.txt file generated by MaxQuant was filtered to exclude PSMs with PIF *<* 0.5 and PEPs *≥* 0.02. Scans of type ‘MSMS’ were also removed, as these lack precursor mass and intensity information. The single-cell and control sample reporter ion intensities from each experiment were tallied across 30 intensity bins on the basis of whether they were associated with endogenous or spike-in proteins. These distributions were then plotted as side-by-side violin plots.

#### Regression of normalized reporter-ion intensities on spike-in concentrations

The evidence.txt file generated by MaxQuant was filtered to exclude PSMs with PIF *<* 0.5 and PEPs *≥* 0.02. Scans of type ‘MSMS’ were also removed, as these lack precursor mass and intensity information. Finally, only PSMs corresponding to the 4 sequences (AYFTAPSSER, VEVDSFS-GAK, TSIIGTIGPK, ELYEVDVLK) generated by tryptic digestion of the spike-in peptides were selected for comparing measured and spiked-in levels. On a per-experiment and per-precursor basis, the median reporter-ion intensity was taken of the 3 single-cell samples containing 1x concentrations of each spike-in peptide. All reporter-ion intensities for a given precursor were then divided by this median to generate relative reporter-ion intensities. As a given peptide sequence may be associated with multiple charge states, the per-sample normalized reporter-ion intensities were then condensed to a median value for each peptide sequence. The set of all median normalized reporter-ion intensities were then regressed on their respective spike-in amounts, setting the intercept to zero and using partial least squares. In the associated plot, the log_2_(Concentration) of zero corresponds to the 1x concentration, with each subsequent increment corresponding to the 2x, 4x, 8x, and 16x spike-in concentrations, respectively. The grey circle in the associated plot corresponds to the median normalized reporter-ion intensity for all spike-in peptides across all samples at the concentration level, while the error bars reflect the median +/- the standard deviation of the normalized reporter-ion intensities. The text at the top of the plot indicates the number of sequences observed across all 8 injections for that concentration. The data shown in this figure corresponds to a set of 8 prioritized experiments acquired via MaxQuant.live, in which the spike-in peptides were on the high priority tier and were allowed to fill for the default MS2 fill time of 300ms.

### Figures 4/5/6, BMDM samples

#### Protein set enrichment analysis (PSEA)

PSEA was performed using the vector of principal-component-associated protein weights produced by PCA analysis of the imputed, batch-corrected, and normalized cells x proteins matrix, generated from the prcomp function in R. The human gene set database was acquired from GOA^52^. The gene set database was filtered to remove entries corresponding to cellular components, in favor of entries annotated to molecular function and biological process. Gene-level annotation was used to map gene sets to the protein weights. The factor weights for all proteins matching a protein set were compared against the background distribution of protein weights using the two-tailed wilcoxon rank-based test of significance. The following filters were used to determine whether a statistical comparison was made: At least 5 proteins from a protein set must have been present in the data, at least 10% of the proteins within a protein set must have been present in the data, and protein sets must contain fewer than 200 entries. The median loading per protein set was then transformed to a z-score for interpretability. The p values were then converted to q values using the Benjamini and Hochberg approach, and results were filtered to 5% FDR. Single-condition PSEA: the PC-based PSEA performed in Fig. 4b was applied to each treatment group separately in Fig. 5a. For the PSEA performed on the HEK and melanoma samples shown in Fig. 2e, the minimum protein count was raised to 15, due to the higher protein coverage present in that set of experiments.

#### Endocytosis analysis, histogram, Figure 3b

The vectors of Dextran: AF568 MFI and event counts were retrieved from the Sony MA900 used to sort the dextran uptake subpopulations, and this data was filtered for MFIs greater than 1,000 and less than 50,000. The MFIs were then *log*_10_ transformed and plotted as a normalized histogram in Fig. 5b and for the LPS-treated samples and untreated samples, respectively.

#### Endocytosis analysis, volcano plot, Figure 3b

Differentially abundant proteins between the low and high-dextran uptake samples were identified via the DIA differential protein analysis script introduced earlier, and the results were plotted, such that proteins with *|*log_2_(FC)*| >* 3 were annotated. The volcano plots associated with the LPS-treated samples and untreated samples are presented in Fig. 5c and Extended Data Fig. 10, respectively.

#### Endocytosis analysis, PCA color-coded by endocytic proteins, Figure 3c

For each treatment condition (24hr LPS-treated or untreated), the set of proteins with statistically significant fold changes between the high and low dextran-uptake conditions (*|*log_2_(FC)*| ≥* 1; fold-change q-value *≤* 0.01) was intersected with the set of quantified proteins for the respective set of single-cell samples. The median abundance per cell was calculated for the sets of proteins associated with low dextran uptake or high dextran uptake, each vector of median abundances was then z-scored and extreme values capped at a z-score of *±*2, and the single-condition PCAs were color-coded by these z-scored protein abundances. The figures associated with the LPS-treated samples and untreated samples are presented in Fig. 5c and Extended Data Fig. 10, respectively.

#### Validation of MEROPS peptide quantitation, Figure 4a

MEROPS substrates which were n-terminally labeled with TMTpro in the bulk discovery experiments and whose cross-condition fold change was comparable between the bulk and single-cell experiments were taken to be validated measurements.

To benchmark the relative quantitation between the bulk discovery and single-cell samples, the treatment-condition-associated bulk samples detailed in the “MEROPS Bulk validation experiments, BMDMs” section, above, were filtered to an elution group PEP *≤* 0.01, and entries mapping to the same precursor species were condensed such that the observation with the highest intensity was taken to be representative. The LPS-treated and untreated samples were then joined by precursor, and the abundances column (sample) and row (precursor) normalized, by median and mean, respectively. The ratio of the normalized precursor abundances between the LPS-treated and untreated samples were then calculated. The matrix of batch-corrected unimputed protein abundances per single-cell from the prioritized BMDM analyses was then condensed to a representative abundance by treatment condition by taking the median protein abundance across all cells from a treatment group. The relative protein abundance ratio between treatment conditions was then computed, and the vector of fold changes was subset for the MEROPS cleavage products detected with TMT-Pro labeling of the neo-n-termini in the bulk experiments.

#### Biological annotation of MEROPS peptides, Figure 4b

Using the MaxQuant evidence.txt output from the DDA analysis of the bulk TMTpro-labeled duplex sample containing 24-hour LPS-treated (127C) and untreated (128N) BMDM samples (wGH215.raw), proteins which were differentially abundant between treatment conditions were identified in the following way: search results were filtered to contain precursors with PEPs *≤* 0.02 and PIFs *>* 0.8, and reverse matches and contaminants were filtered out; the reporter ion intensities for the two samples were column and row normalized by their means; the per-protein distributions of relative precursor abundances for each sample were subjected to a two-sided wilcoxon rank sum test; p-values were FDR corrected via the Benjamini and Hochberg approach and filtered to 1% FDR; the median relative abundance of the precursors mapping to a given protein were taken to reflect the relative abundance of that protein; differential proteins whose relative abundance ratio (LPS-treated:Untreated) was *>* 1 were annotated as marker proteins associated with LPS-treatment, or the untreated condition, otherwise.

A second set of marker proteins associated with pro-inflammatory M1-like macrophages or anti-inflammatory M2-like macrophages determined by transcriptomic analysis of monocytes, intermediate macrophages, fully differentiated macrophages, classically activated macrophages, and alternatively activated macrophages was also used in this analysis^41^. Genes with a log_2_ M1-to-M2 ratio greater than zero in the publication-associated database were annotated to be M1-associated, while genes with an M1-to-M2 ratio less than zero were annotated to be M2-associated.

Actin L104 cleaved by cathepsin (fragment 1), citrate synthase H26 cleaved by cathepsin E (fragment 2), and Actin L288 cleaved by Cathepsin D (fragments 1 and 2) which had been validated via bulk analysis were then tested for significant associations with either the treatment-condition-associated protein panels or the M1/M2-associated protein panels using a permutation test.

The matrix of single cells by batch-corrected unimputed protein abundances was filtered to contain the four MEROPS cleavage products and proteins annotated to the supplied list of marker proteins (either treatment-condition specific or macrophage-polarization specific), and a protein-protein correlation matrix was produced from this filtered matrix. The median correlation was then calculated between each MEROPS cleavage product and the set of proteins annotated to either of the two reference conditions (LPS-treated or untreated; M2 or M1), and the difference between the median correlation associated with each treatment condition was recorded.

The same procedure was then repeated 10,000 times, permuting the column names of the cells by proteins matrix each time. The p-value of the original correlation distance was subsequently determined to be the fraction of times a correlation distance as extreme as the one initially observed was generated by chance alone. The set of p-values was then FDR corrected using the Benjamini and Hochberg approach. If the original p value was zero, meaning no value generated by chance was as extreme as the initially observed value, then a q-value of 10*^−^*^5^ was used.

## Extended Data Figures

### Fraction of inclusion-list precursors detected and analyzed, Extended Data Figure 1

The MaxQuant.Live log files associated with the comparison of MaxQuant.Live with and without prioritization Fig. 1b/c were imported into the R environment and the lists of precursors detected by MaxQuant.live during the survey scan and subsequently sent for MS2 were extracted from the log files. The unique numeric precursor id was then matched to the associated inclusion list for each experiment to generate Extended Data Fig. 1. The precursors detected during the survey scan in each experiment are shown in Extended Data Fig. 1a, while the precursors sent for MS2 analysis in each experiment are shown in Extended Data Fig. 1b.

### Quantification variability across single-cell and control samples, Extended Data Figure 2

Within the single-cell pipeline^3, 32^, the CV (i.e. the standard deviation scaled by the mean) was computed for the relative abundances of all filtered precursors that mapped to a given leading razor protein on a per-sample basis (precursor-filtration metrics discussed in detail in the data filtration and normalization sections). The mean protein-level CV was then calculated on a per-sample basis, and a CV threshold was chosen which well separated the control samples from the single-cell samples. The distribution of CVs for single-cell and control samples associated with Fig. 2b are shown in Fig. 2a; the distribution of CVs for single-cell and control samples associated with Fig. 2c/d are shown in Fig. 2b; the distribution of CVs for single-cell and control samples associated with Fig. 4, Fig. 5, and Fig. 6 are shown in Extended Data Fig. 2a

### Peptide Properties and ID rates, Extended Data Figure 3

The shotgun and prioritized search results for the technical consistency experiments Fig. 2c/d were imported into the R environment, and the set of precursors not identified at 1% FDR in the prioritized analyses was determined. The median spectral confidence of identification and number of matching fragments for each of the precursors in this set was then calculated across all shotgun experiments using the evidence.txt and msms.txt files, respectively, and plotted in Extended Data Fig. 3.

### MaxQuant iMBR and pSCoPE contrast, Extended Data Figure 4

The 8 shotgun and 8 prioritized sample analyses associated with Figure 2a were selected for this contrast. While the search parameters associated with the prioritized sample analyses were unchanged, the 8 shotgun analyses were searched using the same search parameters as indicated previously in section FixRef, except iMBR was enabled. The set of identifications generated from the pSCoPE experiments was filtered at 1% FDR, and reverse matches and contaminants were removed. To generate the boxplots in panel a, the number of PSMs per experiment of type “MULTI-MATCH” and “MULTI-MATCH-MSMS” were summed to generate the number of MBR matches per experiment in the “All Precursors” facet. Only the PSMs of type “MULTI-MATCH-MSMS” were used to generate the number of MBR matches per experiment in the “Precursors with MS2 Scans” facet. To generate the boxplots in panel b, the following data handling steps were conducted: for the shotgun experiments, the set of all PSMs at 1% FDR was selected, all reverse matches and contaminants were removed, and all forward sequences identified via iMBR were added to this data frame; the evidence file for the prioritized analyses were filtered at 1% FDR; The precursors identified via iMBR in the shotgun runs were intersected with the prioritized inclusion list, and this group of peptides was then subset from both the shotgun and prioritized data sets; the number of experiments that each of these precursors was identified in was then tallied and faceted by priority level.

### Fraction of inclusion-list precursors detected and analyzed, Extended Data Figure 5

The MaxQuant.Live log files associated with the technical coverage and consistency experiments Fig. 2a/b/c/d/e were imported into the R environment and the lists of precursors detected by MaxQuant.live during the survey scan and subsequently sent for MS2 were extracted from the log files. The unique numeric precursor id was then matched to the associated inclusion list for each experiment to generate Extended Data Fig. 5. The MaxQuant.Live log statistics associated with Fig. 2a are shown in Extended Data Fig. 5a, while those associated with Fig. 2b/c/d/e are shown in Extended Data Fig. 5b.

### BMDM technical comparison, Extended Data Figure 6

The matrix of precursor abundances by single-cell samples for 20 shotgun analyses and 20 pSCoPE analyses was condensed to the peptide level by summing the relative intensities across charge states on a sample-specific basis. This was conducted as a means to determine peptides without detectable reporter ion signal in a given sample. The inclusion list was also condensed to the peptide level by associating a peptide sequence with the highest priority tier it appeared in. The fraction of peptides with detected reporter ion signal was calculated on a per sample and per priority-tier basis and displayed in Fig. 6a.

To calculate the percent data completeness on a per-protein level, the same procedure was followed for condensing precursors to proteins and for associating proteins with priority tiers. To calculate the number of peptides with detectable reporter ion signal per single cell, the matrix of precursor abundances by single-cell samples for 20 shotgun analyses and 20 pSCoPE analyses was condensed to the peptide level as performed previously, and the number of non-NA values was tallied on a per-single-cell basis. The number of proteins with detectable reporter ion signal per single cell was calculated from the matrix of unimputed protein abundances per cell. These tallies are displayed in Extended Data Fig. 6b.

To generate the histogram of representative precursor abundances and their corresponding fill times, the precursor intensities for all precursors identified in the Shotgun and pSCoPE single-cell BMDM analyses were split into tertiles. If a top tier precursor appeared in the bottom intensity tertile, it was allotted an MS2 fill time of 1000ms; if a top tier precursor appeared in the middle intensity tertile, it was allotted an MS2 fill time of 750ms; if a top-tier precursor appeared in the top intensity tertile, it was allotted an MS2 fill time of 500ms.

The matrix of precursor abundances by single-cell samples for 20 shotgun analyses and 20 pSCoPE analyses was subsetted to contain only those precursors that were allotted fill times of 750ms and 1000ms in the pSCoPE analyses. The percent data completeness was then calculated on a per-single-cell basis for the set of precursors allotted longer fill times present in the filtered matrix of precursors by single-cell samples. The results from this analyses are presented in Extended Data Fig. 6c.

### PCA from unimputed protein-level data, Extended Data Figure 7

In order to assess whether the qualitative trends observed in the cross-condition PCA or the PSEA based on the PCA-derived protein weight vectors were compromised by imputation, PCA was performed on the correlation matrix generated by the batch-corrected, unimputed cells x proteins matrix, and the resulting PCA plot was color-coded by cell type or the median relative abundances of the proteins corresponding to Type I IFN signaling or phagosome maturation, Extended Data Fig. 7.

### PCA color-coded by precursor-level data completeness, Extended Data Figure 8

In order to assess whether the sample separation observed in the cross-condition PCA (shown in Fig. 4) was driven by missing data, the single-cell data points were color-coded by the data completeness percentage. The post-data-filtration matrix of precursors by single-cells (filtration metrics described in the data processing and normalization section) was used to calculate the data completeness on a per-sample basis. The fraction of filtered precursors with observed reporter ion signal relative to the total number of filtered precursors detected across all experiments was then multiplied by 100 to generate the data completeness percentage.

### FACS gating parameters and staining controls, Extended Data Figure 9

FSC-A, SSC-A, and Dextran:PE-Texas Red FACS gate settings and sorting parameters were directly acquired from the Sony MA900 FACS instrument. Additional detail regarding the associated experimental parameters can be found in the “BMDM Endocytosis Assay” subsection of the Methods.

### Endocytosis Panel for Untreated BMDMs, Extended Data Figure 10

This plot was constructed in the same manner as the main text figure corresponding to the 24-hr LPS-treated samples.

## Extended Data Figures

**Extended Data Fig. 1.**
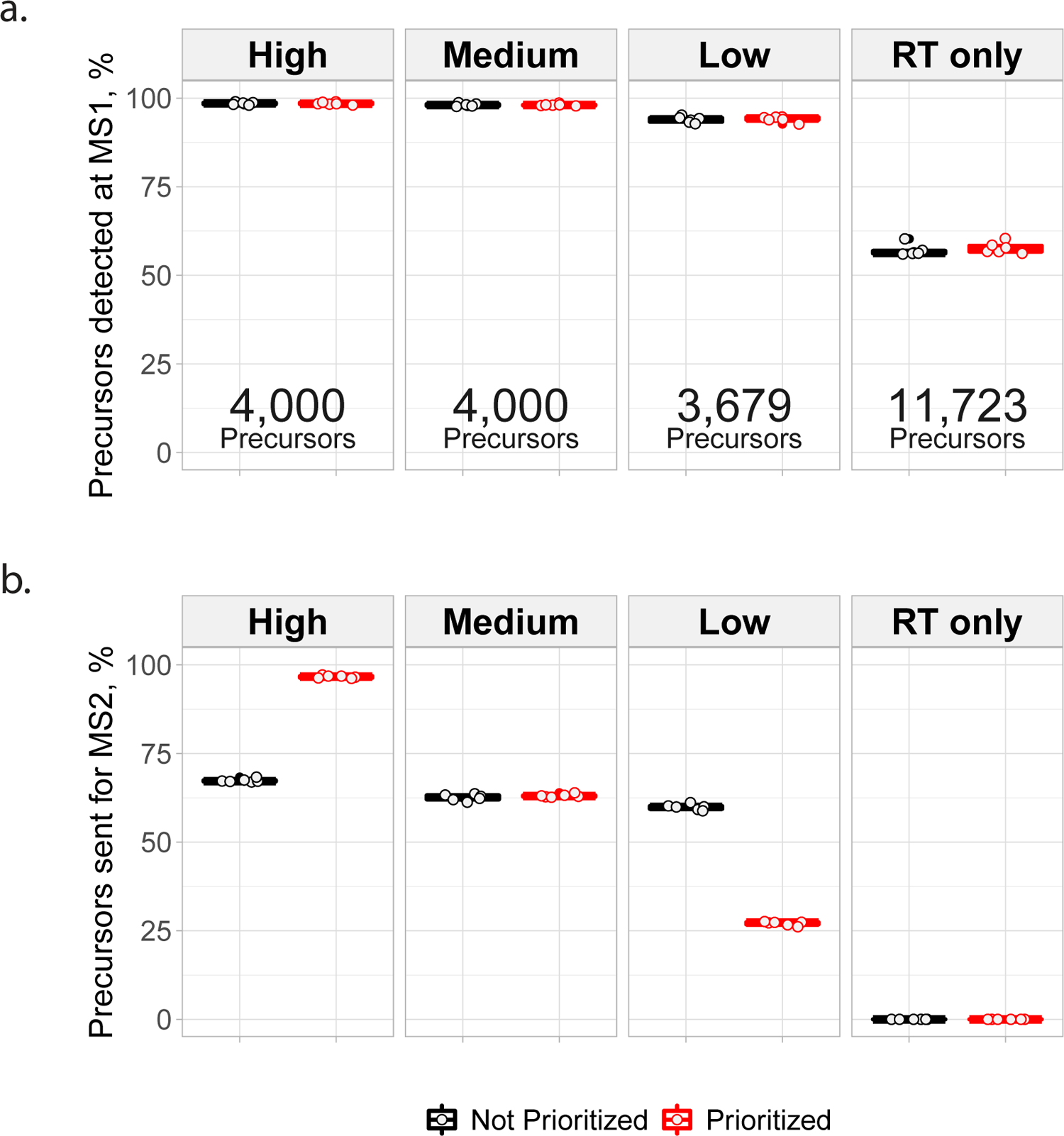
Percent of inclusion-list precursors detected and analyzed in platform benchmark runs, for MaxQuant.Live with and without prioritization enabled. (**a**) MS1 detection rates for precursors in the platform benchmark experiments displayed in Fig. 1a,b. Data collected using MaxQuant.Live in default mode are shown in black, while data collected using prioritization are shown in red. The precursor count displayed at the bottom of each priority level’s facet corresponds to the number of precursors present on the priority level of the inclusion list. (**b**) MS2 analysis rates for precursors in the platform benchmark experiments displayed in Fig. 1a,b. While the MS1 precursor detection rates are similar for both platforms, the MS2 analysis rates are correlated to the priority levels for prioritized analysis, but not for default MaxQuant.Live analyses. Each boxplot shown above contains 6 data points, one for each LC-MS/MS analysis.

**Extended Data Fig. 2.**
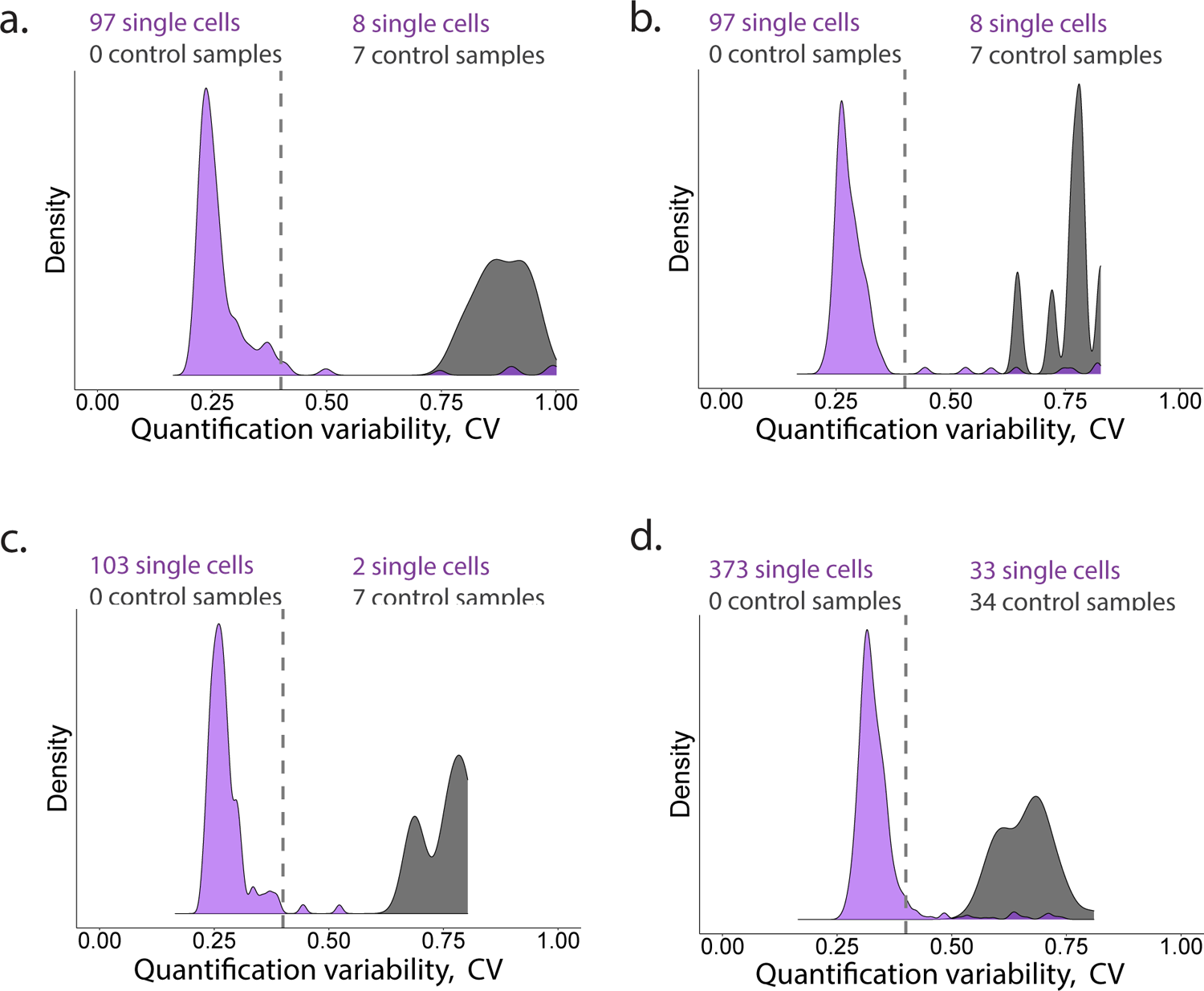
Single-cell quality controls The median coefficient of variation (i.e. the standard deviation scaled by the mean) of all peptide-level relative abundances that map to the same leading razor protein is used to separate successfully prepared single cells from those that will not generate accurate data. By choosing a CV threshold that separates control samples (droplets which received all reagents but did not contain a single cell) from single cells, cells with noisier protein-level quantitation can be removed prior to further data processing. The single cell and control tallies above each figure represent the number of single cells or control wells that passed the CV threshold of 0.4. (**a**) contains the CV distributions for the single-cell samples associated with Fig. 2a,b,c,d,e, analyzed by shotgun LC-MS/MS methods. (**b**) contains the CV distributions for the single-cell samples associated with Fig. 2a, analyzed by pSCoPE. (**c**) contains the CV distributions for the single-cell samples associated with Fig. 2b,c,d,e, analyzed by pSCoPE. (**d**) contains the CV distributions for the single-cell samples associated with Fig. 4, Fig. 5, and Fig. 6.

**Extended Data Fig. 3.**
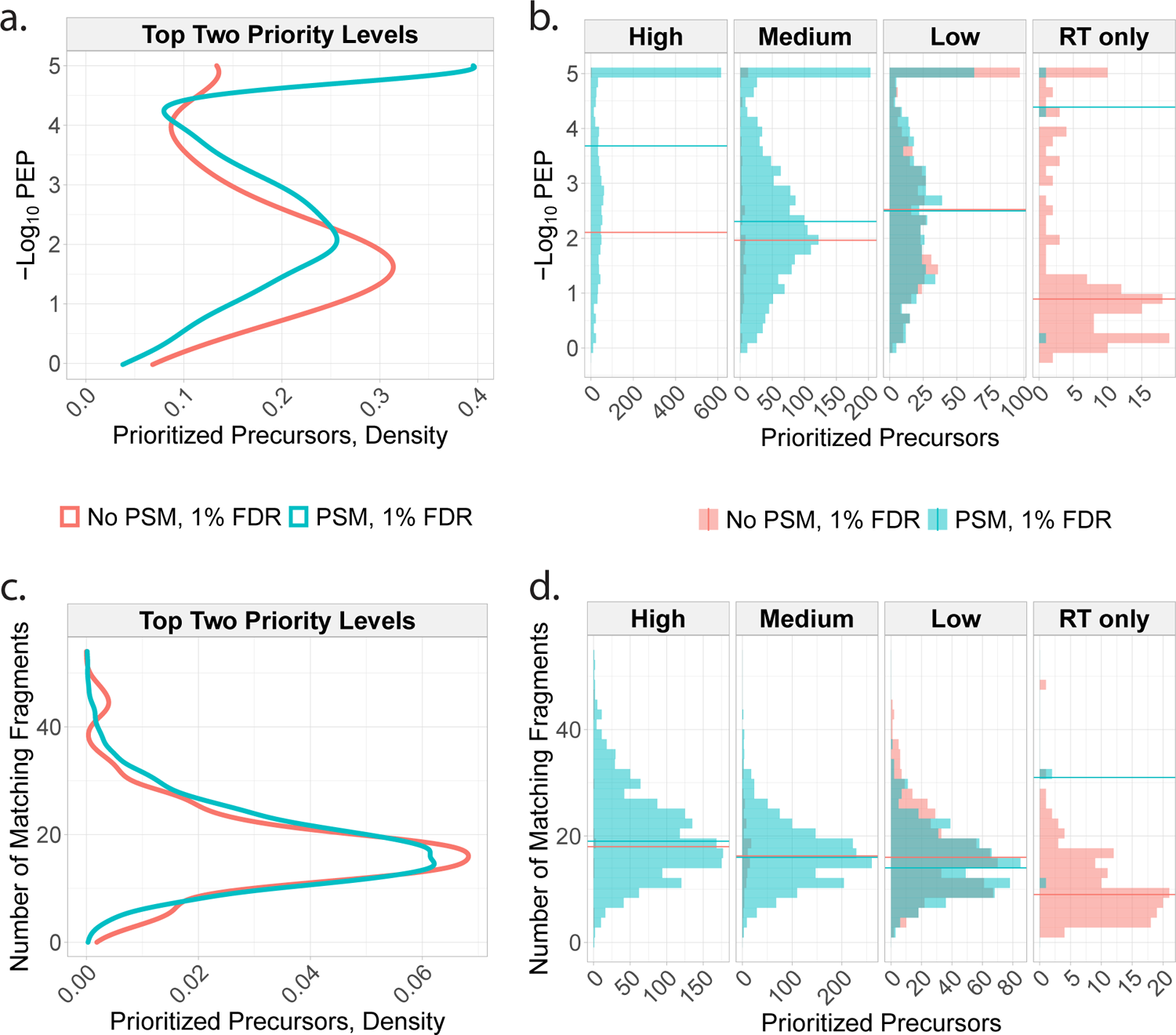
Properties of peptides successfully identified in pSCoPE runs The precursors from the inclusion list were split into those that resulted in confident PSMs and those that did not, and the properties of each set analyzed based on the shotgun runs used for making the inclusion lists. (**a**) Confidence of identification (quantified by the posterior error probability; PEP) and number of matching peptide fragments for successful and unsuccessful precursors. The data are shown for all prioritized peptides across all priority tiers. (**b**) The data from panel a are shown faceted by priority tier. All data shown are from the consistency experiments from Fig. 2a. In previous analyses conducted during a period of suboptimal instrument performance, the number of matching fragments was shown to effectively distinguish between the the peptides which were identified at 1% FDR and those that were not identified, which was reported in version 1 of our preprint^53^. This trend is not observed in the current dataset, which was acquired by the same instrument but with more efficient ion isolation by its quadruple.

**Extended Data Fig. 4.**
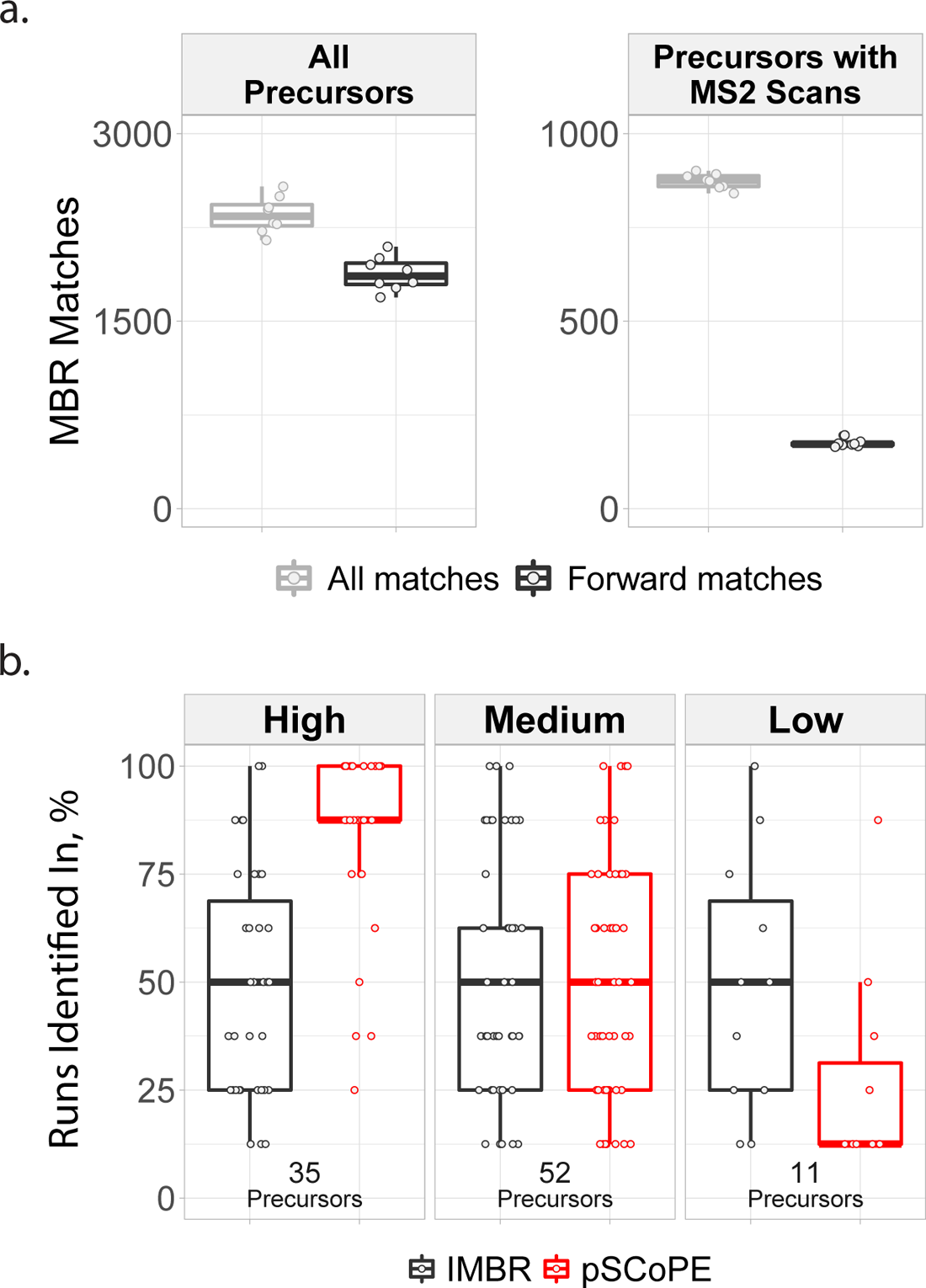
pSCoPE outperforms isobaric Match Between Runs (iMBR) for increasing consistency of identification across single-cell experiments. (**a**) The “All Precursors” facet heading indicates the total number of MBR-facilitated precursor identifications in each of 8 shotgun analyses. The “Precursors with MS2 Scans” facet heading indicates the total number of MBR-facilitated precursor identifications that are associated with MS2 scans, enabling reporter ion quantitation. In both facets, the identifications are segmented into “All matches”, a category which includes matches to reverse sequences, and “Forward matches”, which does not. Each point represents an experiment. Data derived from shotgun experiments shown in Fig. 2a,b,c,d,e. (**b**) The intersected precursors between the MBR-facilitated forward sequence matches and the corresponding prioritized analyses were then compared based on consistency of identification across the 8 experiments associated with each acquisition method. Each point represents a precursor. Data derived from shotgun and pSCoPE analyses shown in Fig. 2a.

**Extended Data Fig. 5.**
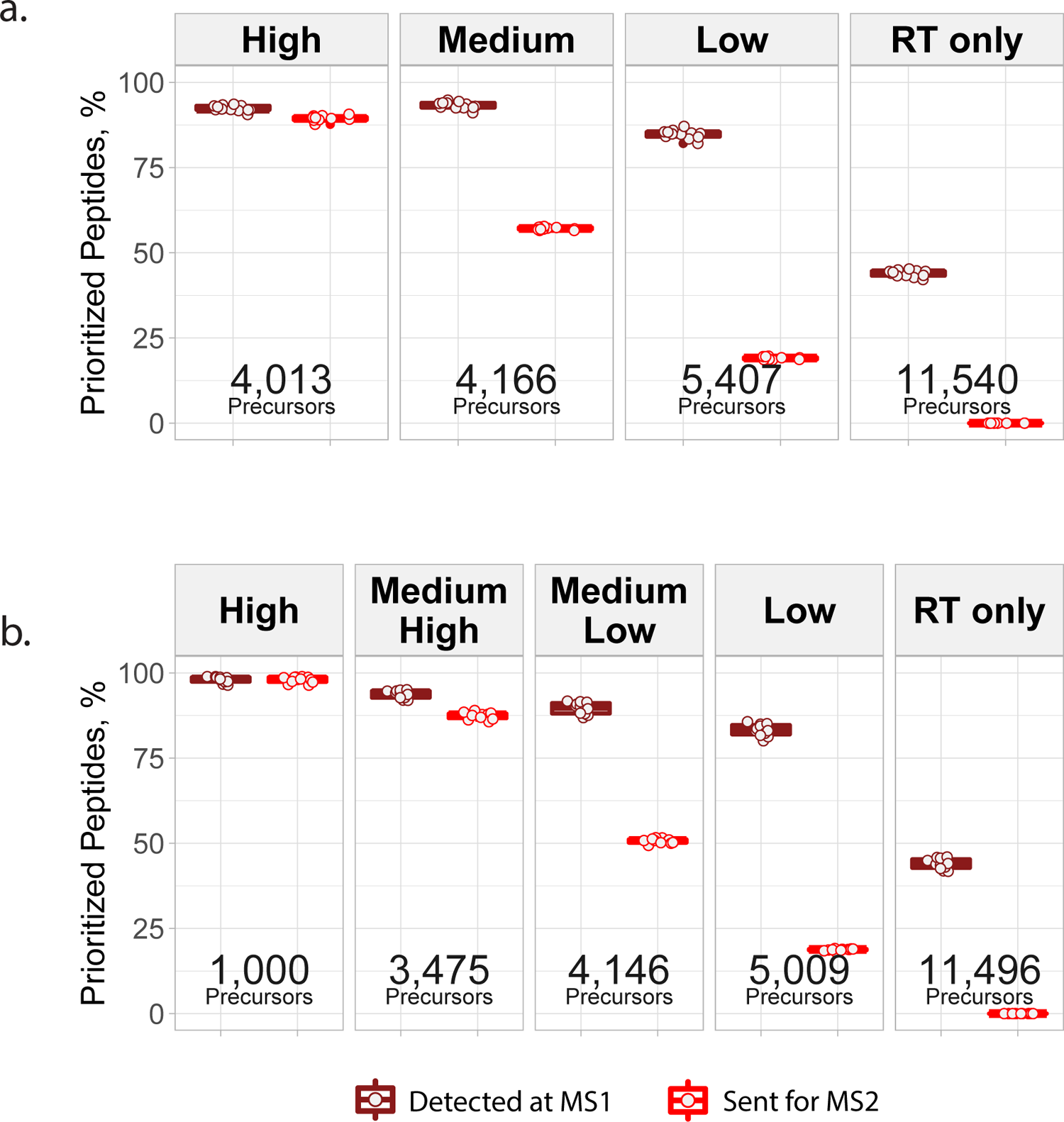
Fraction of inclusion-list precursors detected and analyzed in pSCoPE runs. (**a**) MS1 detection and MS2 analysis rates for prioritized precursors in the benchmark experiments displayed in Fig. 2a. Each boxplot contains 8 data points, one for each LC-MS/MS analysis. (**b**) MS1 detection and MS2 analysis rates for prioritized precursors in the benchmark experiments displayed in Fig. 2b,c,d,e. Each boxplot contains 8 data points, one for each LC-MS/MS analysis. In both panels, the statistics are shown for each tier along with the number of precursors in the tier.

**Extended Data Fig. 6.**
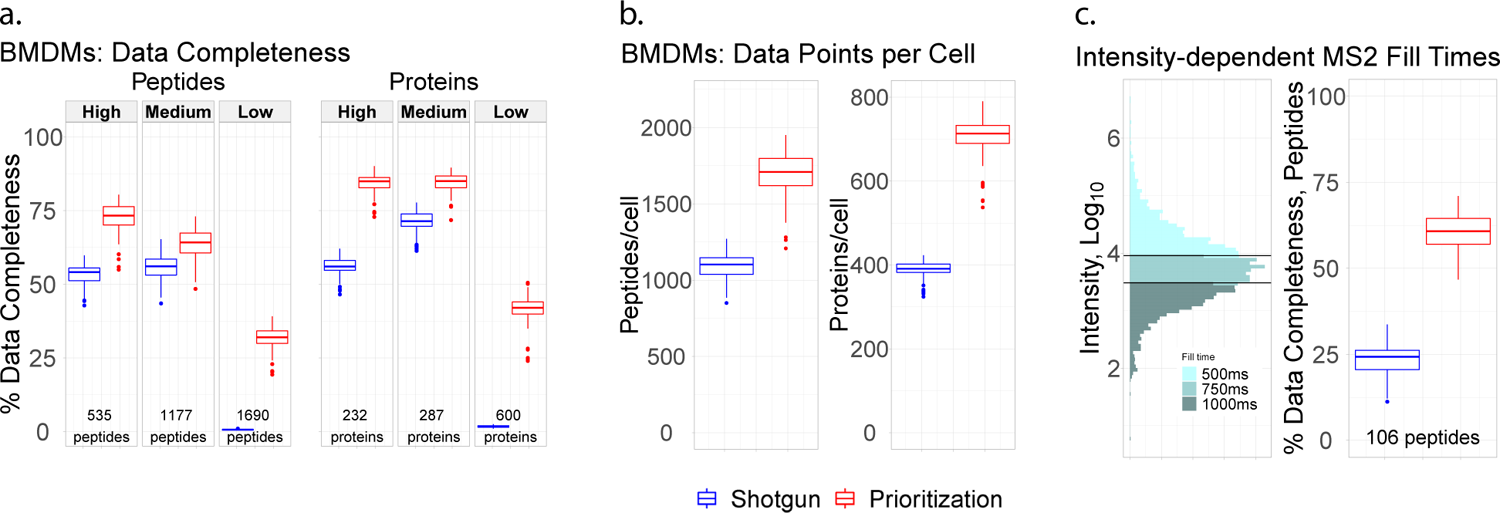
Data completeness and proteome coverage for single BMDM analysed by shotgun or prioritized methods. (**a**) Percent data completeness tallied for peptides and proteins quantified across twenty shotgun and twenty pSCoPE experiments, faceted by priority tier. (**b**) Number of peptides and proteins per single-cell sample across twenty shotgun and twenty pSCoPE experiments. (**c**) Illustration of precursor-intensity-dependent MS2 fill times for precursors on the top priority tier. Percent data completeness contrast for precursors which were allotted increased fill times in the pSCoPE analyses.

**Extended Data Fig. 7.**
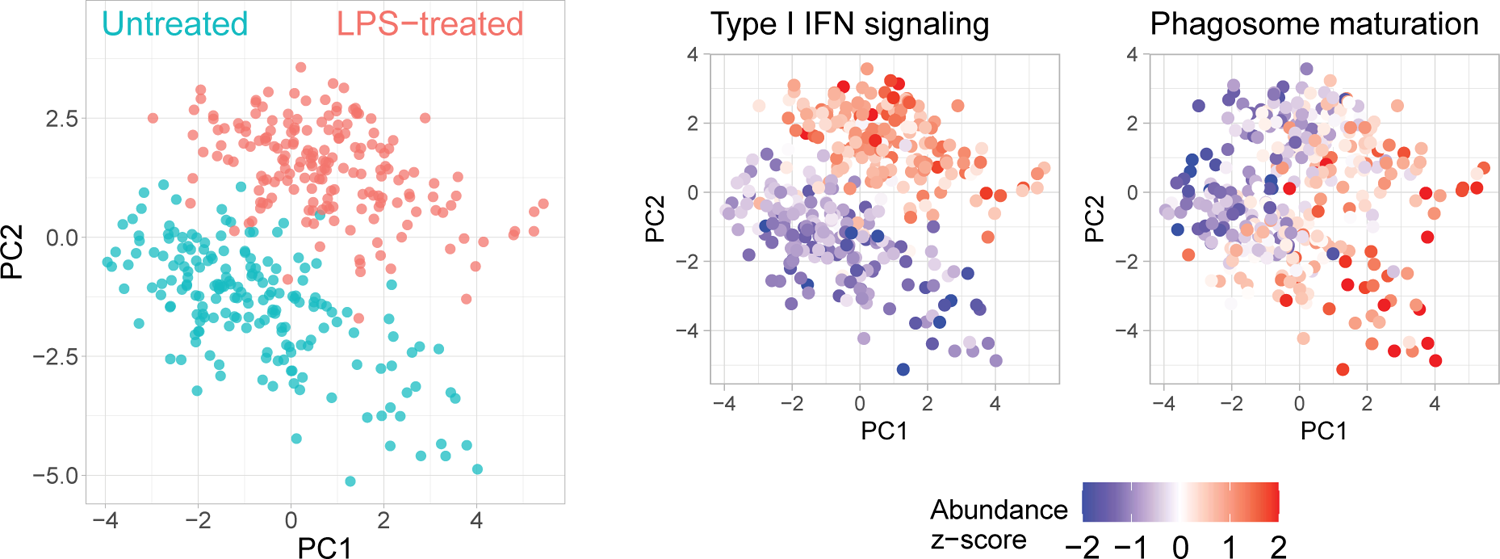
PCA of single BMDM using only observed data points To evaluate the robustness of our results to uncertainties stemming from missing data, we performed PCA of unimputed BMDM data. The single cells are color-coded by treatment condition, with adjoining PCA plots color-coded by the median relative abundance of proteins corresponding to type I interferon-mediated signaling and phagosome maturation.

**Extended Data Fig. 8.**
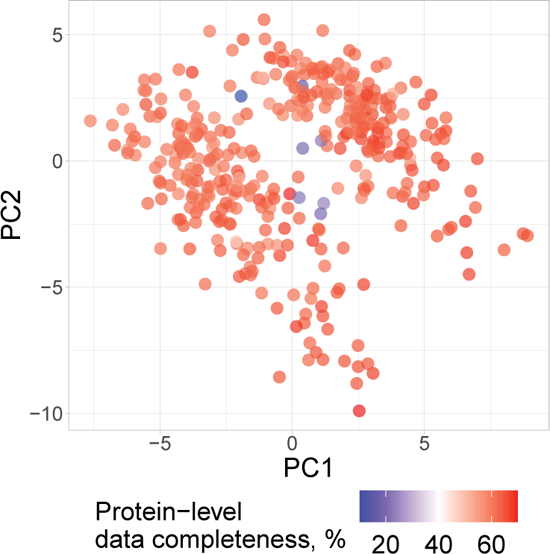
PCA color-coded by protein-level data completeness To evaluate the whether the biological conclusions we drew from our PC-weight-based PSEA could have been influenced by separation due to data completeness, we color-coded our cross-condition BMDM PCA by the percent data completeness on a per-cell basis.

**Extended Data Fig. 9.**
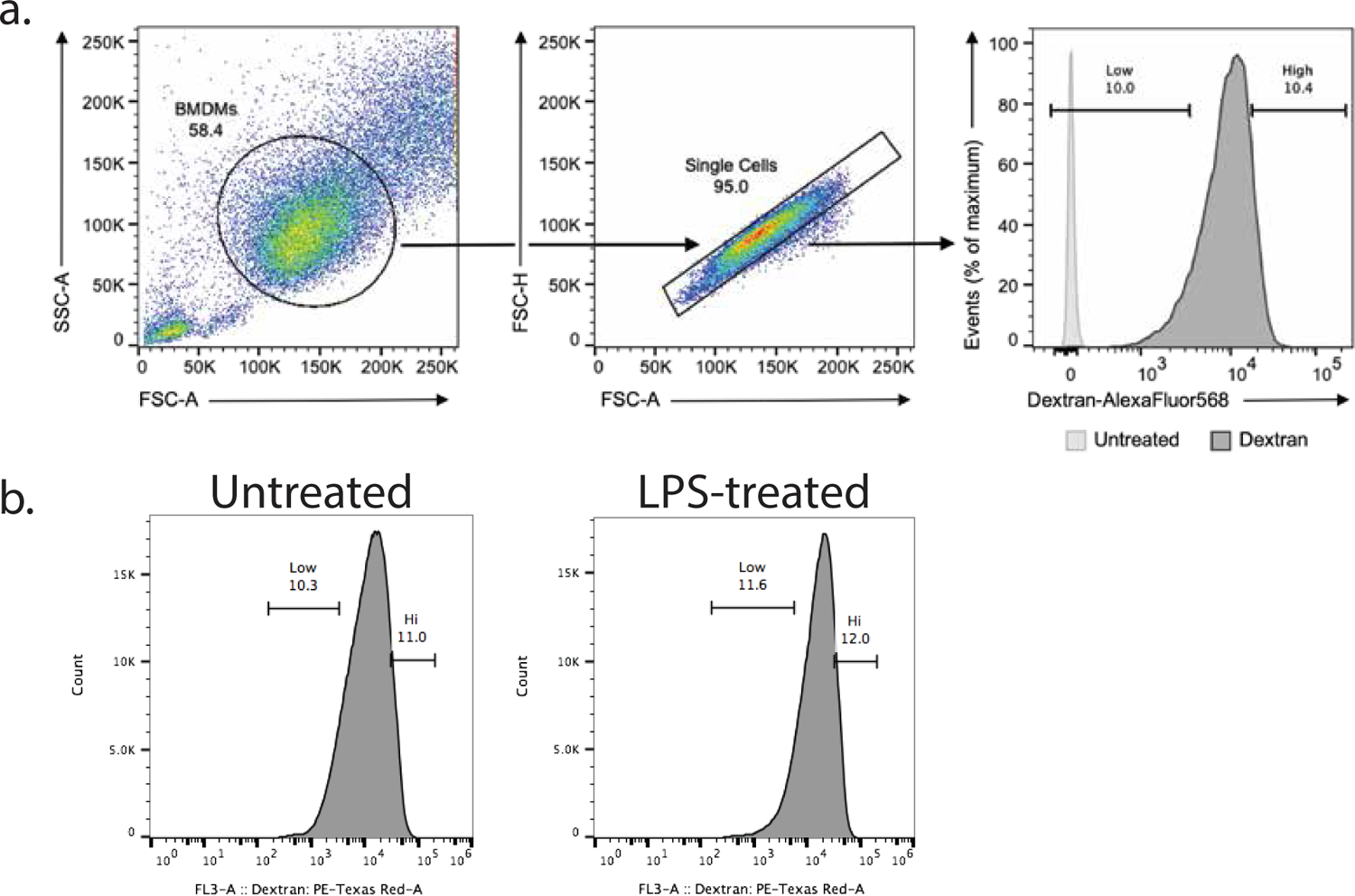
FACS gating parameters and staining controls. (**a**) FSC-A and SSC-A gates for sorted bone-marrow-derived macrophages and positive/negative staining populations. (**b**) Dextran:PE-Texas Red gating parameters for isolating the most and least endocytic BMDM populations from each treatment group (untreated and LPS-treated).

**Extended Data Fig. 10.**
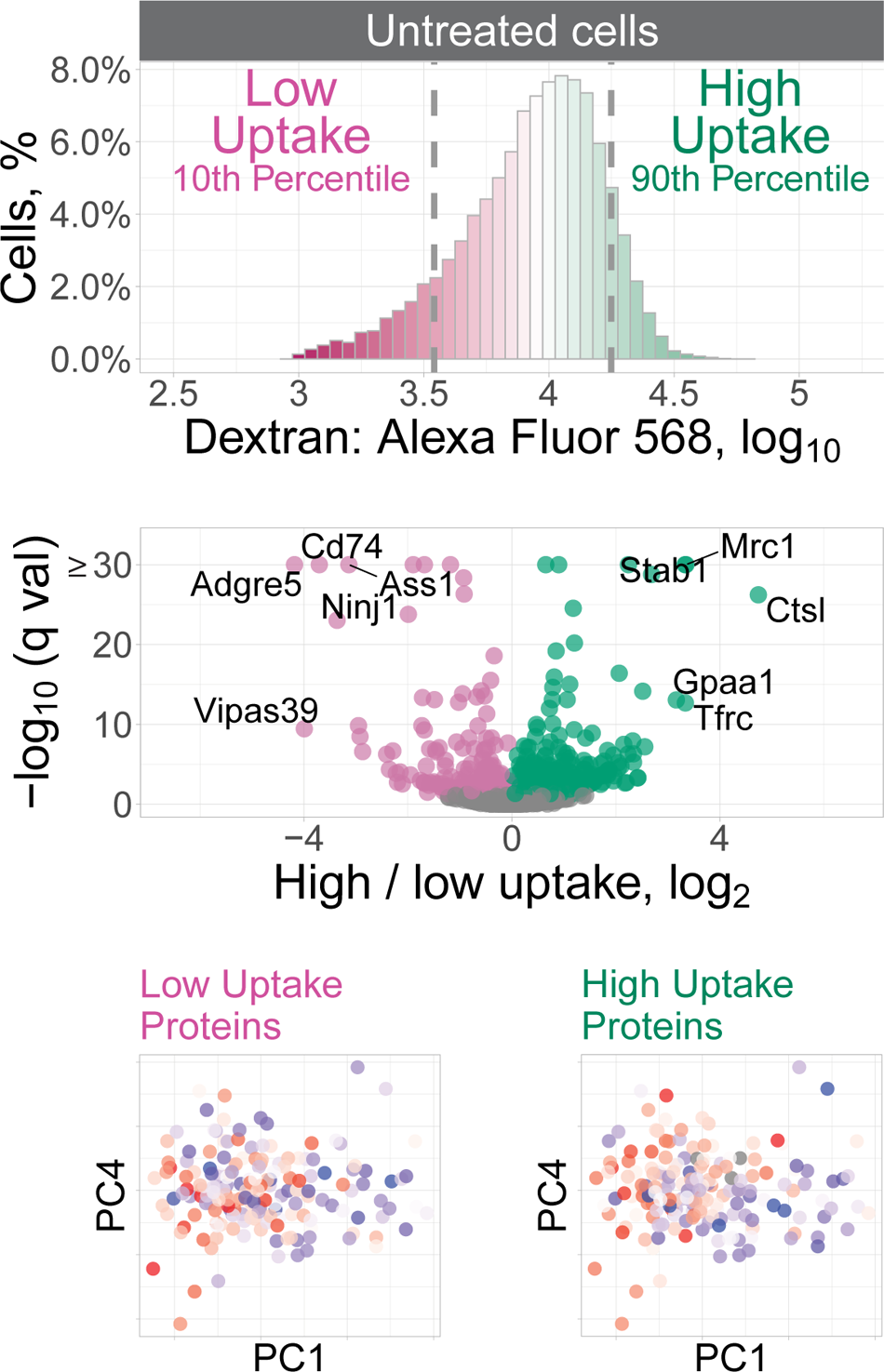
Dextran uptake in untreated BMDM samples The uptake of fluorescent dextran by the untreated macrophages was measured by FACS, and the cells with the lowest and highest uptake were isolated for protein analysis. The volcano plot displays the fold changes for deferentially abundant proteins and the associated statistical significance. The untreated macrophages were displayed in the space of their PCs and color-coded by the median abundance of the low-uptake or the high-uptake proteins. Both the low and the high-uptake proteins correlate inversely to PC1 (low-uptake: Spearman r = *−*0.29, *q ≤* 6 *×* 10^−4^; high-uptake: Spearman r = *−*0.37, *q ≤* 4 *×* 10^−6^).

## Supplemental Figures and Tables

**Supplemental Fig. S1.**
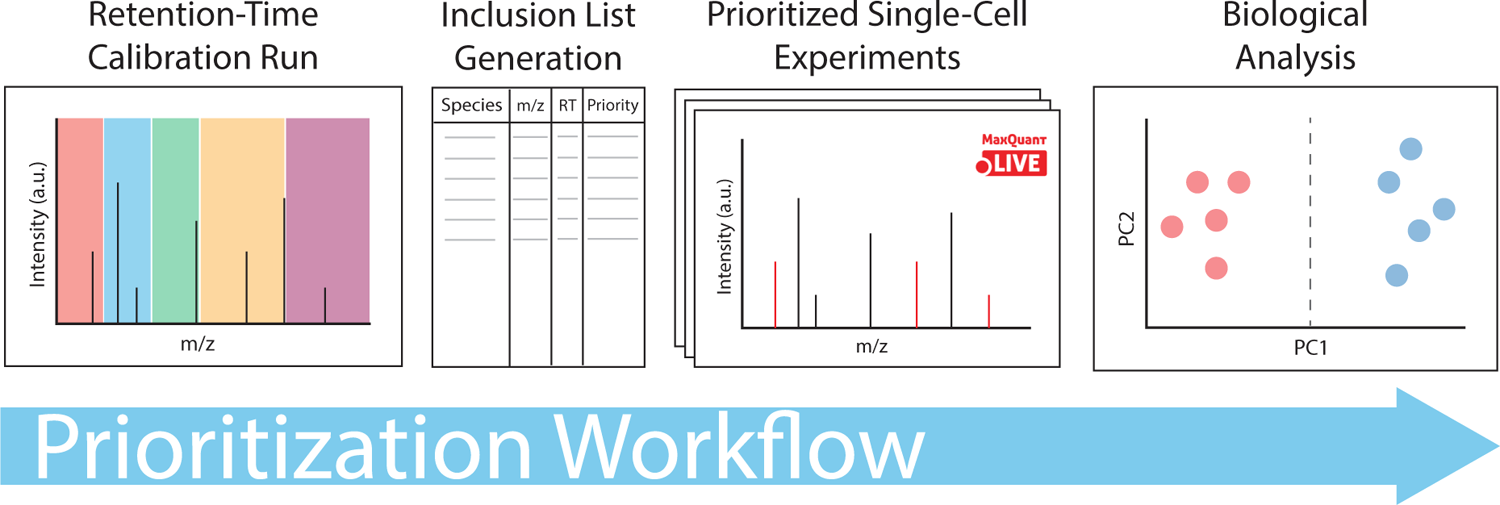
pSCoPE workflow schematic The pSCoPE workflow begins with the assembly of a number of precursors of experimental interest which can originate from prior DDA or DIA analyses of bulk samples, DDA analyses of previous SCoPE samples, or literature. A DIA method is then applied to a 1x injection of carrier and reference material to generate accurate retention times for the peptides of interest. The user then stratifies the peptides identified during the retention-time calibration run into priority tiers, with the top priority tier containing peptides of highest experimental interest. The prioritized inclusion list is imported into MaxQuant.Live for prioritized analysis of SCoPE samples.

---

**Supplemental Table S1.**
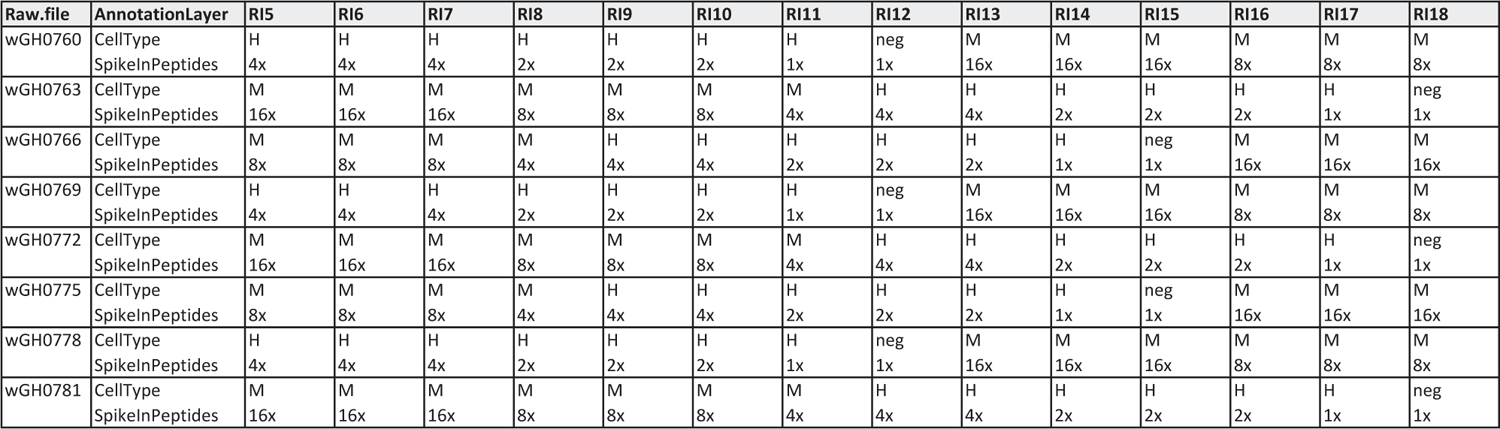
Experimental design for spike-in analyses. Schematic for the design of the single-cell sets used to benchmark reporter-ion quantitation in Fig. 3. Each row corresponds to a pSCoPE and having the indicated cell type and spike-in concentration for each sample. The sample column headers denote the TMTpro 18plex label associated with each sample, with RI5 indicating 128C, RI6 indicating 129N, and so on. In the “cell type” section for each experiment, “H” denotes HEK293 cells, “M” denotes melanoma cells, and “neg” denotes negative control samples, which were identically processed to the single-cell samples, but which did not contain a cell.

| Raw.file | AnnotationLayer | RI5 | RI6 | RI7 | RI8 | RI9 | RI10 | RI11 | RI12 | RI13 | RI14 | RI15 | RI16 | RI17 | RI18 |
| --- | --- | --- | --- | --- | --- | --- | --- | --- | --- | --- | --- | --- | --- | --- | --- |
| wGH0760 | CellType | H | H | H | H | H | H | H | neg | M | M | M | M | M | M |
|  | SpikeInPeptides | 4x | 4x | 4x | 2x | 2x | 2x | 1x | 1x | 16x | 16x | 16x | 8x | 8x | 8x |
| wGH0763 | CellType | M | M | M | M | M | M | M | H | H | H | H | H | H | neg |
|  | SpikeInPeptides | 16x | 16x | 16x | 8x | 8x | 8x | 4x | 4x | 4x | 2x | 2x | 2x | 1x | 1x |
| wGH0766 | CellType | M | M | M | M | H | H | H | H | H | H | neg | M | M | M |
|  | SpikeInPeptides | 8x | 8x | 8x | 4x | 4x | 4x | 2x | 2x | 2x | 1x | 1x | 16x | 16x | 16x |
| wGH0769 | CellType | H | H | H | H | H | H | H | neg | M | M | M | M | M | M |
|  | SpikeInPeptides | 4x | 4x | 4x | 2x | 2x | 2x | 1x | 1x | 16x | 16x | 16x | 8x | 8x | 8x |
| wGH0772 | CellType | M | M | M | M | M | M | M | H | H | H | H | H | H | neg |
|  | SpikeInPeptides | 16x | 16x | 16x | 8x | 8x | 8x | 4x | 4x | 4x | 2x | 2x | 2x | 1x | 1x |
| wGH0775 | CellType | M | M | M | M | H | H | H | H | H | H | neg | M | M | M |
|  | SpikeInPeptides | 8x | 8x | 8x | 4x | 4x | 4x | 2x | 2x | 2x | 1x | 1x | 16x | 16x | 16x |
| wGH0778 | CellType | H | H | H | H | H | H | H | neg | M | M | M | M | M | M |
|  | SpikeInPeptides | 4x | 4x | 4x | 2x | 2x | 2x | 1x | 1x | 16x | 16x | 16x | 8x | 8x | 8x |
| wGH0781 | CellType | M | M | M | M | M | M | M | H | H | H | H | H | H | neg |
|  | SpikeInPeptides | 16x | 16x | 16x | 8x | 8x | 8x | 4x | 4x | 4x | 2x | 2x | 2x | 1x | 1x |

**Supplemental Table S2.** **MS Methods for DIA Acquisitions:** Parameters for data acquisition for the DIA methods used in this publication. Accompanying DIA window schemes can be found in Supporting Table S4. Chromatographic methods used with these DIA methods can be found in Supporting Table S3. The text in black denotes parameters which are in common with the default method (Method 1), while the red text denotes parameters which vary from the default method.

| **MS1 Settings:** | **Method 1** | **Method 2** | **Method 3** | **Method 4** | **Method 5** |
| --- | --- | --- | --- | --- | --- |
| **Polarity:** | Positive | Positive | Positive | Positive | Positive |
| **Resolution:** | 140k | 140k | 140k | 70k | 140k |
| **AGC Target:** | 3e6 | 3e6 | 3e6 | 3e6 | 3e6 |
| **Max Injection Time:** | 512ms | 512ms | 512ms | 300ms | 512ms |
| **Scan Range (Th):** | 450-1600 | 450-1600 | 450-1258 | 478-1470 | 400-800 |
| **DIA Scan Settings:** |  |  |  |  |  |
| **Default Charge State** | 2 | 2 | 2 | 2 | 2 |
| **Polarity** | Positive | Positive | Positive | Positive | Positive |
| **Resolution** | 35k | 35k | 35k | 35k | 17.5k |
| **AGC target** | 5e5 | 5e5 | 5e5 | 3e6 | 5e5 |
| **Max Injection Time** | auto | auto | auto | 110ms | 64ms |
| **MSX Count** | 1 | 1 | 1 | 1 | 1 |
| **MSX isochronous** | on | on | on | on | on |
| **Fixed first mass** | 200 | 200 | 200 | 200 | 200 |
| **NCE** | 33 | 33 | 33 | 27 | 27 |
| **DIA window scheme** | 1 | 2 | 3 | 4 | 5 |

**Supplemental Table S3.** Gradient Methods for DIA Acquisitions: Chromatographic parameters for the DIA methods used in this publication. Accompanying instrument parameters can be found in Supporting Table S2.

| **Time:** | **Meth. 1** | **Time** | **Meth. 2** | **Time** | **Meth. 3** | **Time** | **Meth. 4** |
| --- | --- | --- | --- | --- | --- | --- | --- |
| 0 min. | 4% B | 0 min. | 4% B | 0 min. | 4% B | 0 min. | 4% B |
| 4 min. | 4% B | 4 min. | 4% B | 4 min. | 4% B | 4 min. | 4% B |
| 12 min. | 8% B | 12 min. | 8% B | 12 min. | 7% B | 22 min. | 8% B |
| 75 min. | 35% B | 75 min. | 35% B | 135 min. | 32% B | 130 min. | 29% B |
| 77 min. | 95% B | 77 min. | 95% B | 137 min. | 95% B | 134 min. | 95% B |
| 80.1 min. | 4% B | 80.1 min. | 4% B | 140.1 min. | 4% B | 140.1 min. | 4% B |
| 95 min. | 4% B | 105 min. | 4% B | 160 min. | 4% B | 155 min. | 4% B |
| end | 4% B | end | 4% B | end | 4% B | end | 4% B |

**Supplemental Table S4.** DIA window schemes for DIA acquisitions: This table contains DIA windows parameters referenced in the associated instrument methods detailed in Supporting Table S2 for the DIA methods used in this publication.

| **DIA Window Scheme** | **1** | **2** | **3** | **4** | **5** |
| --- | --- | --- | --- | --- | --- |
| **Window 1:** |  |  |  |  |  |
| **Loop Count:** | 21 | 21 | 21 | 25 | 20 |
| **Isolation Window (Th):** | 20 | 15 | 20 | 12.5 | 19.5 |
| **Spans (Th):** | 450-860 | 450-755 | 450-860 | 450-778.5 | 400-799.5 |
| **Window 2:** |  |  |  |  |  |
| **Loop Count:** | 7 | 8 | 8 | 7 |  |
| **Isolation Window (Th):** | 50 | 20 | 50 | 25 |  |
| **Spans (Th):** | 859.5-1206.5 | 754.5-911 | 859.5-1256 | 778 - 950 |  |
| **Window 3:** |  |  |  |  |  |
| **Loop Count:** | 1 | 6 |  | 8 |  |
| **Isolation Window (Th):** | 394 | 50 |  | 62.5 |  |
| **Spans (Th):** | 1206 - 1600 | 910.5-1208 |  | 949.5-1446 |  |
| **Window 4:** |  |  |  |  |  |
| **Loop Count:** |  | 1 |  |  |  |
| **Isolation Window (Th):** |  | 393 |  |  |  |
| **Spans (Th):** |  | 1207.5-1600.5 |  |  |  |

**Supplemental Table S5.** **MS methods for Shotgun Acquisitions:** Parameters for data acquisition for the shotgun methods used in this publication. Chromatographic methods used with these shotgun methods can be found in Supporting Table S6. The text in black denotes parameters which are in common with the default method (Method 1), while the red text denotes parameters which vary from the default method.

| **MS1 settings:** | **Method 1** | **Method 2** | **Method 3** | **Method 4** | **Method 5** |
| --- | --- | --- | --- | --- | --- |
| Resolution | 70k | 70k | 70k | 70k | 70k |
| AGC Target | 1e6 | 1e6 | 1e6 | 1e6 | 1e6 |
| Injection Time | 100ms | 100ms | 100ms | 100ms | 100ms |
| Scan Range (Th) | 450-1600 | 450-1600 | 450-1600 | 450-1258 | 450-1258 |
| **MS2 settings:** |  |  |  |  |  |
| Resolution | 70k | 35k | 70k | 140k | 140k |
| AGC Target | 5e4 | 5e4 | 5e4 | 5e4 |  |
| Injection Time | 300 ms | 150 ms | 300ms | 500 ms | 600 ms |
| TopN | 7 | 14 | 7 | 4 | 4 |
| Isolation Window (Th) | 0.7 | 0.7 | 0.5 | 0.5 | 0.5 |
| Offset (Th) | 0.3 | 0.3 | 0 | 0 | 0 |
| Fixed First Mass (Th) | 100 | 100 | 100 | 100 | 100 |
| NCE | 33 | 33 | 33 | 33 | 33 |
| Spectrum Type | Centroid | Centroid | Centroid | Centroid | Centroid |
| Min. AGC Target | 2e4 | 2e4 | 2e4 | 2e4 | 2e4 |
| Apex Triggering | FALSE | FALSE | FALSE | FALSE | FALSE |
| Charge Exclusion | Unknown, 1, and > 6 | Unknown, 1, and > 6 | Unknown, 1, and > 6 | Unknown, 1, and > 6 | Unknown, 1, and > 6 |
| Peptide Match | FALSE | FALSE | FALSE | FALSE | FALSE |
| Exclude Isotopes | Enabled | Enabled | Enabled | Enabled | Enabled |
| Dynamic Exclusion (sec) | 30 | 30 | 30 | 30 | 30 |

**Supplemental Table S6.** Gradient Methods for Shotgun Acquisitions: Chromatographic parameters for the shotgun methods used in this publication. Accompanying instrument parameters can be found in Supporting Table S5.

| **Time:** | **Methods 1** | **Time** | **Method 2** |
| --- | --- | --- | --- |
| 0 min. | 4% B | 0 min. | 4% B |
| 4 min. | 4% B | 4 min. | 4% B |
| 12 min. | 8% B | 12 min. | 8% B |
| 75 min. | 35% B | 75 min. | 35% B |
| 77 min. | 95% B | 77 min. | 95% B |
| 80.1 min. | 4% B | 80.1 min. | 4% B |
| 95 min. | 4% B | 105 min. | 4% B |
| end | 4% B | end | 4% B |

**Supplemental Table S7.** Method and Sample Mapping: MaxQuant.Live Contrast Experiments, Fig. 1: This table connects the experiments performed as part of Figure 1 with information regarding their sample type, instrument method, gradient, and inclusion list. More details regarding the instrument methods (Supporting Table S2,Supporting Table S5,Supporting Table S10), gradients (Supporting Table S3 and Supporting Table S6), and inclusion lists can be found in their respective tables and sections.

| Purpose | Sample | MS Method | Gradient | Inclusion List | Figure |
| --- | --- | --- | --- | --- | --- |
| Spectral Library | SQC, 10x | DIA:1 | 1 |  |  |
| Spectral Library | SQC, 10x | DIA:2 | 1 |  |  |
| Spectral Library | SQC, 1x | DIA:1 | 1 |  |  |
| Spectral Library | SQC, 1x | DIA:1 | 1 |  |  |
| RT Calibration | SQC, 1x | DIA:1 | 1 |  |  |
| Scout | SQC, 1x | pSCoPE:1 | 1 | 1 |  |
| Scout | SQC, 1x | pSCoPE:1 | 1 | 2 |  |
| Scout | SQC, 1x | pSCoPE:1 | 1 | 3 |  |
| Scout | SQC, 1x | pSCoPE:1 | 1 | 4 |  |
| pSCoPE Benchmark | SQC, 1x | MaxQuant.Live:1 | 1 | 5 | 1 |
| pSCoPE Benchmark | SQC, 1x | pSCoPE:1 | 1 | 5 | 1 |

**Supplemental Table S8.** Method and Sample Mapping: HEK and Melanoma Experiments, Figs. 2/3: This table connects the experiments performed as part of Figures 2 and 3 with information regarding their sample type, instrument method, gradient, and inclusion list. More details regarding the instrument methods (Supporting Table S2, Supporting Table S5, and Supporting Table S11), gradients (Supporting Table S3 and Supporting Table S6), and inclusion lists can be found in their respective tables and sections.

| Purpose | Sample | MS Method | Gradient | Inclusion List | Figure |
| --- | --- | --- | --- | --- | --- |
| Spectral Library | Carrier and Ref., 10x | DIA:1 | 1 |  |  |
| Spectral Library | Carrier and Ref., 10x | DIA:2 | 1 |  |  |
| Spectral Library | Carrier and Ref., 1x | DIA:1 | 1 |  |  |
| Spectral Library | Carrier and Ref., 1x | DIA:1 | 1 |  |  |
| RT Calibration | Carrier and Ref., 1x | DIA:1 | 1 |  |  |
| Scout | Carrier and Ref., 1x | pSCoPE:2 | 1 | 6 |  |
| Scout | Carrier and Ref., 1x | pSCoPE:2 | 1 | 7 |  |
| Scout | Carrier and Ref., 1x | pSCoPE:2 | 1 | 8 |  |
| Scout | Carrier and Ref., 1x | pSCoPE:2 | 1 | 9 |  |
| pSCoPE Benchmark | Single cell (nPOP) | Shotgun:3 | 1 |  | 2 |
| pSCoPE Benchmark | Single cell (nPOP) | pSCoPE:2 | 1 | 10 and 11 | 2 |
| Spike-in Analysis | Single cell (nPOP) | pSCoPE:3 | 1 | 12 | 3 |

**Supplemental Table S9.** Method and Sample Mapping: BMDM samples, Figs. 4/5/6: This table connects the experiments performed as part of Figure 1 with information regarding their sample type, instrument method, gradient, and inclusion list. More details regarding the instrument methods (Supporting Table S2, Supporting Table S5, Supporting Table S12, and Supporting Table S13), gradients (Supporting Table S3 and Supporting Table S6), and inclusion lists can be found in their respective tables and sections.

| Purpose | Sample | MS Method | Gradient | Inclusion List | Figure |
| --- | --- | --- | --- | --- | --- |
| Differential Protein Analysis | 1,800 cells from each condition (LPS and control) | DDA:1 | 1 |  |  |
| Differential Protein Analysis | 1,800 cells from each condition (LPS and control) | DDA:2 | 1 |  |  |
| Differential Protein Analysis | 1,000 cells from each condition (LPS and control), separately | DIA:1 | 1 |  |  |
| Endocytosis Assay | 10,000 cells from each condition (LPS and control), separately | DIA:5 | 4 |  | 5b |
| MEROPS Validation | 1μg from each condition (LPS and control), separately | DIA:4 | 3 | | 6a |
| Spectral Library | Carrier and Ref., 5x | DIA:3 | 1 |  |  |
| Spectral Library | Carrier and Ref., 1x | DIA:3 | 1 |  |  |
| RT Calibration | Carrier and Ref., 1x | DIA:3 | 1 |  |  |
| Scout | Carrier and Ref., 1x | pSCoPE:4 | 1 | 13 |  |
| pSCoPE Benchmark | Single cell (nPOP) | Shotgun:4 | 1 |  |  |
| pSCoPE Benchmark | Single cell (nPOP) | pSCoPE:5 | 1 | 14 | 4,5,6 |
| Method Development | Single cell (mPOP) | Shotgun:5 | 2 |  |  |
| Method Development | Single cell (mPOP) | pSCoPE:6 | 2 | 15 |  |

**Supplemental Table S10.** Prioritized acquisition parameters for benchmarking MaxQuant.Live with and without prioritization, Fig. 1 This table contains the MaxQuant.Live parameters used in experiments associated with Figure 1. The experiment-type-to-method mapping for these methods can be found in the corresponding table (Supporting Table S7).

|  |  |
| --- | --- |
| **Global settings: Survey scan** | **Scout Runs and Fig. 1 (Method 1)** |
| ScanDataAsProfile | True |
| PositiveMode | True |
| MaxIT | 100ms |
| Resolution | 70,000 |
| AgcTarget | 1,000,000 |
| MzRange | (450,1600) |
| BoxCarScans | 0 |
| **Global settings: TopN** |  |
| NumOfMS2Scans | 0 |
| **RealtimeCorrection** | **Scout Runs and Fig. 1 (Method 1)** |
| MzTolerances | (4.5,5) |
| RetentionTimeTolerances | (0.01,2) |
| SigmaScaleFactorRt | 3 |
| PeptideHistoryLength | 2 |
| MinUsedCorrectionPeptides | 50 |
| IntensityPeakRatioThreshold | 1e-5 |
| PeptideDetectionIsoPeaks | 2 |
| IsotopeTolerance | 9 |
| Ms2DetectionNeeded | False |
| Ms2ExcludeDetectedPeptides | False |
| Ms2MinNormIntensity | 0.1 |
| Ms2MzTolerance | 20 |
| PIFWindowSize | 1.4 |
| **TargetedMs2** | **Scout Runs and Fig. 1 (Method 1)** |
| BatMode | False |
| AutoPriority | True |
| DefaultPriority | 0 |
| MaxNumOfScans | 1 |
| WindowAndOffsetInDalton | False |
| ScanDataAsProfile | False |
| WindowSize | 0.5 |
| MzOffset | 0 |
| LowerMzBound | 100 |
| CollisionEnergy | 33 |
| LifeTime | 2,400ms |
| Resolution | 70,000 |
| MaxIT | 300ms |
| AgcTarget | 3,000,000 |
| PositiveMode | True |

**Supplemental Table S11.** Prioritized acquisition parameters for HEK and melanoma single-cell sets, Fig. 2/3: This table contains the MaxQuant.Live parameters used in experiments associated with Figure 1. The experiment-type-to-method mapping for these methods can be found in the corresponding table (Supporting Table S8).

| **Global settings: Survey scan** | **Scout Runs and Fig. 2 (Method 2)** | **Fig. 3 (Method 3)** |
| --- | --- | --- |
| ScanDataAsProfile | True | True |
| PositiveMode | True | True |
| MaxIT | 100ms | 100ms |
| Resolution | 70,000 | 70,000 |
| AgcTarget | 1,000,000 | 1,000,000 |
| MzRange | (450,1600) | (450,1600) |
| BoxCarScans | 0 | 0 |
| **Global settings: TopN** |  |  |
| NumOfMS2Scans | 0 | 0 |
| **RealtimeCorrection** | **Scout Runs and Fig. 2 (Method 2)** | **Fig. 3 (Method 3)** |
| MzTolerances | (4.5,5) | (4.5,5) |
| RetentionTimeTolerances | (0.01,2) | (0.01,2) |
| SigmaScaleFactorRt | 3 | 3 |
| PeptideHistoryLength | 2 | 2 |
| MinUsedCorrectionPeptides | 50 | 50 |
| IntensityPeakRatioThreshold | 1e-5 | 1e-5 |
| PeptideDetectionIsoPeaks | 2 | 2 |
| IsotopeTolerance | 9 | 9 |
| Ms2DetectionNeeded | False | False |
| Ms2ExcludeDetectedPeptides | False | False |
| Ms2MinNormIntensity | 0.1 | 0.1 |
| Ms2MzTolerance | 20 | 20 |
| PIFWindowSize | 1.4 | 1.4 |
| **TargetedMs2** | **Scout Runs and Fig. 2 (Method 2)** | **Fig. 3 (Method 3)** |
| BatMode | False | False |
| AutoPriority | True | True |
| DefaultPriority | 0 | 0 |
| MaxNumOfScans | 1 | 1 |
| WindowAndOffsetInDalton | False | False |
| ScanDataAsProfile | False | False |
| WindowSize | 0.5 | 0.5 |
| MzOffset | 0 | 0 |
| LowerMzBound | 100 | 100 |
| CollisionEnergy | 33 | 33 |
| LifeTime | 2,100ms | 2,400ms |
| Resolution | 70,000 | 70,000 |
| MaxIT | 300ms | 300ms |
| AgcTarget | 3,000,000 | 3,000,000 |
| PositiveMode | True | True |

**Supplemental Table S12.** Prioritized acquisition parameters for nPOP BMDM single-cell samples, Fig. 4/5/6: This table contains the MaxQuant.Live parameters used in experiments associated with Figure 1. The experiment-type-to-method mapping for these methods can be found in the corresponding table (Supporting Table S9).

| **Global settings: Survey scan** | **Scout sample (Method 4)** | **Single-cell sets (Method 5)** |
| --- | --- | --- |
| ScanDataAsProfile | True | True |
| PositiveMode | True | True |
| MaxIT | 100ms | 100ms |
| Resolution | 70,000 | 70,000 |
| AgcTarget | 1,000,000 | 1,000,000 |
| MzRange | (450,1258) | (450,1258) |
| BoxCarScans | 0 | 0 |
| **Global settings: TopN** | **Scout sample (Method 4)** | **Single-cell sets (Method 5)** |
| NumOfMS2Scans | 0 | 0 |
| **RealtimeCorrection** | **Scout sample (Method 4)** | **Single-cell sets (Method 5)** |
| MzTolerances | (4.5,5) | (4.5,5) |
| RetentionTimeTolerances | (0.01,2) | (0.01,2) |
| SigmaScaleFactorRt | 3 | 3 |
| PeptideHistoryLength | 2 | 2 |
| MinUsedCorrectionPeptides | 15 | 15 |
| IntensityPeakRatioThreshold | 1e-5 | 1e-5 |
| PeptideDetectionIsoPeaks | 2 | 2 |
| IsotopeTolerance | 9 | 9 |
| Ms2DetectionNeeded | False | False |
| Ms2ExcludeDetectedPeptides | False | False |
| Ms2MinNormIntensity | 0.1 | 01 |
| Ms2MzTolerance | 20 | 20 |
| **TargetedMs2** | **Scout sample (Method 4)** | **Single-cell sets (Method 5)** |
| BatMode | False | False |
| AutoPriority | True | True |
| DefaultPriority | 0 | 0 |
| MaxNumOfScans | 1 | 1 |
| WindowAndOffsetInDalton | False | False |
| ScanDataAsProfile | False | False |
| WindowSize | 0.5 | 0.5 |
| MzOffset | 0 | 0 |
| LowerMzBound | 100 | 100 |
| CollisionEnergy | 33 | 33 |
| LifeTime | 2,400ms | 2,100ms |
| Resolution | 140,000 | 140,000 |
| MaxIT | 500ms | 500ms |
| AgcTarget | 1,000,000 | 1,000,000 |
| PositiveMode | True | True |

**Supplemental Table S13.** Prioritized acquisition parameters for mPOP BMDM single-cell samples: This table contains the MaxQuant.Live parameters used in experiments associated with Figure 1. The experiment-type-to-method mapping for these methods can be found in the corresponding table (Supporting Table S9).

|  |  |
| --- | --- |
| **Global settings: Survey scan** | **mPOP Single-cell sets (Method 6)** |
| ScanDataAsProfile | True |
| PositiveMode | True |
| MaxIT | 100ms |
| Resolution | 70,000 |
| AgcTarget | 1,000,000 |
| MzRange | (450,1258) |
| BoxCarScans | 0 |
| **Global settings: TopN** | **mPOP Single-cell sets (Method 6)** |
| NumOfMS2Scans | 0 |
| **RealtimeCorrection** | **mPOP Single-cell sets (Method 6)** |
| MzTolerances | (4.5,5) |
| RetentionTimeTolerances | (0.01,1.5) |
| SigmaScaleFactorRt | 3 |
| PeptideHistoryLength | 2 |
| MinUsedCorrectionPeptides | 15 |
| IntensityPeakRatioThreshold | 1e-5 |
| PeptideDetectionIsoPeaks | 2 |
| IsotopeTolerance | 9 |
| Ms2DetectionNeeded | False |
| Ms2ExcludeDetectedPeptides | False |
| Ms2MinNormIntensity | 0.1 |
| Ms2MzTolerance | 20 |
| **TargetedMs2** | **mPOP Single-cell sets (Method 6)** |
| BatMode | False |
| AutoPriority | True |
| DefaultPriority | 0 |
| MaxNumOfScans | 1 |
| WindowAndOffsetInDalton | False |
| ScanDataAsProfile | NA |
| WindowSize | 0.5 |
| MzOffset | 0 |
| LowerMzBound | 100 |
| CollisionEnergy | 33 |
| LifeTime | 2,400ms |
| Resolution | 140,000 |
| MaxIT | 600ms |
| AgcTarget | 1,000,000 |
| PositiveMode | True |

